# Activity-based profiling of primary brain cells identifies covalent allosteric modulators of HCN channels

**DOI:** 10.64898/2026.08.25.747126

**Authors:** Elva Ye, Alberto Russo, Roberta Castelli, Gustav T. Westlake, Xuan Jiang, Dillon A. Spiro, Sara Quejido, Cassandra L. Henry, Jacqueline L. Blankman, Gabriel M. Simon, Bruno Melillo, Bina Santoro, Anna Moroni, Benjamin F. Cravatt

**Affiliations:** Department of Chemistry, The Scripps Research Institute, La Jolla, CA, USA; Department of Biosciences, University of Milan, Milan, Italy; Lundbeck La Jolla Research Center Inc, San Diego, CA, USA; Vividion Therapeutics, 5820 Nancy Ridge Drive, San Diego, CA 92121, USA; Department of Neuroscience, Zuckerman Institute, Columbia University, New York, NY, USA

**Keywords:** Activity-based protein profiling, chemical proteomics, brain cells, covalent ligands, cysteine, stereoprobes, HCN channels, cyclic nucleotide signaling

## Abstract

Chemical proteomics can provide global portraits of small molecule-protein interactions in native biological systems. Such ligandability maps have, however, been mostly restricted to readily accessible cell lines and primary immune cells. Here, we describe an activity-based protein profiling (ABPP) strategy for mapping the covalent ligandability of primary brain cells isolated from mice. By investigating sets of stereochemically defined electrophilic small molecules (stereoprobes), we identify liganding events for diverse brain cell proteins, including many with nervous system-enriched expression. In this category were multiple hyperpolarization-activated cyclic nucleotide-gated (HCN) ion channels, which we show are covalently liganded by tryptoline acrylamide stereoprobes at a conserved cysteine in their cyclic nucleotide-binding domain. The stereoprobes were found to block cAMP-dependent shifts in voltage dependence while sparing basal activity of HCN channels. We thus describe an advanced ABPP platform for identifying ligands targeting nervous system-enriched proteins, including chemical probes that modulate HCN channel function in cells.

**Highlights:**

- Adapted ABPP for mapping covalent ligandability of primary mouse brain cells
- Identified stereoprobe ligands for diverse nervous system-enriched proteins
- Stereoprobes target a conserved allosteric cysteine in HCN channels
- Stereoprobes block cAMP modulation of HCN channels while sparing basal activity

## INTRODUCTION

Advances in genetic sequencing and perturbation technologies have increased our molecular understanding of human disease.^1,2^ Converting this knowledge into new medicines requires the discovery of small- or large-molecule agents that can perturb the functions of disease-relevant proteins. Many human proteins, however, still lack chemical probes, and some protein classes have even been considered traditionally undruggable.^3,4^

Several innovative methods have been introduced to assay proteins for interactions with small molecules, including, for instance, fragment-based ligand discovery, DNA-encoded libraries, and chemical proteomics.^5–10^ Among these approaches, chemical proteomic methods such as activity-based protein profiling (ABPP) can map small molecule-protein interactions in living cells and thus account for the myriad dynamic post-translational mechanisms that regulate protein structure and function.^4,5^ When combined with focused libraries of electrophilic small molecules, ABPP has led to the discovery of covalent ligands for a wide range of proteins, including enzymes, adaptors, RNA-binding proteins, and transcription factors.^11–15^

To date, the small molecule-protein interaction (or ligandability) maps generated by ABPP mostly originate from readily accessible cell types, such human cancer cell lines or primary peripheral immune cell populations. Many disease-relevant proteins, however, exhibit tissue-restricted expression and may be absent from immortalized cell lines. Proteins may also be subject to tissue-specific forms of post-translational regulation.^16–19^ These factors underscore the importance of expanding the cell and tissue types amenable to chemical proteomic analysis, while also preserving, to the greatest possible extent, the integrity of their biological states.

Previous ABPP studies, in particular those targeting the serine hydrolase class of enzymes^20,21^ have evaluated the ligandability of mouse brain lysates,^22^ leading to the discovery of covalent inhibitors for enzymes that regulate key lipid signaling pathways in the CNS (e.g., endocannabinoid^23–28^ and lysophospholipid^29,30^ lipases). Here, we sought to establish a protocol for performing ABPP in intact primary cells from the mouse brain and apply this method to map protein interactions for sets of cysteine-targeted, stereochemically defined electrophilic small molecules (or ‘stereoprobes’). We show that cell suspensions of gently disassociated adult mice brains (brainocytes) are amenable to ABPP using both SDS-PAGE (gel-ABPP) and quantitative tandem mass tagging (TMT) mass spectrometry (MS) readouts (protein-^13^ and cysteine-^13,31–33^ directed ABPP). Comparison of the stereoprobe ligandability maps of brainocytes versus whole brain lysates identified several proteins that show preferential liganding *in cellulo* versus *in vitro*. The brainocyte ligandability maps further contained many CNS-enriched proteins that were absent or underrepresented in previous ABPP studies of peripheral cell types and lines. We verify the stereoselective and site-specific liganding of multiple CNS-enriched proteins, leading to the discovery of tryptoline acrylamide stereoprobes that: i) engage the DPYSL2 adaptor in a complexoform-restricted manner; and ii) block cAMP-induced modulation of hyperpolarization-activated cyclic nucleotide-gated (HCN) channels. Our findings both establish an ABPP protocol for globally mapping covalent small molecule-protein interactions in intact brain cell populations and demonstrate the utility of this method for identifying ligands that perturb the functions of proteins with important roles in CNS physiology and disease.

## RESULTS

### An ABPP platform for analyzing intact mouse brain cells

In considering ways to generate global ligandability maps of CNS cell types, we felt it was important to establish an ABPP protocol that could be scaled for the MS-based proteomic analysis (e.g., TMT multiplexed^34^ experiments) of many (10s-100s of) electrophilic compounds. We accordingly favored a method that could leverage adult mouse brain tissue as opposed to primary brain cell (e.g., neuronal) cultures generated from embryonic sources. To evaluate brain cell proteomes in their native states, we initially tested two preparations from adult C57BL/6 mice: 1) brain tissue lysates generated by Dounce homogenization and probe sonication; and 2) acutely dissociated, intact brain cells (“brainocytes”) gently homogenized by a combination of enzymatic and heated mechanical methods using the MACS Octo-dissociator (**Figure 1A**). Flow cytometry analysis showed a consistently high viability (∼90%) of the brainocyte preparations (**Supplementary Dataset S1**). We then compared these preparations by gel- and MS (protein-directed)-ABPP using an initial set of alkynylated tryptoline acrylamide stereoprobes^13^ (WX-01-05/06/07/08 (5 µM, 1 h); **Figure 1B** and **Figure S1A**)).

**Figure 1.**
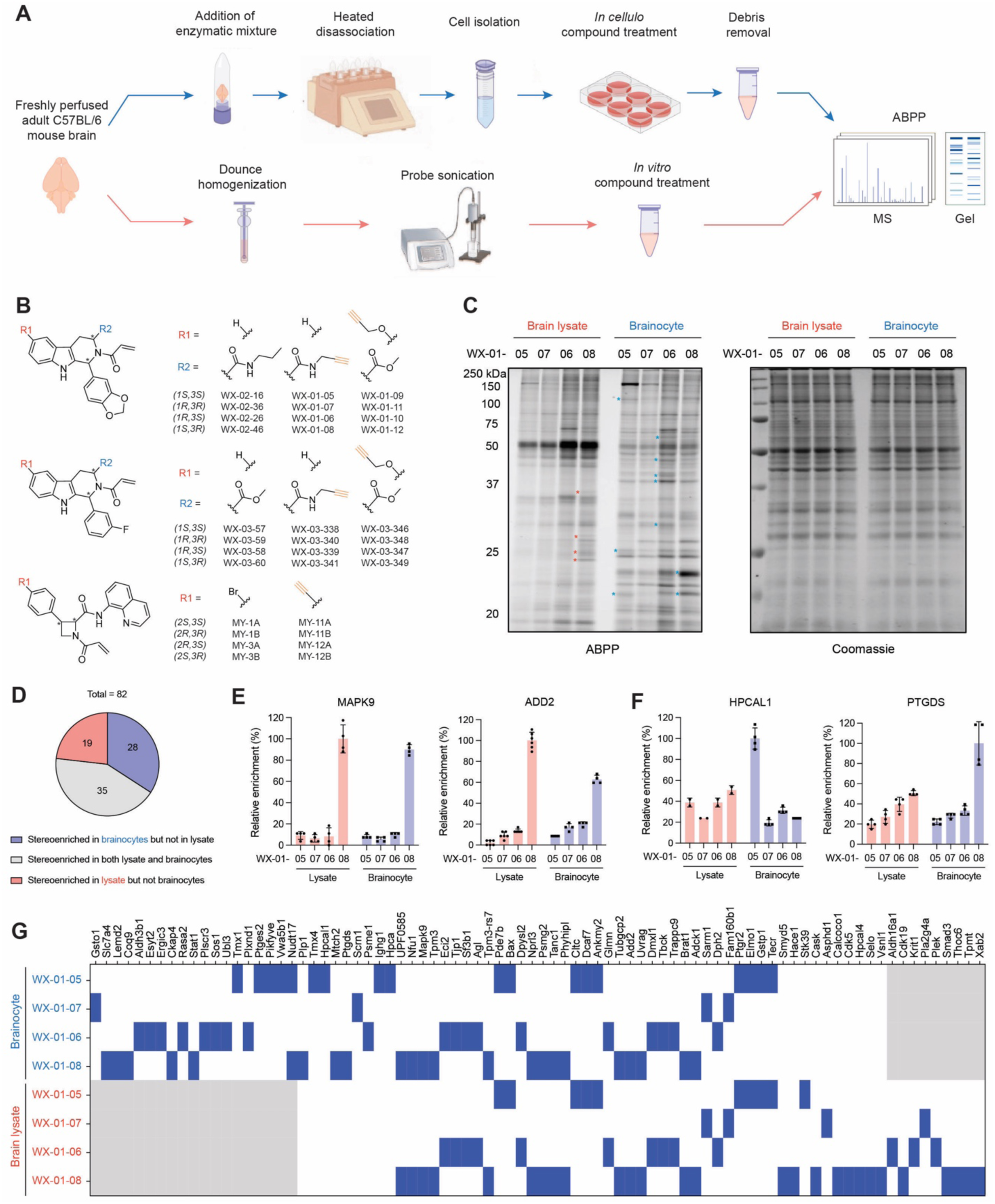
An ABPP protocol for analyzing intact mouse brain cells. **(A)** Workflow for *in cellulo* ABPP of acutely dissociated intact brain cells (“brainocytes”). Blue arrows: freshly perfused brain tissue from adult C57BL/6 mice was gently homogenized by enzymatic and heated mechanical methods using the MACS Octodissociator, after which brain cells were isolated by 70 μm cell strainers, resuspended with DMEM media, and treated with compounds (stereoprobes) in six-well dishes (∼20 million cells per well) following established protein- or cysteine-directed ABPP protocols.^13^ Debris from the brainocyte preparation was removed after stereoprobe treatment by low-speed centrifugation (3000 g, 5 min) prior to gel- or MS-analysis. Red arrows: alternative freshly perfused brain tissue samples were converted into tissue lysates by sequential Dounce homogenization and probe sonication to generate material for *in vitro* ABPP experiments. **(B)** Chemical structures of parent and stereochemically matched alkynylated tryptoline and azetidine acrylamide stereoprobes used in this study. **(C)** Gel-ABPP data for brain lysates and brainocytes treated with alkynylated tryptoline acrylamides WX-01-05/06/07/08 (5 μM, 1 h). Stereoprobe-reactive proteins were visualized by copper-catalyzed azide-alkyne cycloaddition (CuAAC or click^123,124^) conjugation to an azide-rhodamine reporter group, SDS-PAGE separation, and in-gel fluorescence scanning.^125^ Left and right gels show rhodamine-ABPP data and Coomassie blue staining, respectively. For ABPP data, red and blue asterisks mark proteins showing stereoselective reactivity with the tryptoline acrylamides in brain lysates and brainocytes, respectively. Data are from a single experiment representative of at least two independent experiments. **(D)** Pie chart showing the number of enantioselectively enriched proteins in brainocytes, brain lysates, or both, from protein-directed ABPP experiments performed with alkynes WX-01-05/06/07/08 (5 μM, 1 h). **(E)** Protein-directed ABPP data showing representative proteins enantioselectively enriched by the indicated stereoprobes in both brainocytes and brain lysates. Data are average values ± SD from four to six independent experiments normalized to the sample with the maximum mean value. **(F)** Protein-directed ABPP data showing representative proteins enantioselectively enriched by the indicated stereoprobes in brainocytes, but not brain lysates (HPCAL1: WX-01-05; PTGDS: WX-01-08). Data are average values ± SD from four to six independent experiments normalized to the sample with the maximum mean value. **(G)** Heatmap presentation of proteins stereoenriched in brainocytes and/or brain lysates from protein-directed ABPP experiments performed as described in **(D)**. Gray boxes indicate that the protein was not quantified in that sample.

Initial gel-ABPP experiments revealed clear differences in the proteome-wide reactivity profiles of brain lysates vs brainocytes, including several proteins exhibiting preferential stereoselective engagement in one of the two preparations (asterisks, **Figure 1C**, left). In contrast, brain lysates and brainocytes showed similar protein signals by Coomassie blue staining (**Figure 1C**, right), indicating the stereoprobe reactivity differences were not secondary consequences of alterations in protein abundance.

Protein-directed ABPP experiments confirmed striking differences in the stereoprobe reactivity of brain lysates and brainocytes. Using a filter of 2.5-fold enantioselective enrichment by alkyne stereoprobes, we identified a total of 82 enantioenriched proteins, of which 35 were shared by both brain lysates and brainocytes and 19 and 28 were unique brain lysates and brainocytes, respectively (**Figure 1D** and **Dataset S2**). Examples of proteins showing stereoprobe enrichment in both brain lysates and brainocytes versus enrichment only in brainocytes are shown in **Figure 1E** and **F**, respectively. Curious whether the brainocyte-restricted events reflected authentic examples of stereoprobe interactions that required the intact environment of living brain cells, we performed pilot protein-directed ABPP experiments in acute mouse brain slices (**Figure S1C**). These experiments revealed brainocyte-restricted tryptoline acrylamide-protein interactions were generally recapitulated in acute brain slices (**Figure S1D** and **Dataset S2**).

Based on our initial ABPP data, we concluded that the brainocyte preparation provided a suitable and scalable approximation of intact brain cells for global ligandability mapping.

### Electrophilic stereoprobe ligandability maps of mouse brainocytes

Among the various classes of electrophilic stereoprobes described to date,^13,15,31,35–39^ we selected the azetidine and tryptoline acrylamides (**Figure 1B**) for protein- and cysteine-directed ABPP of mouse brainocytes, as these stereoprobes have been found to ligand broad and largely non-overlapping collections of proteins in past studies of human cancer^13,35^ and immune^31,35^ cell types. Protein-directed ABPP experiments were performed generally as described in the previous section, but we included additional samples that were pre-treated with parent (non-alkyne) stereoprobes (20 µM, 2 h) to determine the stoichiometry of stereoprobe engagement of proteins. Proteins were classified as stereoselectively liganded if they displayed greater than 2.5-fold enantioselective enrichment by an alkyne stereoprobe and greater than 33% competitive blockade of this enrichment by the corresponding parent competitor stereoprobe. Similarly, cysteine-directed ABPP experiments were performed using a broad-spectrum iodoacetamide-desthiobiotin (IA-DTB) probe as described previously,^13,31^ where cysteines were considered stereoselectively liganded if they showed >33% decrease in IA-DTB reactivity with a given stereoprobe, and this decrease was at least 2.5-fold greater in magnitude than the effect of the enantiomeric stereoprobe. These criteria for stereoprobe liganding events were more relaxed than those used previously for ABPP experiments performed in cell lines (e.g. 33% versus 50% inferred engagement by parent stereoprobes), which we felt was justified due to the apparent lower overall uptake of stereoprobes in the brainocyte preparations compared to standard cultured cell lines (as assessed by gel-ABPP; **Figure S2A**). We performed two independent protein-directed ABPP experiments and three independent cysteine-directed ABPP experiments for each set of stereoprobes, with each of these multiplexed experiments containing two replicates, resulting in four and six independent replicates, respectively.

The ABPP experiments, in aggregate, quantified >11000 cysteines and >5600 proteins, from which a total of 114 stereoprobe-liganded proteins were identified (**Figure 2A** and **Dataset S2**). As visualized in quadrant plots, the liganded proteins distributed across all four stereoisomeric configurations of the stereoprobes (**Figures 2B-D** and **S2B, C**). To better understand the fraction of stereoprobe-liganded proteins with potentially specific roles in the nervous system, we next established a list of CNS-enriched proteins by aggregating information from RNA sequencing data in public repositories (e.g., bioGPS^40^ and GTEx^41^; **Figure S2D** and **Dataset S2**). This analysis revealed that ∼25% of the liganded proteins were enriched in expression in the CNS (**Figure 2E** and **Table 1**). The CNS-enriched liganded proteins originated from diverse structural and functional classes (**Figure 2F**) and were distributed in relative expression across several major brain cell types (**Figure 2G**). Interestingly, the majority of the CNS-enriched liganded proteins were not identified in previous ABPP studies of primary immune and cancer cell lines (**Table 1**), indicating that the brainocyte preparation provided improved access to proteins with restricted expression in the nervous system.

**Figure 2.**
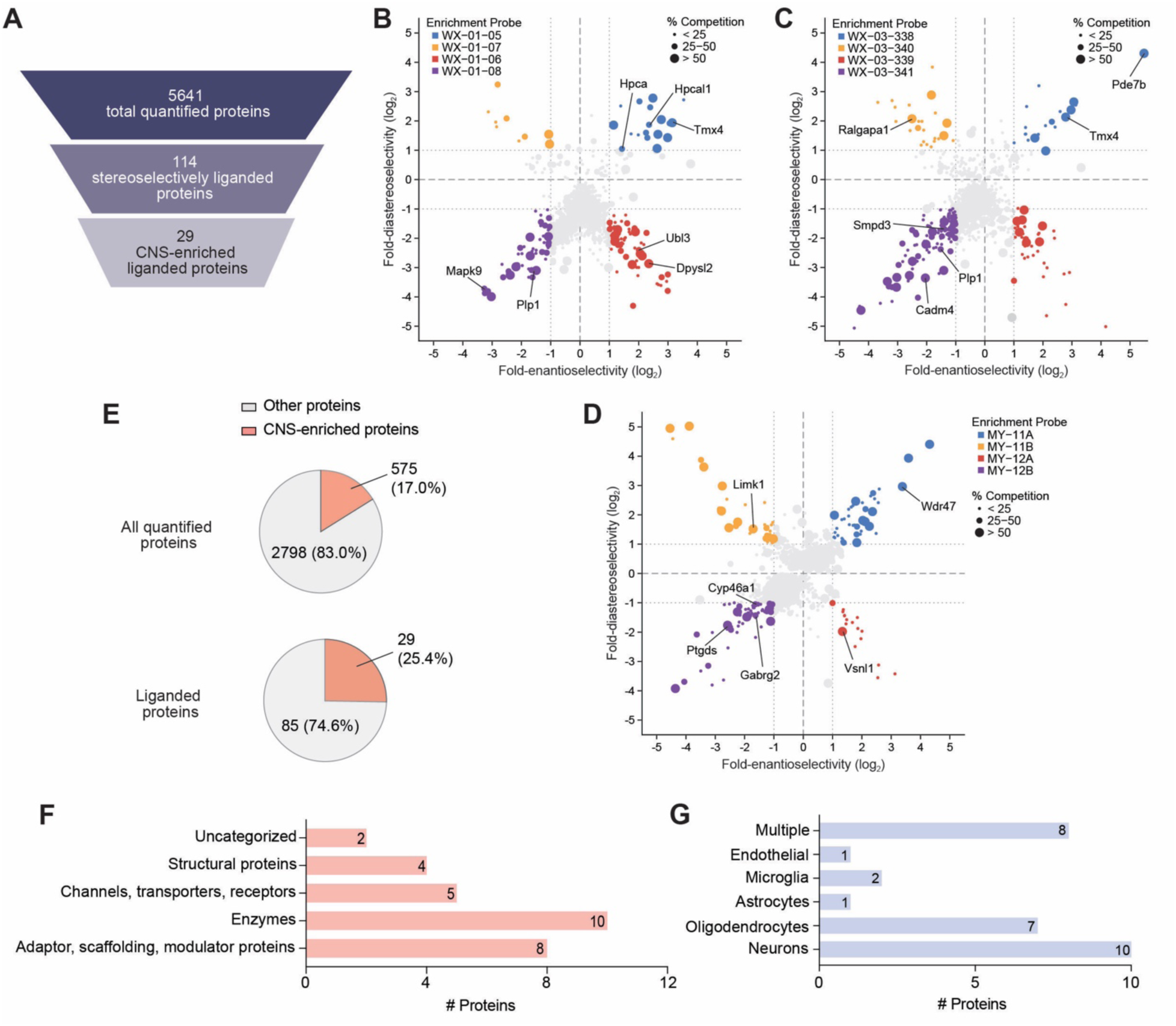
Stereoprobe ligandability maps of mouse brainocytes. **(A)** Total numbers of proteins, and the total liganded and CNS-enriched liganded subsets of proteins, quantified by cysteine-directed ABPP and protein-directed ABPP in brainocytes. Liganded proteins were defined as those showing >2.5x enantioselective enrichment by alkyne stereoprobes and >33% competition by parent stereoprobes in either protein- or cysteine-directed ABPP experiments. **(B-D)** Representative quadrant plots showing protein-directed ABPP data for brainocytes treated with three stereoprobe sets of distinct chemotypes. Colors highlight the stereoselectively enriched proteins for each stereoconfiguration of alkyne stereoprobe, where each protein is displayed once based on its highest enrichment value, and the size of the dots highlight the degree of competitive blockade of enrichment by the corresponding parent stereoprobe. Enantioselectivity (x-axis) is the ratio of enrichment for one stereoisomer versus its enantiomer, and diastereoselectivity (y-axis) is the ratio of enrichment of one stereoisomer versus the average of its two diastereomers. Representative liganded proteins are labeled. For a more complete list of liganded proteins, see **Table S2**. **(E)** Pie charts showing the percentage of CNS-enriched proteins for all quantified proteins and the subset of stereoprobe-liganded proteins in mouse brainocytes. All quantified proteins (top) were determined by unenriched quantitative proteomics of brainocytes, and the liganded proteins (bottom) were determined from cysteine- and protein-directed ABPP experiments, as summarized in panel **(A)**. For a complete list of CNS-enriched liganded proteins, see **Table 1**. **(F)** Functional class distribution of CNS-enriched liganded proteins assigned as described previously using GO (Panther), KEGG BRITE, and UniProt annotations.^13^ **(G)** Distribution of brain cell type expression for CNS-enriched liganded proteins analyzed using mouse Brain RNA-Seq data^126^, where Z-scores were calculated for each brain cell type, and proteins with a brain cell type Z score of >1.5 were classified as enriched for the specified brain cell type. Otherwise “multiple” was used to denote the brain cell type enrichment for a liganded protein.

**Table 1.** List of CNS-enriched stereoprobe-liganded proteins as determined by cysteine- and protein-directed ABPP in brainocytes. Information on the ligandability of proteins and cysteines in cell lines was acquired from previous publications. ^13,37,38^

| Protein Name | Protein Accession | Liganded Cysteine | Functional Class | Brain cell type enrichment | Active competitor Stereoprobes | Liganding previously observed in immune cells or cell lines? |
| --- | --- | --- | --- | --- | --- | --- |
| CADM4 | Q8R464 |  | Structural proteins | Oligodendrocyte | WX-02-46, WX-03-60 | N |
| CYP46A1 | Q9WVK8 |  | Enzymes | Non-specific | MY-3B | N |
| DPYSL2 | O08553 | 504 | Structural proteins | Neuron | WX-02-26 | N |
| FLOT2 | Q60634 |  | Adaptor, scaffolding, modulator proteins | Non-specific | WX-03-60 | N |
| GABRG2 | P22723 |  | Channels, transporters, receptors | Neuron | MY-3B | N |
| HCN1 | O88704 | 531 | Channels, transporters, receptors | Neuron | WX-02-46 | N |
| HPCA | P84075 |  | Adaptor, scaffolding, modulator proteins | Neuron | WX-02-16 | N |
| HPCAL1 | P62748 |  | Adaptor, scaffolding, modulator proteins | Non-specific | WX-02-16 | Y |
| KCNK1 | O08581 |  | Channels, transporters, receptors | Oligodendrocyte | WX-02-46 | N |
| LANCL1 | O89112 | 108 | Enzymes | Neuron | WX-02-26 | N |
| LIMK1 | P53668 | 349 | Enzymes | Microglia | MY-1B | Y |
| MAP6D1 | Q14BB9 |  | Structural proteins | Oligodendrocyte | MY-3B | N |
| MAPK9 | Q9WTU6 |  | Enzymes | Neuron | WX-02-46 | N |
| PDE7B | Q9QXQ1 | 136 | Enzymes | Endothelial | WX-03-57 | N |
| PHYHIP | Q8K0S0 |  | Uncategorized | Non-specific | WX-03-60 | N |
| PLLP | Q9DCU2 |  | Adaptor, scaffolding, modulator proteins | Oligodendrocyte | WX-02-46 | N |
| PLP1 | P60202 | 6 | Structural proteins | Oligodendrocyte | WX-02-46, WX-03-60 | N |
| PTGDS | O09114 | 65 | Enzymes | Oligodendrocyte | WX-03-60, MY-3B | N |
| RALGAPA1 | Q6GYP7 |  | Enzymes | Non-specific | WX-03-58 | Y |
| SCRN1 | Q9CZC8 |  | Enzymes | Non-specific | WX-02-36 | N |
| SIPA1L1 | Q8C0T5 | 1037 | Adaptor,<br>scaffolding,<br>modulator<br>proteins | Microglia | MY-1B | N |
| SLC32A1 | O35633 |  | Channels,<br>transporters,<br>receptors | Neuron | WX-03-60 | N |
| SLC6A1 | P31648 |  | Channels,<br>transporters,<br>receptors | Astrocyte | WX-02-46 | N |
| SMPD3 | Q9JJY3 |  | Enzymes | Neuron | WX-03-60 | Y |
| SRCIN1 | B1AQX9 |  | Adaptor,<br>scaffolding,<br>modulator<br>proteins | Non-specific | WX-03-60 | N |
| TMX4 | Q8C0L0 |  | Enzymes | Neuron | WX-02-16, WX-03-57 | Y |
| UBL3 | Q9Z2M6 |  | Adaptor,<br>scaffolding,<br>modulator<br>proteins | Oligodendrocyte | WX-02-36 | N |
| VSNL1 | P62761 | 187 | Adaptor,<br>scaffolding,<br>modulator<br>proteins | Neuron | MY-3A | N |
| WDR47 | Q8CGF6 |  | Uncategorized | Non-specific | MY-1A | Y |

### Characterization of stereoprobe-protein interactions

We next selected four CNS-enriched liganded proteins for further characterization—PLP1, PDE7B, DPYSL2, and HCN1. These stereoprobe targets were chosen because they: i) belong to different structural and functional classes of proteins (scaffolding (PLP1), enzyme (PDE7B), adaptor (DPYSL2), channel (HCN1)); ii) had not been identified in previous ABPP studies of other non-CNS cell types; and iii) are implicated in neurological and neurodegenerative disorders. We further evaluated the human ortholog of each stereoprobe target so that we could determine whether liganding was a shared feature across species.

PLP1 is a multi-pass transmembrane protein that serves as a principal component of myelin and is crucial for oligodendrocyte development and axonal survival.^42,43^ Mutations in the *PLP1* gene cause the hypomyelinating leukodystrophy Pelizaeus-Merzbacher disease in humans.^44^ In our protein-directed ABPP experiments, PLP1 was stereoselectively engaged by the (1*S*, 3*R*) pair of WX-01-12/WX-02-46 pair of tryptoline acrylamides (**Figure 3A** and **Dataset S2**). Cysteine-directed ABPP of mouse brainocytes revealed that WX-02-46 produced a stereoselective decrease in the IA-DTB reactivity of the tryptic peptide containing C6 and C7 of PLP1, while other quantified cysteines in the protein were generally unaffected (**Figure 3B** and **Dataset S2**). Since cysteine-directed ABPP cannot distinguish the liganding of cysteines that reside on the same tryptic peptide, we performed site-directed mutagenesis, which revealed that only a C6A-PLP1 mutant, but not a C7A-PLP1 mutant, lost reactivity with WX-01-12 (**Figure 3C**). We interpret these results to indicate that C6 is the stereoprobe-liganded cysteine in PLP1. Interestingly, C6 is also a site for palmitoylation in PLP1, and this post-translational modification is important for targeting the protein to nascent myelin membranes (**Figure 3D**),^45–47^ suggesting that covalent ligands targeting C6 may have the potential to perturb PLP1 trafficking and function.

**Figure 3.**
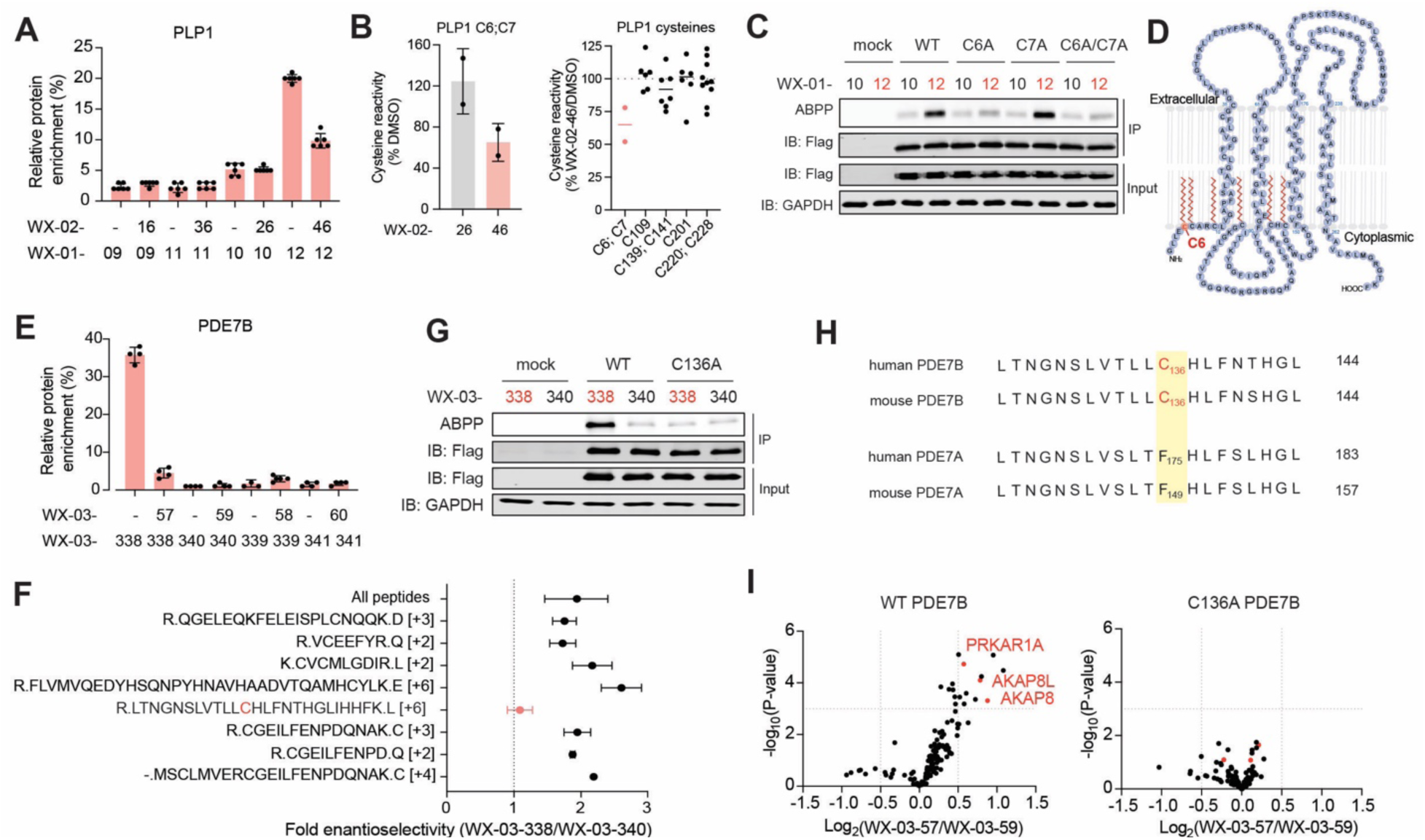
Characterization of representative CNS-enriched liganded proteins. **(A)** Protein-directed ABPP data showing stereoselective enrichment of PLP1 by WX-01-12 (5 μM, 1 h) and blockade of this enrichment by pre-treatment with WX-02-46 (20 μM, 2 h) in brainocytes. Data are average values ± SD from four independent experiments. **(B)** Left, cysteine-directed ABPP data showing that a tryptic peptide containing C6 and C7 of PLP1 is stereoselectively liganded by WX-02-46 (20 μM, 3 h) in brainocytes. Right, reactivity profiles of all quantified cysteine residues in PLP1 in WX-02-46-treated brainocytes as determined by cysteine-directed ABPP. Data are average values ± SD from six independent experiments. **(C)** Gel-ABPP data showing stereoselective engagement of recombinant FLAG-tagged human WT-PLP1 and a C7A-PLP1 mutant, but not C6A-PLP1 or C6A/C7A-PLP1 mutants by WX-01-12 (5 μM, 1 h) in HEK293T cells. Gel-ABPP was performed on protein samples after an anti-FLAG immunoprecipitation (IP) step as described in the Methods. Data are from a single experiment representative of at least two independent experiments. **(D)** Topology map of PLP1 showing C6, highlighted in red as a known palmitoylation site.^45^ **(E)** Protein-directed ABPP data showing stereoselective enrichment of PDE7B by WX-03-338 (5 μM, 1 h) and blockade of this enrichment by WX-03-57 (20 μM, 2 h) in brainocytes. Data are average values ± SD from four independent experiments. **(F)** Stereoselective enrichment profiles of cysteine-containing and all tryptic peptides from PDE7B by WX-03-338 (5 μM, 1 h) quantified from protein-directed ABPP experiments performed in WT-PDE7B-expressing in HEK293T cells. The corrupted enrichment of the tryptic peptide containing C136 is highlighted in red. Data are average values ± SD from two independent experiments. **(G)** Gel-ABPP data showing stereoselective engagement of recombinant FLAG-tagged human WT-PDE7B, but not a C136A-PDE7B mutant by WX-03-338 (5 μM, 1 h) in HEK293T cells. Data are from a single experiment representative of at least two independent experiments. **(H)** Sequence ment of the indicated amino acid region of mouse and human PDE7B and PDE7A (numbers on right refer to the C-terminal residue number for each sequence), showing that PDE7B_C136 is not conserved in paralog PDE7A. **(I)** Volcano plots comparing co-enriched proteins from anti-FLAG IP-MS experiments performed with SH-SY5Y cells expressing FLAG-tagged WT- or C136A-PDE7B treated with WX-03-57 or WX-03-59 (20 μM, 3h). Data are average values from six independent experiments. *p* values were calculated using two-tailed Welch’s *t*-test.

PDE7B is a CNS-enriched phosphodiesterase involved in terminating the signaling function of intracellular cyclic nucleotide second messengers (cAMP, cGMP)^48^. The genetic or pharmacological perturbation of PDE7 enzymes, including PDE7B, has been shown to exhibit neuroprotective effects in animal models.^49–51^ In our protein-directed ABPP experiments, PDE7B was stereoselectively engaged by the (1*S*, 3*S*) WX-03-338/WX-03-57 pair of tryptoline acrylamides (**Figure 3E** and **Dataset S2**). Cysteine-directed ABPP experiments did not identify a cysteine in PDE7B that was perturbed by WX-03-57; however, protein-directed ABPP experiments performed in PDE7B-expressing HEK293T cells revealed a corrupted stereoenrichment for the tryptic peptide containing C136, a profile that we have found to be a hallmark of stereoprobe-liganded cysteines in past studies^13^ (**Figure 3F**). We confirmed by gel-ABPP that WX-03-338 did not react with a C136A-PDE7B mutant expressed in HEK293T cells (**Figure 3G**). Similarly, the C136-containing tryptic peptide was not stereoselectively enriched in protein-directed ABPP experiments performed in PDE7B-stably expressing SH-SY5Y cells (**Figure S3B, C**) and was the only non-stereoenriched cysteine that abolished WX-03-338 labeling when mutated to alanine in gel-ABPP experiments (**Figures S3D**). Taken together, these data indicate that C136 is the likely site of stereoprobe engagement in PDE7B. Of note, C136 is not conserved in the paralogous protein PDE7A (**Figure 3H**), and we did not observe PDE7A as a stereoprobe target by protein-directed ABPP (**Dataset S2**), suggesting that WX-03-338/WX-03-57 are isoform-restricted ligands for PDE7B.

An AlphaFold2 model of PDE7B indicated that C136 is located at a non-orthosteric site distal to the active site of the enzyme (**Figure S3E**). We further found that WX-03-57 did not inhibit PDE7B-dependent hydrolysis of cAMP (**Figure S3F**). Immunoprecipitation-mass spectrometry (IP-MS) experiments performed in SH-SY5Y cells stably expressing WT- or C136A-PDE7B, on the other hand, revealed that WX-03-57 stereoselectively and site-specifically enhanced WT-PDE7B interactions with several proteins, including PRKAR1A, AKAP8L, and AKAP8 (**Figure 3I** and **S3G**). The PKA-AKAP-PDE complex or microdomain is thought to serve as a mechanism for cells to spatially regulate cAMP signaling by compartmentalizing the levels of this second messenger at specific subcellular locations (**Figure S3H**).^52–54^ Our data thus suggest that covalent ligands targeting C136 in PDE7B may offer a way to further tune the structure and function of PKA-AKAP-PDE complexes in brain cells.

DPYSL2 is a member of a family of five cytosolic phosphoproteins (DPYSL1-5) that share high sequence identity (50-75%) and are also termed collapsin response mediator proteins (CRMP1-5) in recognition of their roles in neuronal development and polarity, growth cone collapse, cell migration, and cytoskeleton formation.^55–57^ The DPYSL proteins possess a dihydropyriminidase fold, but lack catalytic residues and enzymatic activity,^58,59^ and instead form homo- and hetero-tetrameric structures that serve as adaptors connecting kinase-mediated signaling pathways to cytoskeletal dynamics.^60^ Human genetic data implicate DPYSL2 in neurodevelopmental and neuropsychiatric diseases, including non-synonymous mutations that confer loss-of-function effects on the protein.^56,60^

In our protein-directed ABPP experiments, DPYSL2 showed strong stereoselective engagement by the (1*R*, 3*S*) WX-01-06/WX-02-26 pair of tryptoline acrylamides (**Figure 4A** and **Dataset S2**). Other DPYSL proteins did not exhibit evidence of liganding with the exception of DPYSL5, which displayed modest stereoselective engagement by WX-01-06/WX-02-26 (**Figure S4A** and **Dataset S2**). Notably, DPYSL2 has been previously quantified in Ramos cells but showed no stereoselective enrichment (**Figure S4B**).^13^ We also noticed that the tryptic peptide containing C504 of DPYSL2 (aa 497-511) consistently showed corrupted stereoenrichment in protein-directed ABPP experiments (**Figure 4B**); however, we did not observe a decrease in IA-DTB reactivity for DPYSL2_C504 or any of the other six cysteines in DPYSL2 in WX-02-26-treated brainocytes by cysteine-directed ABPP (**Figure 4C**).

**Figure 4.**
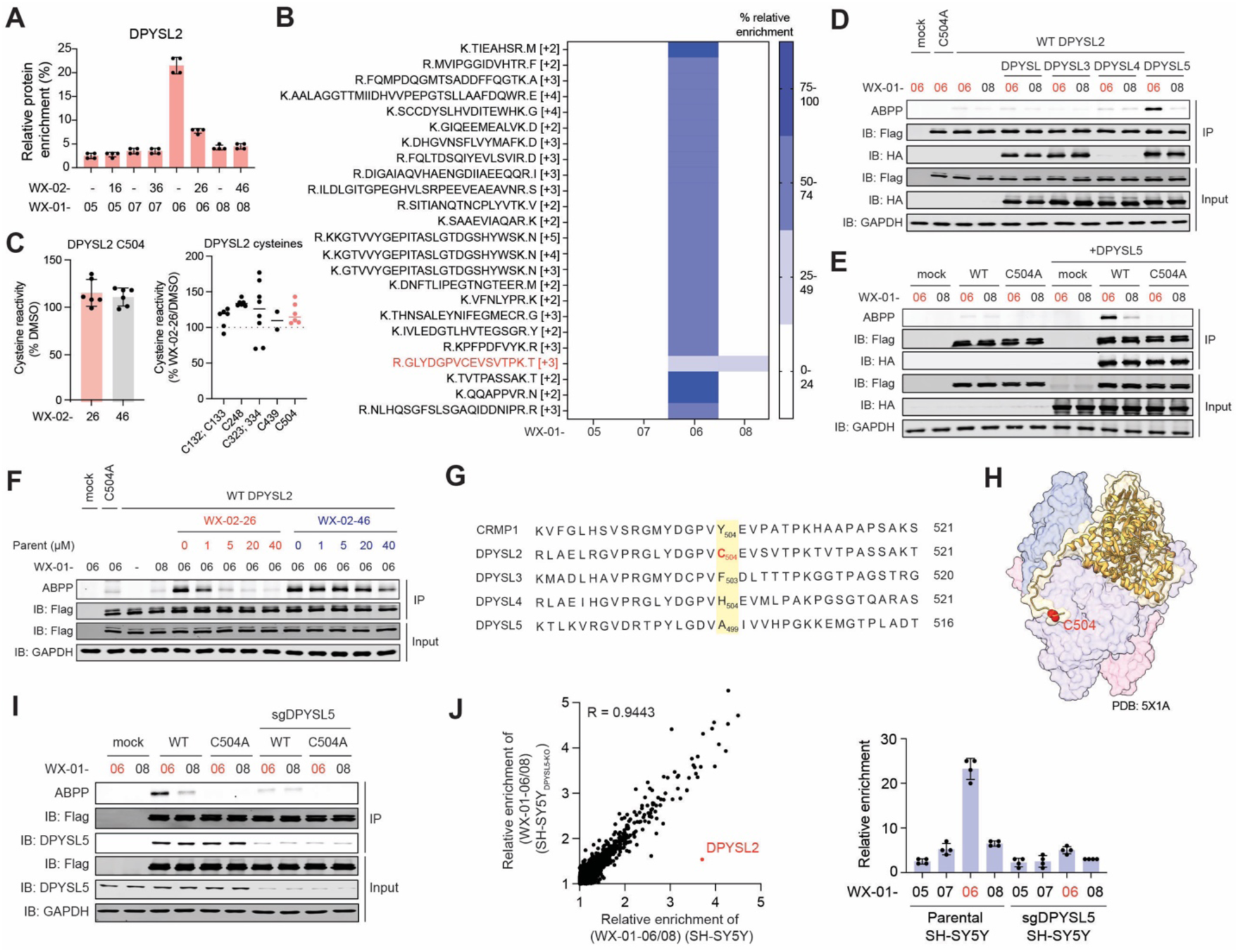
Characterization of complexoform-restricted stereoprobe ligands for DPYSL2. **(A)** Protein-directed ABPP data showing stereoselective enrichment of DPYSL2 by WX-01-06 (5 μM, 1 h) and blockade of this enrichment by pre-treatment with WX-02-26 (20 μM, 2 h) in brainocytes. Data are average values ± SD from four independent experiments. **(B)** Tryptic peptide map of DPYSL2 from protein-directed ABPP experiments showing stereoselective enrichment of all quantified DPYSL2 peptides in WX-01-06-treated brainocytes with the exception of the peptide containing C504 (red). **(C)** Cysteine-directed ABPP data for DPYSL2_C504 (left) and all quantified DPYSL2 cysteines (right) from brainocytes treated with WX-02-26 or WX-02-46 (20 μM, 3 h) (left). Cysteines separated by a semicolon (;) are located on the same tryptic peptide. Data are average values ± SD from six independent experiments. **(D)** Gel-ABPP data showing stereoselective engagement of FLAG-tagged human DPYSL2 by WX-01-06 (5 µM, 1 h) in HEK293T cells co-expressing HA-tagged human DPYSL5, but not other DPYSL paralogs. Data are from a single experiment representative of at least two independent experiments. **(E)** Gel-ABPP data showing stereoselective engagement of FLAG-tagged WT, but not a C504A-DPYSL2 mutant by WX-01-06 (5 µM, 1 h) in HCT116 cells co-expressing HA-tagged DPYSL5. Data are from a single experiment representative of at least two independent experiments. **(F)** Gel-ABPP data showing concentration-dependent and stereoselective blockade of WX-01-06 (5 μM, 1 h) engagement of DPYSL2 by WX-02-26 (indicated concentrations, 2 h) in Neuro2a cells stably expressing both FLAG-tagged DPYSL2 and HA-tagged DPYSL5. Data are from a single experiment representative of at least two independent experiments. **(G)** Sequence alignment of the indicated amino acid regions of DPYSL family proteins (numbers on right refer to the C-terminal residue number for each sequence), showing that C504 (in red) is unique to DPYSL2. **(H)** Crystal structure of DPYSL2 homo-tetramer (PDB ID: 5X1A^127^). DPYSL2 monomer 1 is represented in yellow, monomers 2-4 are represented in shades of light purple. C504 (highlighted in red in monomer 1) is located on the C-terminal tail that resides at an interface between monomers. **(I)** Gel-ABPP showing stereoselective engagement of recombinant WT-, but not C504A-DPYSL2 by WX-01-06 (5 µM, 1 h) in parental or DPYSL5-disrupted (sgDPYSL5) Neuro2a cells. DPYSL5 was genetically disrupted in Neuro2a cells by CRISPR/Cas9 methods as described in the Methods. Mock cells correspond to parental Neuro2a cells without recombinant expression of DPYSL2. Data are from a single experiment representative of at least two independent experiments. **(J)** Protein-directed ABPP data showing stereoselective enrichment of endogenous DPYSL2 by WX-01-06 (5 μM, 1 h) in parental SH-SY5Y cells but not sgDPYSL5 SH-SY5Y cells. Left, scatter plot comparing the stereoselective enrichment profiles (WX-01-06/08, 5 μM, 1 h) of proteins in parental (x axis) versus sgDPYSL5 (y-axis) SH-5Y5Y cells. Right, quantification of DPYSL2 signals in protein-directed ABPP data. Data are average values from four independent experiments. The linear regression analysis was performed excluding the DPYSL2 data.

Our initial gel-ABPP experiments with recombinant human DPYSL2 expressed in HCT116 cells did not show evidence of stereoselective reactivity with WX-01-06 (**Figure S4C**). Recognizing that DPYSL proteins form both homo and hetero-oligomers, we next co-expressed DPYSL2 with other DPYSL paralogs, which revealed that co-expression specifically with DPYSL5 restored stereoselective reactivity of DPYSL2 with WX-01-06 (**Figure 4D**). Additionally, this stereoprobe reactivity profile was not observed when co-expressing a C504A-DPYSL2 mutant with DPYSL5 (**Figure 4E**). Competitive gel-ABPP experiments confirmed that pre-treatment with WX-02-26, but not the enantiomer WX-02-46, produced a concentration-dependent blockade of WX-01-06 reactivity with WT-DPYSL2 in cells co-expressing DPYSL5 (**Figure 4F**).

We interpreted the aforementioned data to indicate that WX-01-06/WX-02-26 react with C504 of DPYSL2 only when this protein forms hetero-oligomeric complex with DPYSL5. C504 is unique to DPYSL2 (**Figure 4G**), which could further explain why this protein is the only member of the DPYSL family that showed liganding in our ABPP experiments (and, by extension, the more marginal stereoprobe interactions observed for DPYSL5 may reflect indirect co-enrichment of this protein as part of hetero-oligomeric complexation with DPYSL2). We recently discovered a similar type of complexoform-restricted liganding for the pleotropic adaptor protein TRMT112 when bound to one of its methyltransferase partners METTL5, where the stereoprobe-liganded cysteine was found to be located at the TRMT112:METTL5 interface.^15^ Analogously, C504 of DPYSL2 is located at the inter-subunit interface of DPYSL2 homo-tetramers (**Figure 4H**), which could indicate that structural differences in this interface in the DPYSL2:DPYSL5 complex support WX-01-06/WX-02-26 binding and reactivity. Under this model, if only a small portion of DPYSL2 is associated with DPYSL5 in brainocytes, then the impact of WX-02-26 on bulk DPYSL2_C504 would be negligible in cysteine-directed ABPP experiments; on the other hand, in protein directed-ABPP experiments, the alkyne stereoprobe WX-01-06 would be expected to engage only the fraction of DPYLS2 in brainocytes that is interacting with DPYSL5, and this engagement would, in turn, be sensitive to disruption by pre-treatment with WX-02-26 (**Figure S4E**), thus providing a means to map the liganding of a rare complexoform of an abundant adaptor protein. Consistent with this model, we found that WX-01-06 engagement of recombinant DPYSL2 could be visualized by gel-ABPP without co-expression of DPYSL5 in the mouse neuroblastoma cell line Neuro2A, which expresses substantial amounts of endogenous DPYSL5 (**Figure 4I** and **S4F**). Conversely, CRISPR/Cas9-mediated genetic disruption of DPYSL5 in the Neuro2A cells or the human neuroblastoma cell line SH-SY5Y blocked stereoselective engagement of recombinant and endogenous DPYSL2 by WX-01-06 as determined by gel-(**Figure 4I**) or protein-directed ABPP (**Figures 4J** and **S4G**), respectively. These data suggest that tryptoline acrylamides may offer chemical tools to study the specific functions of the DPYSL2:DPYSL5 complex while sparing other paralogous DPYSL complexes in cells.

HCN1 is a member of a family of hyperpolarization-activated cyclic nucleotide-gated ion channels that regulate neuronal excitability and cardiac pacemaker activity.^61^ Our combined protein- and cysteine-directed ABPP data indicated that HCN1 was stereoselectively liganded at C531 by the (1*S*, 3*R*) WX-01-08/WX-02-46 pair of tryptoline acrylamides (**Figure 5A, B** and **Dataset S2**), with the cysteine-directed ABPP data supporting that this conserved cysteine residue was also liganded by WX-02-46 in additional HCN channels (HCN2_C584, HCN4_C662; **Figure 5B, S5A**). We confirmed by gel-ABPP the stereoselective and site-specific engagement of recombinant HCN1, HCN2, and HCN4 individually expressed in HEK293T cells by alkyne stereoprobe WX-01-08 (**Figure 5C** and **Figure S5B, C**).

**Figure 5.**
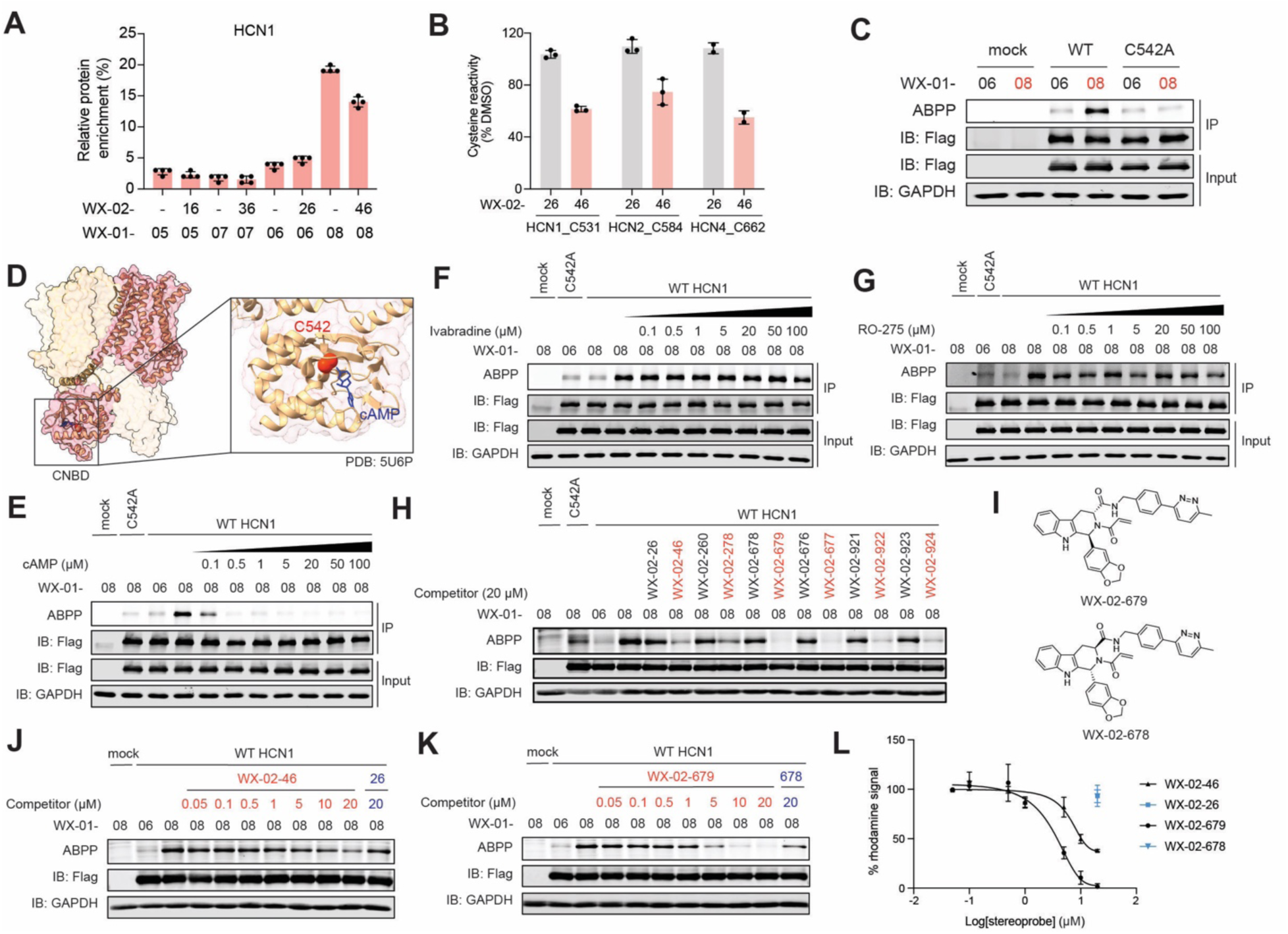
Characterization of stereoprobe ligands targeting the cyclic nucleotide binding domain of HCN channels. **(A)** Protein-directed ABPP data showing stereoselective enrichment of HCN1 by WX-01-08 (5 μM, 1 h) and blockade of this enrichment by WX-02-46 (20 μM, 2 h) in brainocytes. Data are average values ± SD from four independent experiments. **(B)** Cysteine-directed ABPP data showing stereoselective engagement of a conserved cysteine in HCN channels (HCN1_C542, HCN2_C584, and HCN4_C662) by WX-02-46 (20 μM, 3 h) in brainocytes. Data are average values ± SD from two or three independent experiments. **(C)** Gel-ABPP data showing stereoselective engagement of recombinant FLAG-tagged human WT-HCN1, but not a C542A-HCN1 mutant by WX-01-08 (5 μM, 1 h) in HEK293T cells. Data are from a single experiment representative of at least two independent experiments. **(D)** Cryo-EM structure of the HCN1 homo-tetrameric channel (PDB: 5U6P^63^). HCN1 monomer 1 is represented in pink, monomers 2-4 are represented in shades of yellow. C542 (highlighted in red in monomer 1) is located in proximity to the cyclic AMP (cAMP)-binding site in the cyclic nucleotide binding domain (CNBD). cAMP is shown in blue. **(E)** Gel-ABPP data showing concentration-dependent blockade of WX-01-08 (5 μM, 1 h) engagement of recombinant HCN1 by pre-treatment with cAMP (20 min, indicated concentrations) performed in HEK293T cell lysates. Data are from a single experiment representative of at least two independent experiments. **(F)** and **(G)** Gel-ABPP data showing that pre-treatment (1 h) with the indicated concentrations of the HCN pore blocking compounds ivabradine **(F)** and RO-275 **(G)** did not affect the reactivity of WX-01-08 (5 μM, 1 h) with recombinant HCN1 in HEK293T cells lysates. Data are from a single experiment representative of at least two independent experiments. **(H)** Gel-ABPP screen showing effects of pre-treatment with the indicated analogs of WX-02-46 and inactive enantiomer WX-02-26 (20 µM, 2 h) on WX-01-08 (5 μM, 1 h) engagement of recombinant HCN1 in HEK293T cell lysates. Data are from a single experiment. **(I)** Chemical structures of HCN ligand WX-02-679 and inactive enantiomer WX-02-678. **(J-L)** Gel-ABPP data **(J, K)** and quantification of data **(L)** comparing potencies of parent (WX-02-26 and WX-02-46) and analog (WX-02-678 and WX-02-679) stereoprobes at blocking WX-01-08 (5 μM, 1 h) engagement of HCN1 in HEK293T cells. IC50_WX-02-46_ = 14.4 μM, 95% CI = 10.9 – 18.6 μM; and IC50_WX-02-679_ = 4.4 μM, 95% CI = 3.5 – 5.2 μM. Data are average values ± SD from two or three independent experiments.

A review of HCN structures revealed that the conserved stereoprobe-liganded cysteine is located in the cyclic nucleotide-binding domain (CNBD) of HCN channels in close proximity to the binding pocket for cAMP (**Figure 5D**).^62–64^ The CNBD plays an important role in modulating HCN function by allosterically converting cAMP binding to accelerated channel activation kinetics and a shift in the conductance voltage curves toward more depolarized potentials.^63,65,66^ We found that WX-01-08 reactivity with recombinant HCN1 was blocked in a concentration-dependent manner by cAMP at concentrations as low as 0.5 µM (**Figure 5E**). cAMP also blocked WX-01-08 reactivity with HCN2 and HCN4, but at somewhat higher concentration (≥ 5 µM) (**Figure S5D, E**). In contrast, the HCN pore-blocking compounds Ivabradine^67,68^ and RO-275^69^ did not affect WX-01-08-HCN1 interactions either in cell lysates or in cells (**Figures 5F, G** and **S5F, G,** respectively). We interpret these results to indicate that tryptoline acrylamide stereoprobes target a distinct domain (the CNBD) compared to previously described ligands (direct pore blockers) for HCN channels. We next set out to investigate the functional consequences of stereoprobe binding to the CNBD of HCN channels.

### Stereoprobes block cAMP-induced modulation of HCN channels

To facilitate functional studies of HCN channels, we first screened a focused set of analogs of WX-02-46 (and their respective enantiomers) for interactions with HCN1 by gel-ABPP, which identified a compound WX-02-679 bearing a biaryl appendage in place of the propyl group of WX-02-46 that displayed improved potency with retained stereoselectivity (**Figures 5H, I** and **S5H**). Using gel-ABPP, we measured IC_50_ values of 4.4 and 14 µM for engagement of HCN1 by WX-02-679 and WX-02-46, respectively (**Figure 5J-L**). WX-02-679 also engaged HCN2 and HCN4, with comparable potency (5.1 and 3.7 µM) (**Figure S5I**). We did not further investigate HCN3 in these studies, as this isoform was not detected in our platform, and prior reports have shown that the CNBD of HCN3 does not substantially regulate channel activity.^70,71^

We next evaluated the impact of WX-02-679 on the functional properties of HCN channels expressed in HEK293T cells by patch clamp electrophysiology. In these experiments, we used HCN2-expressing cells—due to the larger cAMP-induced activation shift in HCN2 compared to HCN1— pre-treated with WX-02-679 or control enantiomer WX-02-678 (10 µM each, 20 min) or DMSO, followed by whole-cell patch clamp recordings (**Figure 6A**). Consistent with previous studies, the addition of cAMP in the pipette solution induced a rightward shift in the voltage-dependent activation of mouse HCN2 to more depolarizing potentials (**Figure 6B**).^65,72^ Data fitting to the Boltzmann equation (see Materials and Methods) yielded mean half-activation voltage (V_1/2_) for mouse HCN2: control: −92.1 ± 2.0 mV; control + cAMP: −83.5 ± −0.8 mV; cAMP + WX-02-679: −95.0 ± 1.8 mV; cAMP + WX-02-678: −84.0 ± −0.8 mV (**Figure 6C** and **Dataset S1**). Notably, this cAMP effect was near-completely blocked in cells pre-treated with WX-02-679, but not WX-02-678 (**Figure 6B, C**). WX-02-679 also blocked the cAMP effects on mouse HCN1 and rabbit HCN4 (**Figures 6D** and **S6A, B**), which yielded half-activation voltage (V_1/2_) for mouse HCN1: control: −72.6 ± 0.6 mV; control + cAMP: −64.2 ± 1.9 mV; cAMP + WX-02-679: −71.6 ± 1.1 mV; cAMP + WX-02-678: −65.7 ± 1.1 mV; and rabbit HCN4: control: −103.6 ± 1.2 mV; control + cAMP: −85.3 ± 0.9 mV; cAMP + WX-02-679: −106.1 ± 1.7 mV; cAMP + WX-02-678: −86.3 ± 1.8 mV (**Figure 6D** and **Dataset S1**). The rabbit ortholog of HCN4 was used due to more consistent recombinant expression of this paralog in cells. We interpret the patch clamp data to indicate that WX-02-679 specifically disrupts the cAMP-modulated, but not basal function of HCN channels. We also attempted to assess the impact of WX-02-679 on cysteine mutant forms of each HCN channel (e.g., hHCN1_C542A mutant) but found that these variants exhibited poor or cAMP-independent channel conductance (**Figure S6A, C**), despite similar overall expression compared to WT channels (e.g., see **Figures 5F** and **S5B, C**). We do not yet understand why the cysteine mutant HCN channels were basally dysfunctional, but it is possible that these variants disrupt cAMP binding, impair proper trafficking to the plasma membrane—a process the CNBD has been reported to play a role in^73^—or both.

**Figure 6.**
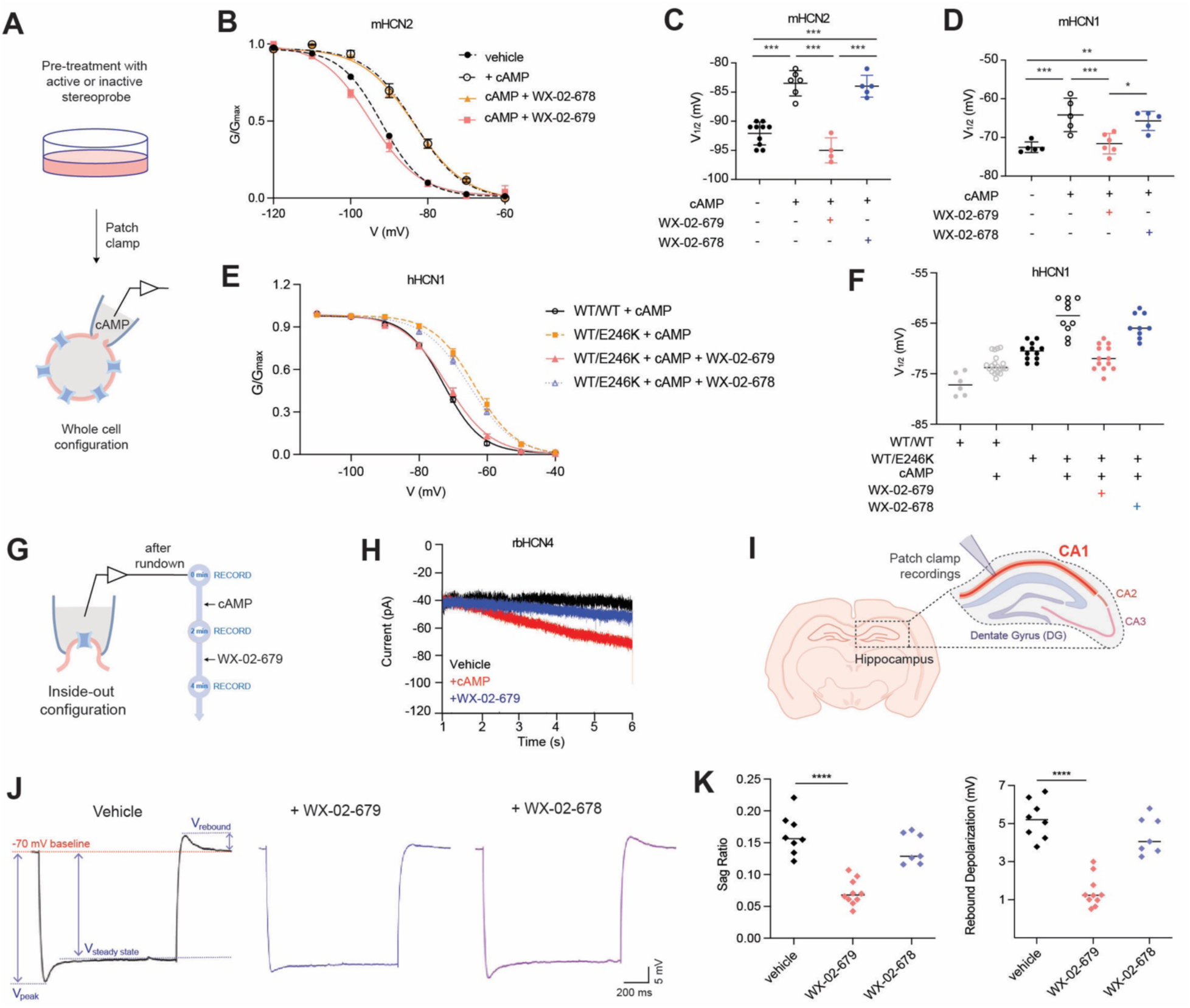
WX-02-679 blocks cAMP-induced changes in HCN channel activation. (A) Workflow of patch clamp electrophysiology experiments performed on HEK293T cells recombinantly expressing HCN channels, where cells are first pre-treated with stereoprobes for 20 min at 37 °C, followed by patch clamp recordings with cAMP added in the patch pipette in the whole cell configuration. **(B)** Mean activation curves measured in HEK293T cells recombinantly expressing mouse (m)HCN2 after pre-incubation with WX-02-678 or WX-02-679 (10 μM, 20 min) or DMSO followed by addition of cAMP (100 μM) in the patch pipette. Data are average values ± SEM from at least two independent experiments (≥4 cells for each condition per experiment), and fit to the Boltzmann equation plotted as lines. **(C)** and **(D)** Half activation potential (V_1/2_) quantified from activation curves from whole cell patch clamp recordings of HEK293T cells recombinantly expressing mHCN2 and mHCN1, respectively. Mouse orthologs of HCN1 and HCN2 were used due to improved protein expression in cells. Data are average values ± SEM from at least two independent experiments (≥4 cells for each condition per experiment). **(E)** Mean activation curves measured in HEK293T cells recombinantly expressing WT/WT hHCN1 homomers or WT/E246K hHCN1 heteromers after pre-treatment with DMSO or WX-02-679 (20 μM, 20 min) followed by addition of cAMP (200 nM) in the patch pipette. Data are average values ± SEM from at least two independent experiments (≥4 cells for each condition per experiment), and fit to the Boltzmann equation are plotted as lines. **(F)** Half activation potential (V_1/2_) quantified from activation curves from whole cell patch clamp recordings of HEK293T cells recombinantly expressing WT/WT hHCN1 homomers or WT/E246K hHCN1 heteromers after pre-treatment with WX-02-678 or WX-02-679 (20 μM, 20 min) or DMSO followed by addition of vehicle or cAMP (200 nM) in the patch pipette. Data are average values ± SEM from at least two independent experiments (≥4 cells for each condition per experiment). **(G)** Workflow of patch clamp electrophysiology in inside-out configuration that exposes the cytosolic side of the channel to the bath solution. The ligands cAMP (100 uM) and WX-02-679 (20 uM) were added to the bath solution at the end of the current run down. **(H)** Inside-out patch clamp recordings of current traces at -105 mV on HEK293T cells recombinantly expressing rabbit (rb)HCN4 (vehicle, black) showing inhibition of cAMP-induced effects (red) on HCN4 channel activity by WX-02-679 (blue). Shown here is the full effect of WX-02-679 (20 uM) that was recorded after 2 minutes of cAMP followed by 2 minutes of WX-02-679 exposure. Data are from a single experiment representative of at least two independent experiments. **(I)** Schematic of coronal section of mouse brain showing CA1 pyramidal layer used for *ex vivo* brain slice patch clamp electrophysiology experiments in mouse hippocampal neurons. **(J)** and **(K)** Voltage sag ratio (V_sag_, left) and rebound depolarization (V_rebound_, right) recordings and quantification in CA1 neurons from mouse hippocampal slices showing stereoselective effects by WX-02-679 (20 μM, 6 min). V_rebound_ measured as positive deflection from baseline (-70 mV). V_sag_ measured as (V_peak_-V_steady state_)/V_peak_). **(J)** Representative traces. **(K)** Quantification of data. Data are average values ± SEM (vehicle: N = 8, WX-02-679: N = 10, WX-02-678: N = 7).

Among epilepsy-associated pathogenic HCN1 variants, a high prevalence exhibit gain-of-function phenotypes that right shift the channel activation curve.^74^ A cAMP antagonistic peptide derived from neuronal protein TRIP8b, termed TRIP8bnano, has been shown to partly or fully rescue these gain-of function properties in HEK293T cells.^75^ This prompted us to investigate whether the allosteric modulatory effects of WX-02-679 can similarly rescue these pathogenic HCN1 variants. Electrophysiological studies of HEK293T cells expressing the recently identified heterozygous HCN1 variant WT/E246K^75^ that results in a right shift in V_1/2_ (+10 mV) revealed that pre-treatment with WX-02-679 (20 µM, 6 min) stereoselectively restored the voltage-dependent activation of this mutant channel to that observed for the WT/WT HCN1 channel in the presence of cAMP (**Figures 6E, F**). We then evaluated the WT/M153I HCN1 variant, which is linked to severe childhood epilepsy and causes a much larger right shift in activation potential (+ 18 mV),^75,76^ and found that pre-treatment with WX-02-679 also partially rescued this severe voltage shift (**Figure S6D**).

We finally confirmed the direct effect of WX-02-679 on cAMP modulation of HCN4 by performing inside-out patch clamp experiments (**Figure 6G**). In this experiment, cAMP (100 µM) was provided from the intracellular side of the channel (i.e. in the bath solution) after the run down of the current was completed, and the cAMP effect was evaluated as current increase at the single voltage step of - 105 mV (**Figure 6H**). Further addition of 20 µM WX-02-679 fully recovered the cAMP-induced increase in current within 2 minutes from application (**Figure 6H**).

We were next interested in studying the impact of WX-02-679 on endogenous HCN channel function, which we investigated by whole cell patch clamp recordings in CA1 pyramidal neurons in mouse hippocampal slices (**Figure 6I**). These neurons express a mix of HCN1 and HCN2 channels that underlie the *I_h_* current, which regulates several electrophysiological properties.^77–79^ Activation and deactivation of HCN channels, induced by current injections, elicits in these cells well-characterized “voltage sag” (V_sag_) and “rebound depolarization” (V_rebound_) responses.^77^ The sag ratio—reflecting the extent of the voltage sag relative to the peak hyperpolarization—quantifies the magnitude of *I_h_* activation and is a canonical readout of HCN channel function in neurons. Similarly, the rebound depolarization that follows termination of the hyperpolarizing step reflects the transient inward current carried by HCN channels upon deactivation and is indicative of the level of *I_h_* in the cell. We found that hippocampal slices pre-incubated with WX-02-679 (20 µM, 6 min) exhibited a stereoselective decrease in V_sag_, V_rebound_, and sag ratio (**Figure 6J, K**) indicating a reduction in *I_h_*. When we examined the passive properties of the membrane, we further observed a stereoselective decrease in resting membrane potential (RMP) and increase in input resistance (IR) (**Figure S6E**), consistent with loss of the tonic depolarizing *I_h_* that normally counteracts membrane hyperpolarization and maintains background conductance.^80,81^ The consequent stereoselective increase in firing frequency (**Figure S6F**) further reflects the elevated input resistance (R_input_), as the cell is rendered more excitable to equivalent synaptic input. Collectively, the stereoselective effects on sag ratio, rebound depolarization, RMP, R_input_, and firing frequency support a model where WX-02-679 decreases endogenous HCN channel activity in CA1 pyramidal neurons, corroborating the hyperpolarizing effects on the channel activation curve observed in heterologous expression systems. We additionally measured a range of electrophysiological properties in hippocampal slices related to single action potential morphology that are typically unaffected by HCN channels and found that none were substantially altered by WX-02-679 (**Figure S6G**), further supporting on-target activity for this compound in native neuronal contexts.

Altogether, our findings demonstrate that WX-02-679 is a distinct chemical tool for studying HCN channels that targets their cAMP-regulated gating mechanism rather than acting as direct pore blockers.

## DISCUSSION

Efforts to discover ligands for human proteins have benefited from advances in both screening technologies and small-molecule library designs.^82,83^ Here, we have attempted to address another important bottleneck, namely the breadth of proteins, and states of those proteins, that are assayed for interactions with small molecules. By extending the reach of ABPP beyond readily accessible immortalized cell lines to include primary cells from the adult brain, we have identified covalent ligands for several CNS-enriched proteins. The brain contains diverse neuronal and glial populations^19,84,85^, and, while we do not yet know from which of these cell types the stereoprobe-liganded proteins originate, extrapolation from public RNAseq data would indicate our brainocyte ligandability maps contain both neuron-(e.g., HCN channels) and glial-(e.g., PLP1) derived proteins. Complementary sources of CNS proteins for future chemical proteomic investigations may include brain organoids and iPSC-derived brain cells.^86,87^ Considering further that tissue and cell type-dependent expression represents one of the largest sources of proteomic diversity, we hope that our findings will inspire the establishment of ligandability maps for additional organs in the mammalian body.

Multiple features of the brainocyte ligandability maps highlight attributes of screening endogenous proteins for interactions with small molecules. First, several stereoprobe-protein interactions were identified in brainocytes, but not brain lysates, suggesting that these liganding events depend on the environment of the intact cell. Curiously, we also noted some stereoprobe-protein interactions that showed the opposite profile of exclusive ligandability in brain lysates (**Figure 1G** and **Dataset S2**). While we do not yet understand the basis for this *in vitro* ligandability profile, it is possible that, for a kinase like CDK5, competition with ATP may preclude stereoprobe binding in intact brainocytes.

Our ABPP studies of brainocytes also identified a remarkable example of complexoform-restricted liganding of DPYSL2, which occurred exclusively when this adaptor protein was co-expressed with one of its paralogs DPYSL5. DPYSL2 is a CNS-enriched protein with additional expression in immune cells, and, interestingly, we had quantified this protein, but not observed evidence of specific stereoprobe interactions, in past ABPP experiments performed in the B lymphocyte cell line Ramos.^13^ Once having identified herein that DPYSL2 interactions with tryptoline acrylamide stereoprobes requires the co-expression of DPYSL5, we identified brain cell lines (human: SH-SY5Y; mouse: Neuro2A) that express high levels of endogenous DPYLS5 and showed clear evidence of stereoprobe liganding of DPYSL2 in these cells. Considering further that tryptoline acrylamides engage a paralog-restricted cysteine (C504) in DPYSL2 that is located in proximity to the homo/hetero-oligomeric interface of the protein, we interpret our findings to indicate that the DPYSL2:DPYSL5 complex may create a unique ligandable pocket proximal to DPYSL2_C504. More generally, our results illuminate how cell context can shape not only protein-protein, but also protein-small molecule interactions, and provide further evidence for the potential to develop complexoform-selective covalent ligands.^15^ Future studies are required to determine how tryptoline acrylamides may affect the functions of DPYSL2:DPYSL5 complexes, which have been implicated in critical cell biological processes like microtubule transport and axon elongation.^88^

Our functional studies of the HCN family support that tryptoline acrylamide ligands offer a differentiated set of chemical tools for studying this biomedically important class of ion channels. Unlike more classical direct pore blockers,^68^ including the FDA-approved drug ivabradine – a pan-HCN inhibitor for management of chronic heart failure – the tryptoline acrylamides target a conserved cysteine in the allosteric CNBD and exhibit an intriguing pharmacological profile of suppressing cAMP-induced shifts in HCN activation while sparing basal channel function. Considering further that the gain-of-function channel phenotypes of some pathogenic HCN1 variants causing epilepsy and neurodevelopmental delay^75,76^ can seemingly be corrected to variable degrees by tryptoline acrylamide WX-02-679 and the peptide TRIP8bnano,^88,89^ we speculate that such CNBD modulators may offer an attractive alternative class of therapeutic agents for targeting HCN channels. Indeed, individual HCN isoforms have differential sensitivity to the CNBD domain modulation,^65,73^ and ligands targeting this domain may accordingly exert more diverse subtype selectivity than pan-pore blockers like ivabradine. Other complementary approaches include efforts to generate subtype-selective pore blockers.^69,89–92^

A more complete understanding of the potential for CNBD-targeting agents to modulate the functions of HCN channels in (patho)physiological settings would benefit from structural studies of stereoprobe-HCN complexes, as well as more advanced ligands optimized for *in vivo* activity. We have recently achieved the latter goal for tryptoline acrylamides targeting the BAX adaptor protein,^93^ but this work did not require that the compounds exhibit CNS penetrance. If new chemotypes are required for the generation of CNS-penetrant CNBD ligands, we have shown that stereoprobes can serve as the basis for high-throughput assays for screening larger compound libraries against allosteric pockets on proteins.^94,95^ More generally, our findings provide another compelling example of the diverse ways that allosteric ligands can regulate ion channel function,^96–98^ in this instance, providing tools to study the specific effects of cyclic nucleotides on HCN function while leaving basal channel activity intact.

### Limitations of the study

Ligandability maps of disassociated brain cell preparations may not account for small molecule-protein interactions that are posttranslationally regulated *in vivo* by dynamic forms of cell-cell communication (e.g., transsynaptic signaling). Rare cell types also may represent too small of a percentage of the cell suspensions prepared from whole brain for deep ligandability mapping. These challenges may be addressed in the future by performing ABPP experiments on acute slices from specified brain regions or brain organoid cultures. Many of the stereoprobe liganding events occur on proteins lacking orthosteric pockets (e.g., DPYSL2, PLP1) or occur at non-orthosteric sites on proteins (e.g., PDE7B). Characterizing the functional impact of these non-orthosteric small molecule-protein interactions can be technically challenging, especially for proteins that lack straightforward activity assays. Even in cases where stereoprobe liganding events are not found to exert a functional effect, they can be converted into heterobifunctional compounds to promote, for instance, degradation of target proteins.^99^ Finally, many pockets in the proteome may lack proximal cysteines, and future studies of brainocytes with electrophilic^37,100–118^ or photoreactive^119,120^ small molecules targeting alternative amino acids should further expand the scope of ligandable CNS-enriched proteins.

### Significance

The ligandability maps generated by activity-based protein profiling (ABPP) have, to date, been restricted to readily accessible cell types (cell lines or primary immune cells), leaving proteins with tissue-restricted expression—such as those enriched in the nervous system— underexplored for small molecule interactions. In this study, we established an ABPP protocol for intact primary cells freshly dissociated from adult mouse brain (“brainocytes”) and used this method to generate global maps of protein interactions for sets of stereochemically defined electrophilic small molecules (stereoprobes). We identified stereoprobe liganding events for proteins for diverse structural and functional classes, many of which also showed nervous system-enriched expression and had not been observed in prior chemical proteomic studies of peripheral cell types. Among these, we identified tryptoline acrylamide stereoprobes that: i) engage the adaptor protein DPYSL2 in a complexoform-restricted manner dependent on its assembly with a paralogous protein partner; and ii) target a conserved allosteric cysteine in the cyclic nucleotide-binding domain of HCN channels to selectively disrupt cAMP-dependent gating while sparing basal channel activity. Together, these findings establish a robust scalable strategy for globally mapping covalent small molecule-protein interactions in intact primary brain cell populations and demonstrate its utility for discovering ligands that perturb proteins with key roles in CNS physiology and disease.

## Supporting information

Supplementary Chemistry Information

## Resource Availability

### Lead contact

Further information and requests for resources and reagents should be directed to and will be fulfilled by the Lead Contact, Benjamin F. Cravatt.

### Materials availability

All chemical probes and other elaborated electrophilic compounds in this study are available from the Lead Contact with a completed Materials Transfer Agreement.

### Data and code availability

The mass spectrometry proteomics data have been deposited to the ProteomeXchange Consortium^121^ via the PRIDE^122^ partner repository with the dataset identifier PXD082934. Previously published datasets^13^ relevant to this study are under data identifier PXD042541. Raw proteomic files were searched using the ProLuCID algorithm using a reverse concatenated, non-redundant variant of the Human UniProt database (release 2016-07) or Mouse UniProt database (release 2017-07). Processed proteomic data are provided in Supporting Dataset 2.

## Acknowledgements

This work was supported in part by the Claudia Skaggs Luttrell Endowed Fellowship, the Lundbeck La Jolla Research Center, and a Toni Rosenberg Memorial Fellowship. We thank Dr. Xuedong Liu and Dr. Bing Chen (WuXi AppTec) for small-molecule synthesis, Dr. Melissa Dix for assistance with mass spectrometry-based proteomics experiments, Quynh Nguyen Wong, Jillian Smith, Jason Lee, Catherine Chiang, Dr. Brandon Orzolek (Scripps Automated Synthesis Facility), and Dr. Chris Reinhardt for support with high-resolution mass spectrometry, Brian Seegers (Scripps flow core facility) for assistance with flow cytometry, Steven A. Siegelbaum (Columbia University) for assistance and supervision of whole cell patch clamp electrophysiology studies in acute brain slices. The funders had no role in study design, data collection and analysis, decision to publish or preparation of the manuscript.

## Author contributions

E.Y. and B.F.C. conceived the study. E.Y. prepared brain lysate, disassociated brain cells, and acute brain slice samples and performed flow cytometry analyses. E.Y. generated the proteomic data. E.Y., B.M. and B.F.C. performed analysis of the proteomic data. E.Y. performed cloning and generated all CRISPR/Cas9 knockout and lentiviral transduced stable cell lines. E.Y., G.W., D.S., and S.Q. performed gel-based profiling and western blotting. E.Y. performed IP-MS experiments. E.Y. performed LC MS/MS enzymatic assays. A.R., X.J. and A.M. performed in vitro whole cell patch clamp electrophysiology experiments in cell lines. R.C. and A.M. performed *ex vivo* whole cell patch clamp electrophysiology experiments in brain slices. X.J. and A.M. performed inside-out patch clamp electrophysiology experiments. G.S. generated list of CNS enriched proteins. B.M. supervised compound synthesis and characterization. J.L.B. provided support and resources. E.Y., B.M. and B.F.C. wrote and edited the manuscript. B.F.C. supervised this study.

---

**Figure S1.**
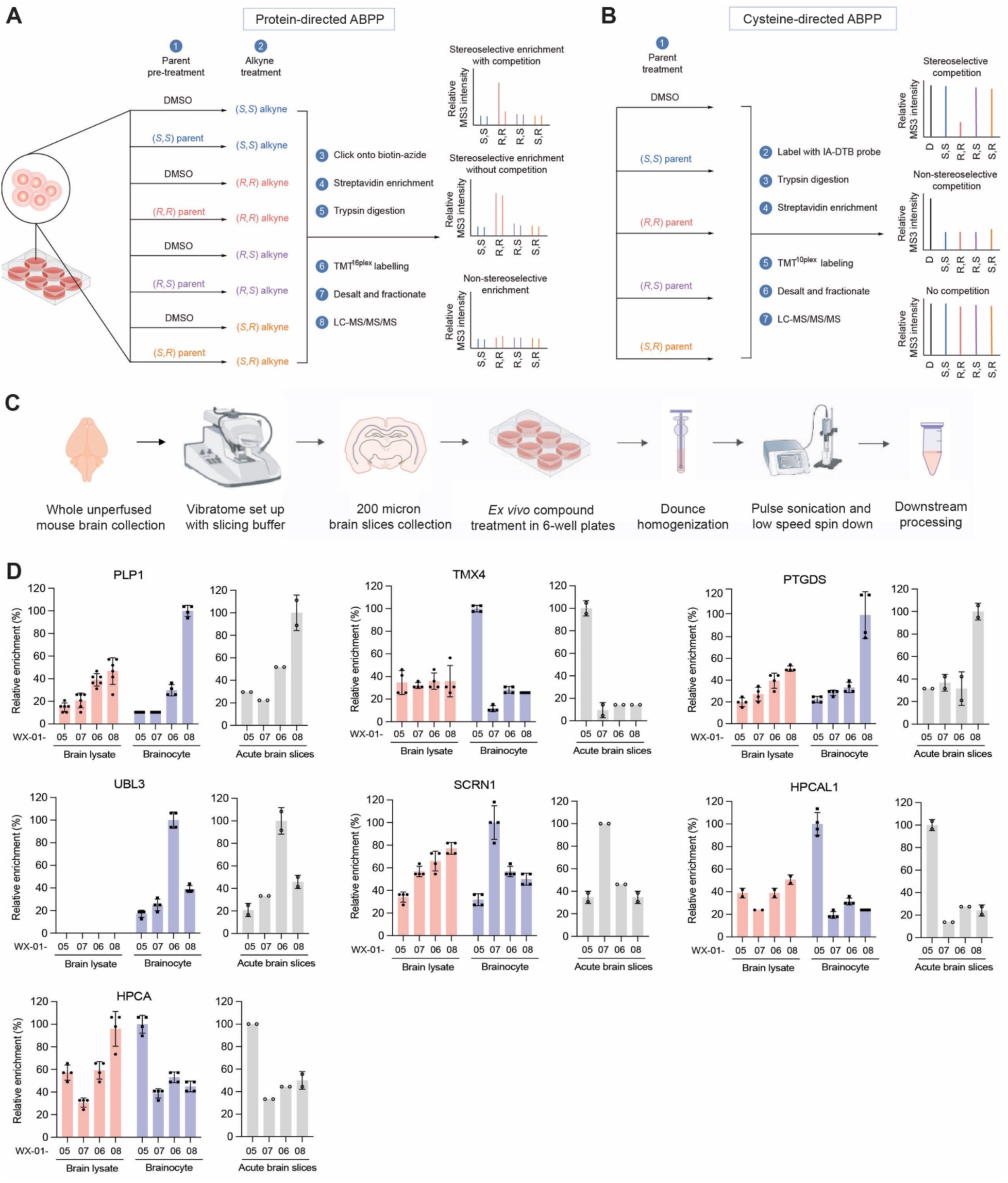
An ABPP protocol for analyzing intact mouse brain cells and slices, related to Figure 1. **(A)** Workflow for protein-directed ABPP experiments where the stereoselective enrichment of proteins by alkyne stereoprobes and blockade of this enrichment by pre-treatment with corresponding non-alkyne parent stereoprobes are determined by multiplexed (TMT^16plex^) MS-based proteomics, as previously described.^13^ **(B)** Workflow for cysteine-directed ABPP experiments where stereoprobe liganding events are determined by blockade of reactivity of cysteines with a broad-spectrum iodoacetamide-desthiobiotin (IA-DTB) probe as measured by multiplexed (tandem mass tagging, TMT^10plex^) MS-based proteomics, as previously described.^13^ **(C)** Workflow for preparation of acute brain slices collected from adult C57BL/6 mice and analysis by protein-directed ABPP. Unperfused whole brains from adult C57BL/6 mice were harvested and sliced using a vibratome in oxygenated slicing buffer. Six 200 micron brain slices were seeded in 6-well plates in 2 mL DMEM per well for alkyne stereoprobe treatment for 1 h at 37 °C. Slices were collected and Dounce homogenized in cold DPBS, then further lysed by probe sonication. Lysate was spun down at low speed (3000 g, 5 min) to remove debris for ABPP experiments. **(D)** Protein-directed ABPP data showing representative proteins enantioselectively enriched by the indicated stereoprobes in brainocytes but not brain lysates, left; showing similar enantioenrichment profiles in acute brain slices, right (PLP1: WX-01-08; UBL3: WX-01-06; PTGDS: WX-01-08; HPCAL1: WX-01-05; TMX4: WX-01-05; SCRN1: WX-01-07; HPCA: WX-01-08). Data represents average values ± SD from two to six independent experiments normalized to percent of maximum mean values.

**Figure S2.**
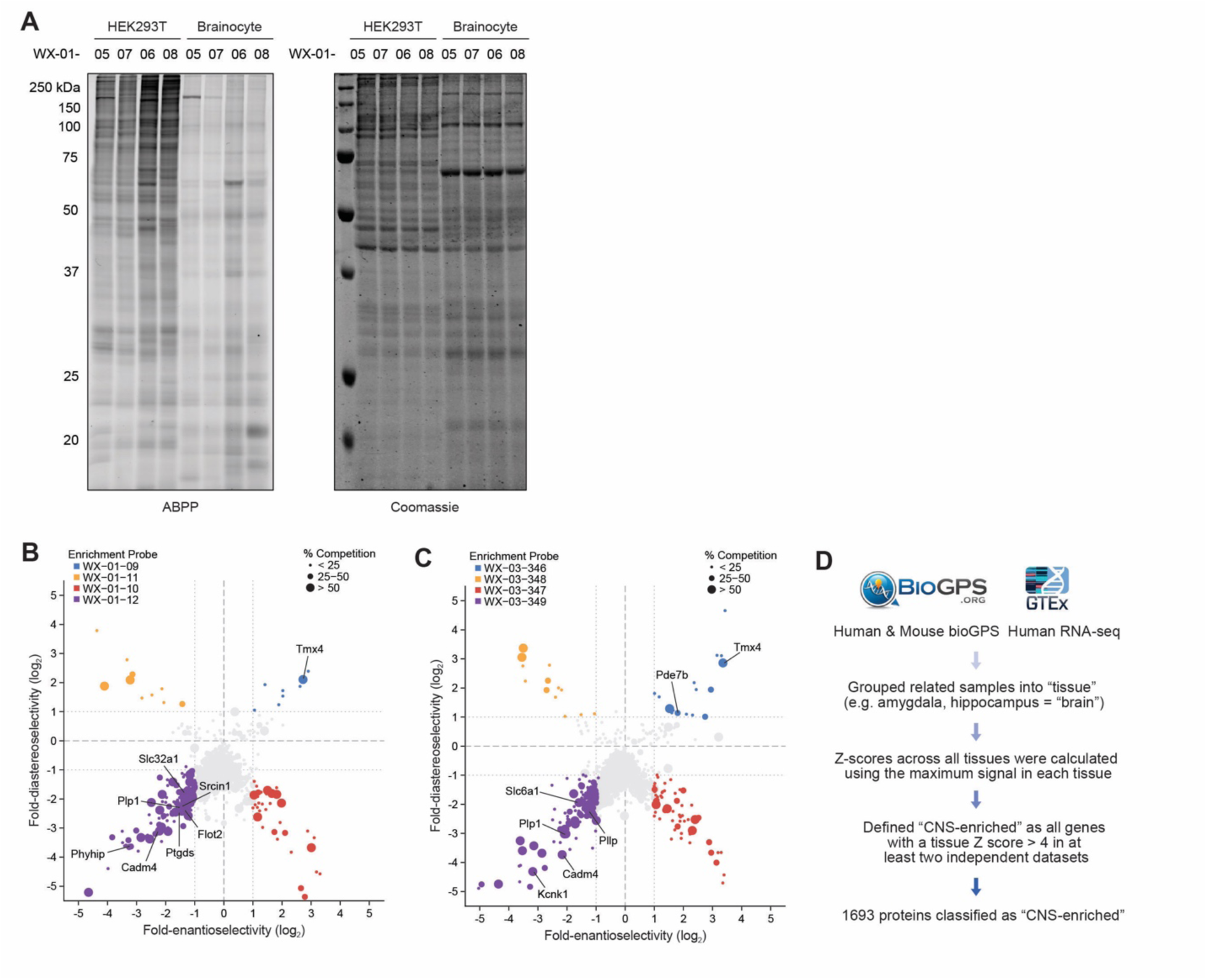
Stereoprobe ligandability maps of mouse brainocytes, related to Figure 2. **(A)** Gel-ABPP data comparing stereoprobe reactivity profiles of mouse brainocytes and HEK293T cells. Each cell preparation was treated with alkynylated tryptoline acrylamides WX-01-05/06/07/08 (5 μM, 1 h), and stereoprobe-reactive proteins were visualized by conjugation to an azide-rhodamine reporter group, SDS-PAGE, and in-gel fluorescence scanning. Left, ABPP data showing lower overall stereoprobe reactivity in brainocytes compared to HEK293T cells. Right, Coomassie blue staining showing similar overall protein amounts for brainocyte and HEK293T cell preparations. Data are from a single experiment representative of at least two independent experiments. **(B), (C)** Quadrant plots highlighting stereoselectively liganded proteins for each stereoconfiguration of the indicted alkyne stereoprobe sets. See Figure 2B-D for further description of quadrant plots. **(D)** Method used to generate a list of CNS-enriched proteins using human bioGPS, mouse bioGPS, and GTEx data.^40,41^ Proteins with a tissue Z score > 4 for the brain in at least two independent datasets were classified as “CNS-enriched”.

**Figure S3.**
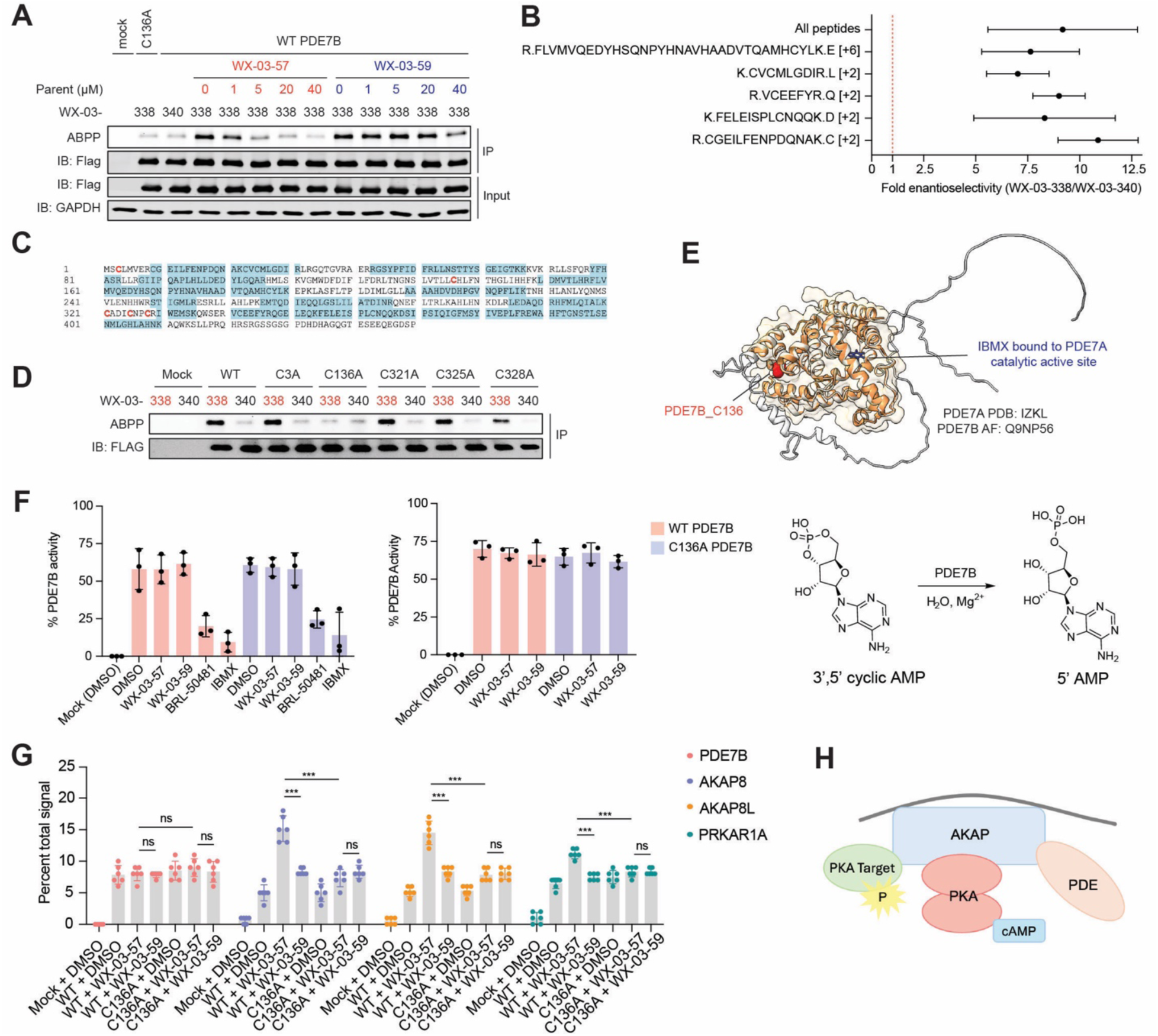
Characterization of representative CNS-enriched liganded proteins, related to Figure 3. **(A)** Gel-ABPP data showing concentration-dependent and stereoselective blockade of WX-03-338 (5 μM, 1 h) engagement of recombinant FLAG-tagged PDE7B by pre-treatment with WX-03-57 (indicated concentrations, 2 h) in HEK293T cells. ABPP was performed on protein samples after an anti-FLAG IP step. Data are from a single experiment representative of at least two independent experiments. **(B)** Stereoselective enrichment of cysteine-containing and all tryptic peptides from PDE7B by WX-03-338 (5 μM, 1 h) quantified from protein-directed ABPP experiments in SH-SY5Y cells stably expressing WT-PDE7B. Data are average values ± SD from two independent experiments. **(C)** Sequence of PDE7B with all quantified tryptic peptides from (B) highlighted in blue. Cysteines not detected in this experiment (C3, C136, C321, C325, C328) are bolded in red. **(D)** Gel-ABPP data showing stereoselective engagement of recombinant WT and C3A, C321A, C325A, and C328A mutant PDE7B proteins, but not a C136A-PDE7B mutant, by WX-03-338 (5 μM, 1 h) in HEK293T cells. ABPP was performed on protein samples after an anti-FLAG IP step. Data are from a single experiment representative of at least two independent experiments. **(E)** Crystal structure of the catalytic domain of PDE7A (PDB: 1ZKL^128^) represented in light orange overlaid with AlphaFold2 structure of PDE7B (AF-Q9NP56-F1{Citation}) represented in white, showing residue PDE7B_C136 (highlighted in red, paralogous to PDE7A_F175) is distal to the IBMX-binding site (IBMX highlighted in blue) of the protein. **(F)** Bar graph showing 3’,5’ cAMP hydrolytic activity of lysates of mock HEK293T cells or HEK293T cells recombinantly expressing WT- or C136A-PDE7B showing the inhibitory effects of established PDE inhibitors BRL-50481 (200 μM, 2 h) and IBMX (100 μM, 2 h), but a lack of inhibitory effects for stereoprobe WX-03-57 applied either to cell lysates (20 μM, 2 h, left graph) or intact cells (20 μM, 3 h, right graph). Reaction schematic of PDE7B-catalyzed hydrolysis of 3’,5’ cAMP to 5’ AMP is shown on the far right. **(G)** Quantification of the co-enrichment of the indicated proteins in anti-FLAG IP-MS experiments performed with SH-SY5Y cells expressing FLAG-tagged WT- or C136A-PDE7B treated with WX-03-57 or WX-03-59 (20 μM, 3h), showing stereoselective and site-specific increased association of PDE7B with AKAP8, AKAP8L, and PRKAR1A. Data are average values ± SD from six independent experiments. Asterisks designate statistical significance from paired sample t-tests for the indicated comparisons. **(H)** Representative schematic of AKAP-PDE-PKA complex.

**Figure S4.**
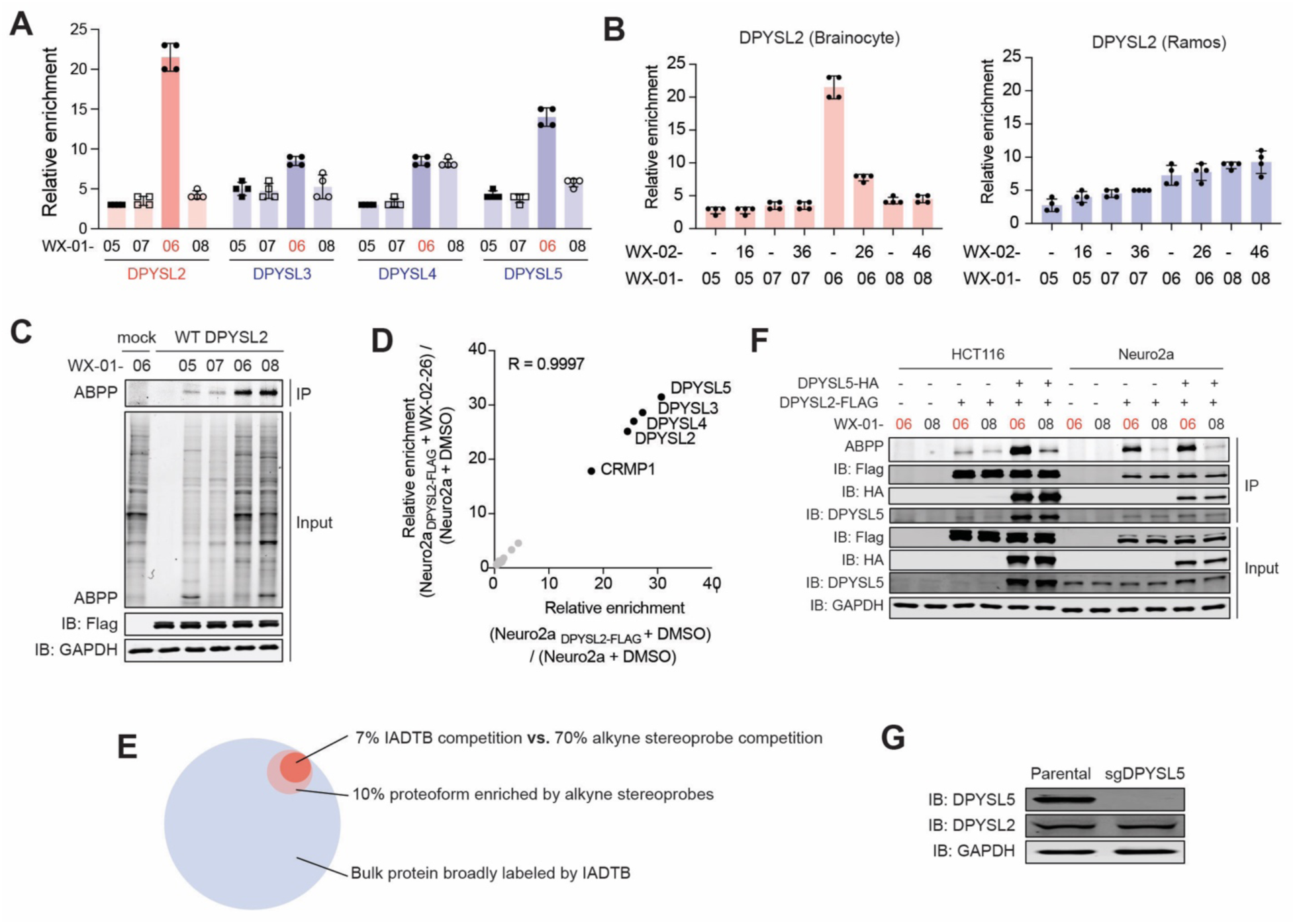
Characterization of complexoform-restricted stereoprobe ligands for DPYSL2, related to Figure 4. **(A)** Protein-directed ABPP data for DPYSL2, DPYSL3, DPYSL4, and DPYSL5 showing strong stereoselective enrichment of DPYSL2, modest stereoselective enrichment of DPYSL5, and no stereoselective enrichment of DPYSL3 and DPYSL4 by WX-01-06 (5 μM, 1 h) in brainocytes. Data are average values ± SD from four independent experiments. **(B)** Comparison of protein-directed ABPP data showing stereoselective enrichment of DPYSL2 by WX-01-06 (5 μM, 1 h) and blockade of enrichment by WX-02-26 (20 μM, 2 h) in brainocytes (left), but no such enrichment in Ramos cells (right). Data are average values ± SD from four independent experiments. **(C)** Gel-ABPP data showing a lack of stereoselective engagement of FLAG-tagged WT-DPYSL2 by WX-01-06 compared to enantiomer WX-01-08 (5 µM, 1 h) in HCT116 cells. Data are from a single experiment representative of at least two independent experiments. **(D)** Proteins co-enriched with DPYSL2 in anti-FLAG IP-MS experiments performed with parental Neuro2a cells (control) or DPYSL2-expressing Neuor2A (Neuro2a_DPYSL2-FLAG_) cells treated with DMSO or WX-02-26 (20 µM, 3 h). Data are average values for four independent experiments. **(E)** Schematic mode for explaining the divergent protein-directed and cysteine-directed ABPP data for DPYSL2 due to proteoform (complexoform)-restricted liganding by WX-01-06/WX-02-26. **(F)** Gel-ABPP data showing stereoselective engagement of recombinant WT-DPYSL2 in Neuro2a cells, but not HCT116 cells by WX-01-06 (5 µM, 1 h), correlating with the expression profile of endogenous DPYSL5 as determined by western blotting. Data are from a single independent experiment. **(G)** Western blot analysis comparing the indicated DPYSL protein signals in parental and sgDPYSL5 SH-SY5Y cells. Data are from a single independent experiment.

**Figure S5.**
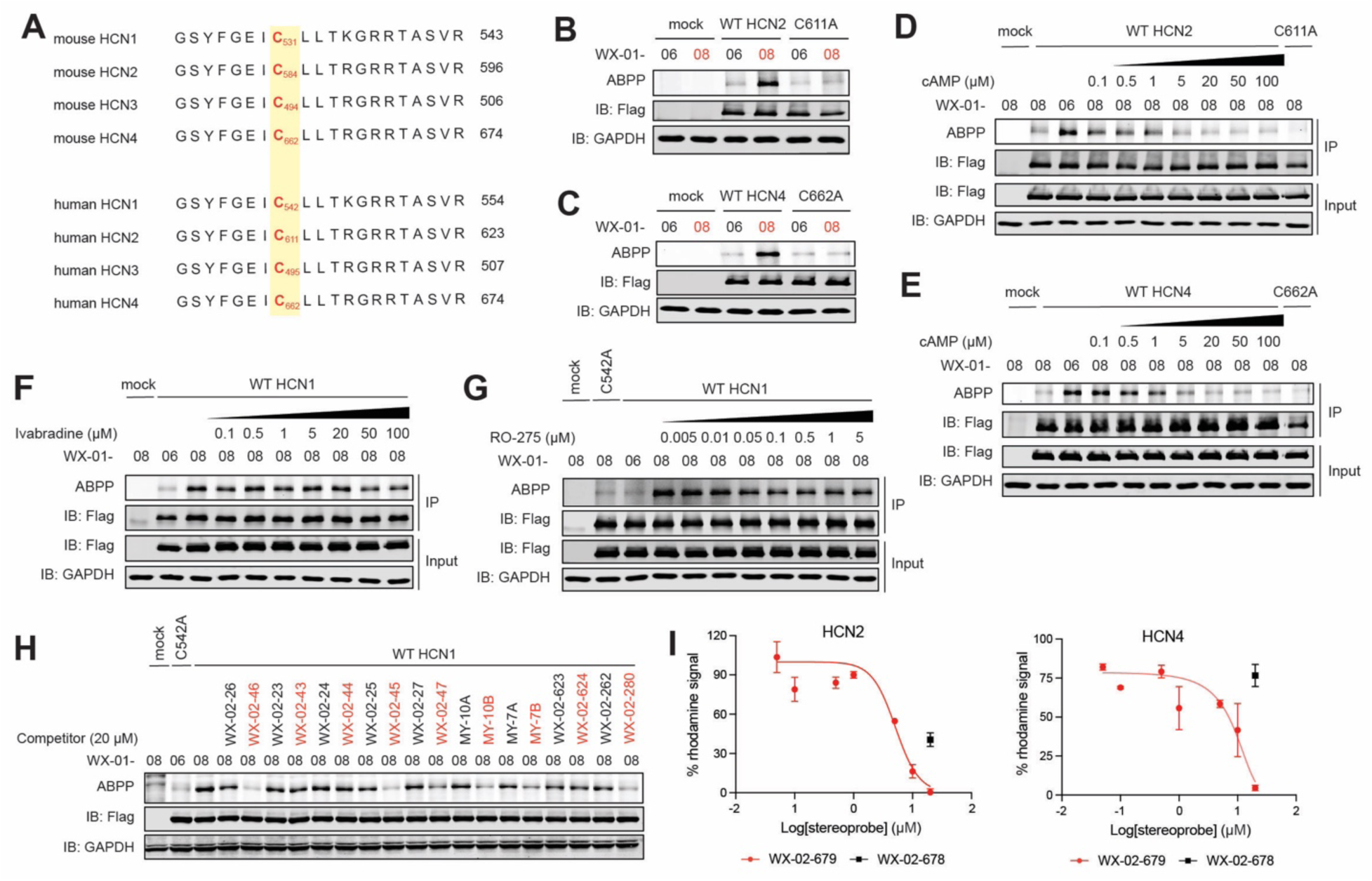
Characterization of stereoprobe ligands targeting the cyclic nucleotide binding domain of HCN channels, related to Figure 5. **(A)** Sequence alignment of the indicated amino acid region of mouse and human HCN1, HCN2, HCN3, and HCN4 (numbers on right refer to the C-terminal residue number for each sequence), showing conservation of mouse HCN1_C531 (in red) and surrounding residues across HCN paralogs and orthologs. **(B)** and **(C)** Gel-ABPP data showing stereoselective engagement of recombinant FLAG-tagged WT-HCN2 but not a C611A-HCN2 mutant **(B)**, and WT-HCN4 but not C662A-HCN4 mutant **(C)** by WX-01-08 (5 μM, 1 h) in HEK293T cells. Data are from a single experiment representative of at least two independent experiments. **(D)** and **(E)** Gel-ABPP data showing concentration-dependent blockade of WX-01-08 (5 μM, 1 h) of WX-01-08 (5 μM, 1 h) engagement of recombinant HCN2 **(D)** and HCN4 **(E)** by pre-treatment with cAMP (20 min, indicated concentrations) performed in HEK293T cell lysates. Data are from a single experiment representative of at least two independent experiments. **(F)** and **(G)** Gel-ABPP data showing that pre-treatment (1 h) with the indicated concentrations of the HCN pore blocking compounds ivabradine **(F)** and RO-275 **(G)** did not affect the reactivity of WX-01-08 (5 μM, 1 h) with recombinant HCN1 in HEK293T cells. Data are from a single experiment representative of at least two independent experiments. **(H)** Gel-ABPP screen showing effects of pre-treatment with the indicated analogs of WX-02-46 and inactive enantiomer WX-02-26 (20 µM, 2 h) on WX-01-08 (5 μM, 1 h) engagement of recombinant HCN1 in HEK293T cell lysates. Data are from a single independent experiment. **(I)** Quantification of gel-ABPP measuring potency of WX-02-679 versus inactive enantiomer WX-02-678 at blocking WX-01-08 (5 μM, 1 h) engagement of HCN2 (left) and HCN4 (right) in HEK293T cells. HCN2 IC50_WX-02-679_ = 5.1 μM, 95% CI = 2.8 – 6.7 μM, and HCN4 IC50_WX-02-679_ = 3.7 μM, 95% CI = 1.9 – 6.8 μM. Data for each plot are average values ± SD from two independent experiments.

**Figure S6.**
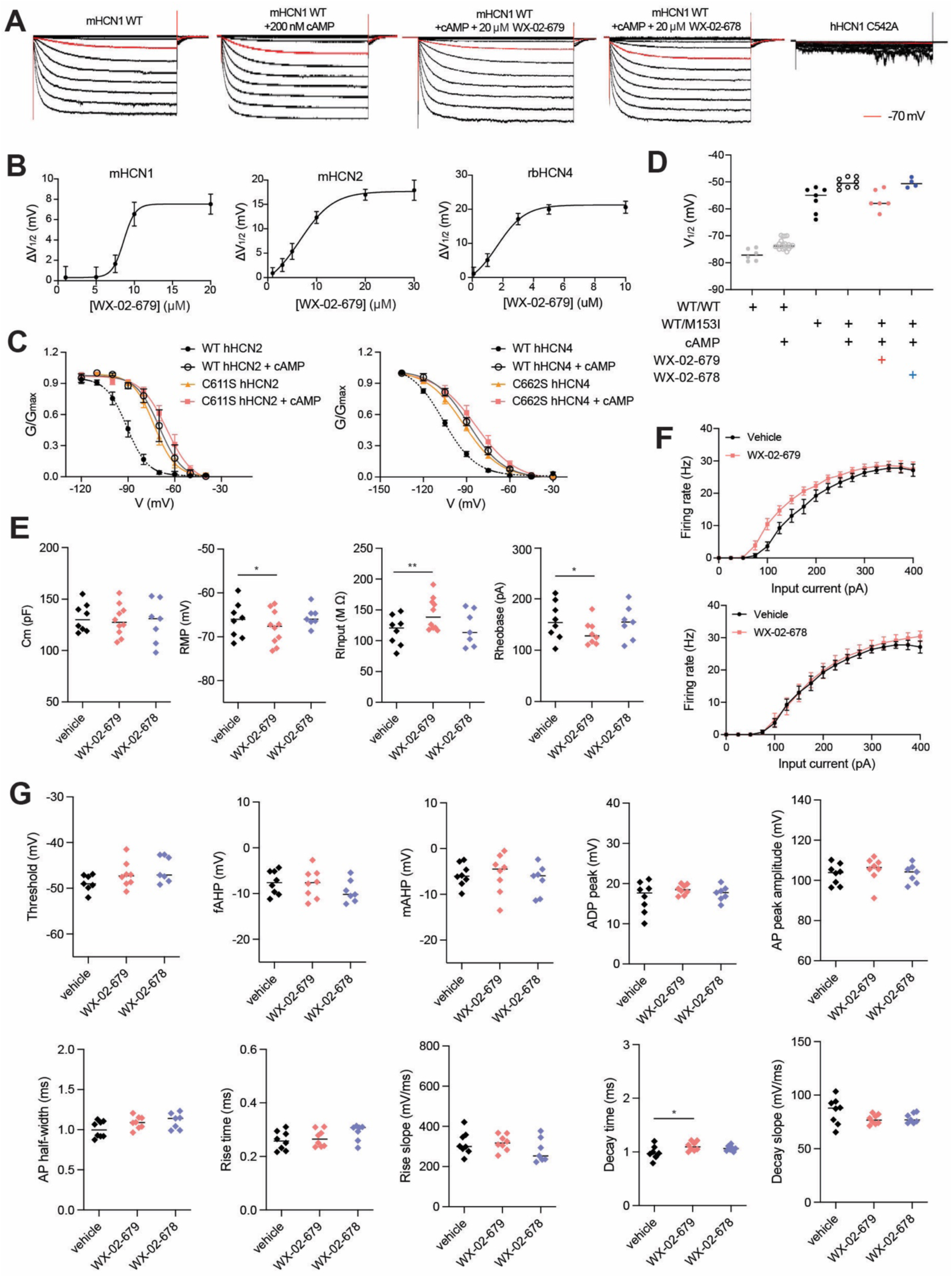
WX-02-679 blocks cAMP-induced changes in HCN channel activation, related to Figure 6. **(A)** Representative whole-cell currents measured in HEK293T recombinantly expressing mouse (m)WT-HCN1 and human (h)C542A-HCN1 after pre-treatment with WX-02-678 or WX-02-679 (10 μM, 20 min) or DMSO followed by addition of cAMP (200 nM) in the patch pipette, showing stereoselective inhibition of cAMP-induced effects on WT-HCN1 by WX-02-679, whereas the C542A-HCN1 mutant exhibits a lack of channel conductance. Red line represents voltage step at -70 mV. **(B)** Concentration-dependent curves for half-activation potentials measured by whole-cell patch clamp recordings in HEK293T cells recombinantly expressing mHCN1 (left), mHCN2 (center) and rbHCN4 (right) after the addition of cAMP (200 nM) through the patch pipette (EC50_mHCN1_ = 8.6 μM, 95% CI = 7.5 – 9.6 μM; EC50_mHCN2_ = 6.5 μM, 95% CI = 5.9 – 9.6 μM; EC50_rbHCN4_ = 1.9 μM, 95% CI = 1.3 – 2.5 μM). Data are average values ± SEM from at least two independent experiments (≥4 cells for each condition per experiment), and fit to the Hill equation plotted as solid lines. **(C)** Mean activation curves measured in HEK293T cells recombinantly expressing hWT- and hC611S HCN2 (left) and hWT- and hC662S-HCN4 with and without addition of cAMP (100 μM) in the patch pipette, showing recombinantly expressed HCN2 and HCN4 cysteine mutants exhibiting cAMP-independent channel conductance. Data are average values ± SEM from at least two independent experiments (≥4 cells for each condition per experiment), and fit to the Boltzmann equation plotted as lines. **(D)** Half-activation potential (V_1/2_) quantified from activation curves from whole-cell patch clamp recordings of HEK293T cells recombinantly expressing WT/WT hHCN1 homomers or WT/M153I hHCN1 heteromers after pre-incubation with WX-02-678 or WX-02-679 (20 μM, 20 min) or DMSO followed by addition of vehicle or cAMP (200 nM) in the patch pipette. Data are average values ± SEM at least two independent experiments (≥4 cells for each condition per experiment). **(E)** and **(F)** Passive membrane properties including cell capacitance (C_m_), resting membrane potential (RMP), input resistance (R_input_), rheobase, and firing rate, respectively, of CA1 neurons upon pre-treatment of mouse hippocampal slices with WX-02-678 or WX-02-679 (20 μM, 5-6 min) or vehicle (DMSO). Data are average values ± SEM (vehicle: N = 8, WX-02-679: N = 10, WX-02-678: N = 7). **(G)** Single action potential properties measured from current clamp recordings of CA1 pyramidal neurons pre-treated with WX-02-678 or WX-02-679 (20 µM, 5-6 min) or vehicle. Data are average values ± SEM (vehicle: N = 8, WX-02-679: N = 10, WX-02-678: N = 7).

## STAR Methods

**Detailed methods are provided in the online version of the paper and include the following:**

### KEY RESOURCE TABLE

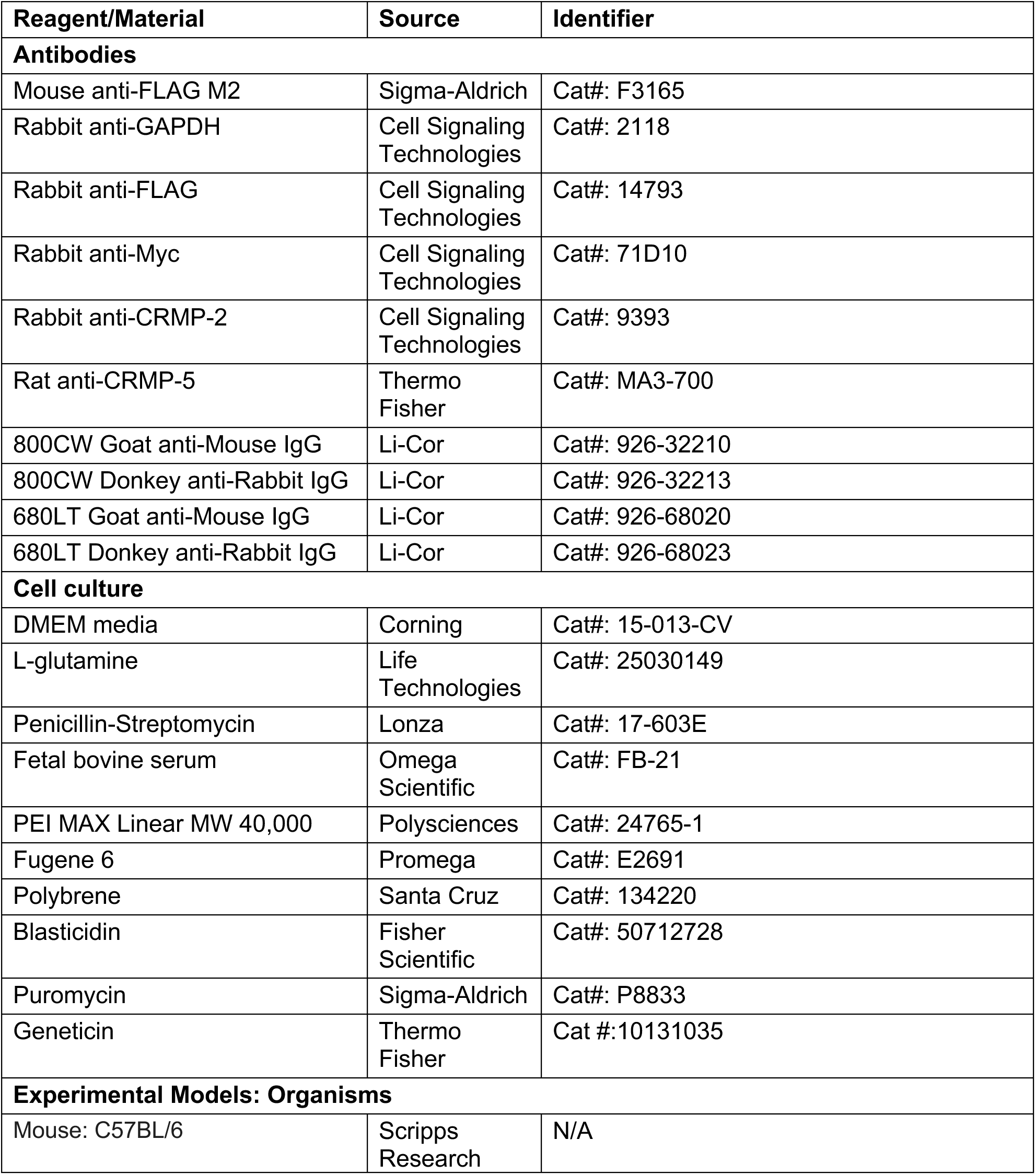
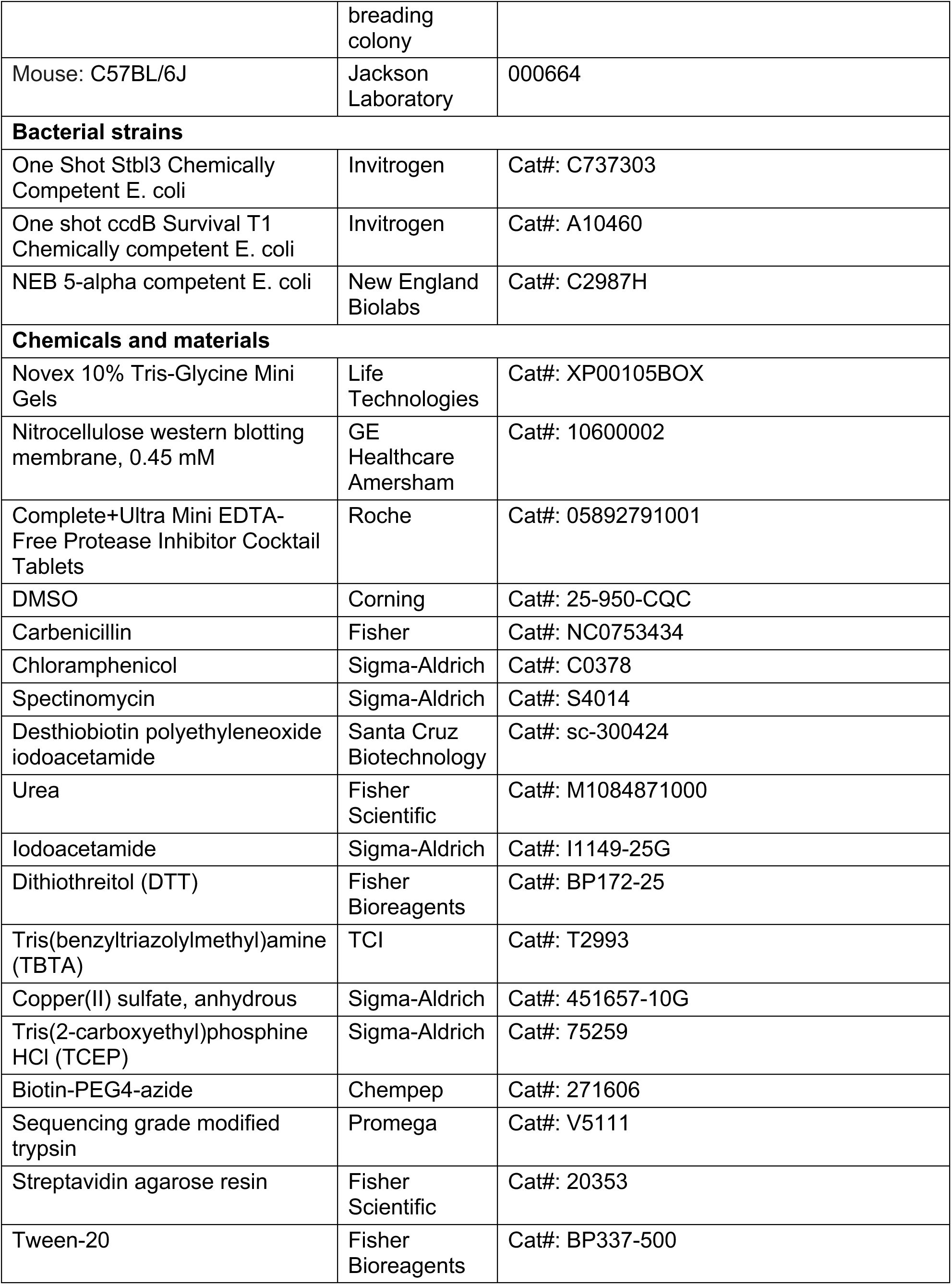
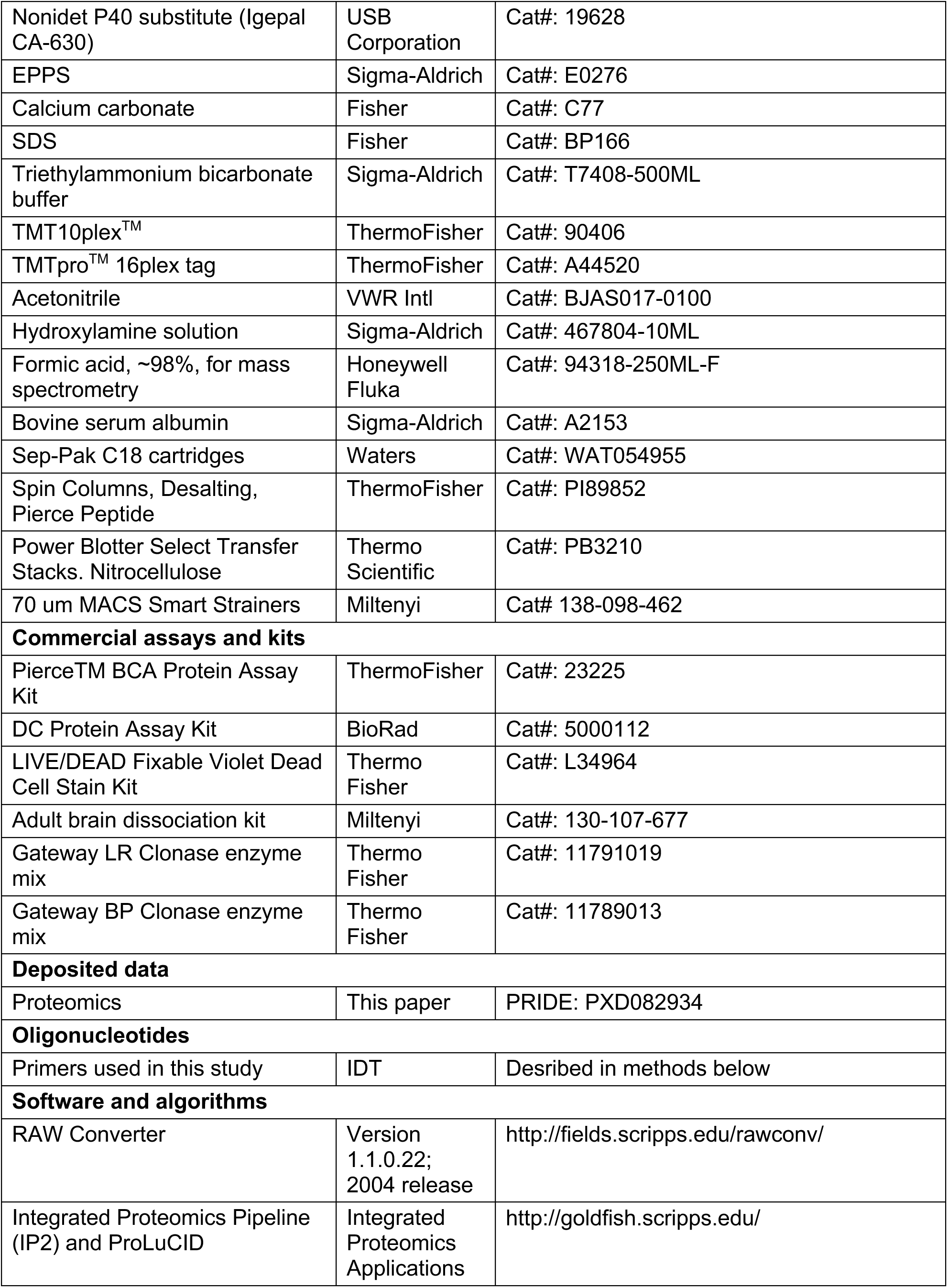
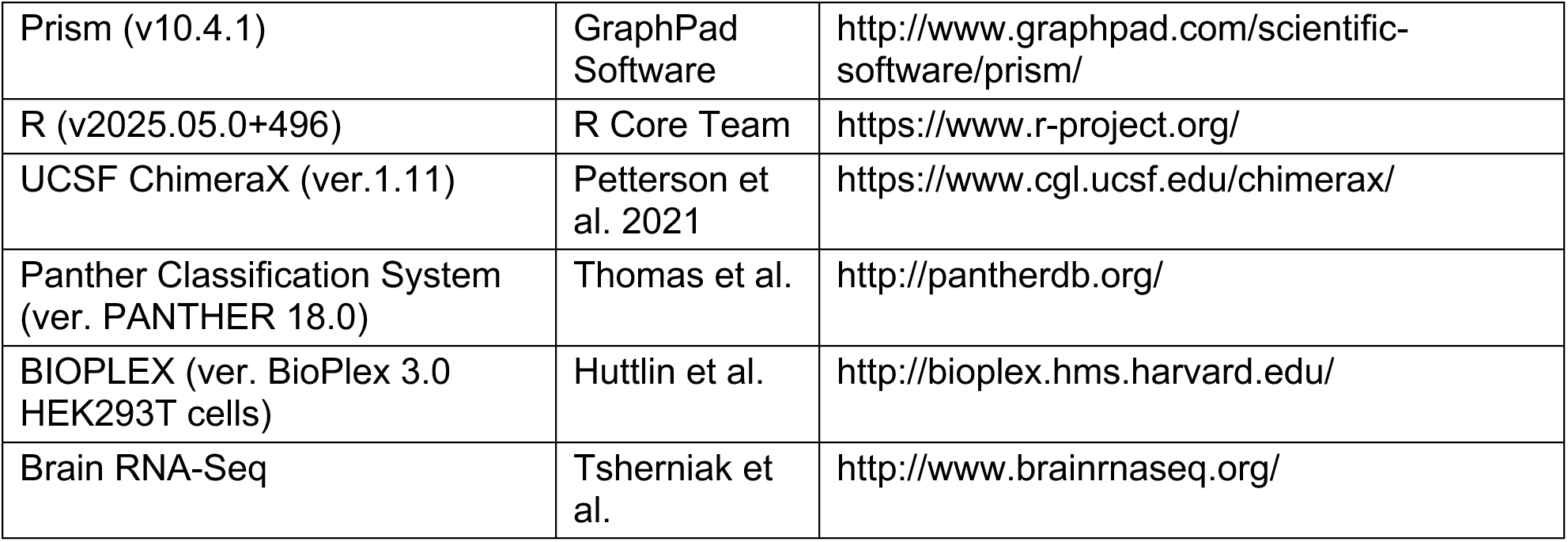

### EXPERIMENTAL MODEL AND STUDY PARTICIPANT DETAILS

#### Research ethics and regulations

All experiments were performed in compliance with protocols approved by The Scripps Research Institute Institutional Review Board.

#### Cell lines and cell culture

All cell lines were obtained from American Type Culture Collection (ATCC). HEK293T (ATCC, CRL-3216), Neuro2a (ATCC, CCL-131), SH-SY5Y (ATCC, CRL-2266) cells were grown Dulbecco’s modified Eagle medium (DMEM), supplemented with 10% fetal bovine serum (FBS), 2 mM L-glutamine, penicillin (100 U/mL), and streptomycin (100 μg/mL), in a humidified, 37 °C/5% CO_2_ tissue culture incubator.

Male 4-8 week old WT C57BL/6 mice were used for all proteomic experiments at Scripps Research, and C57BL/6J mice aged 2-5 months were used for all electrophysiology experiments at Columbia University. All mice were maintained in pathogen-free conditions and handling conformed to requirements of the NIH, The Scripps Research Institute Institutional Animal Care and Use Committee (IACUC) and the Association for Assessment and Accreditation of Laboratory Animal Care (AAALAC). For animals housed at Columbia University, mice were maintained on a 12 hr light–dark cycle (with light turning on at 7 am) under standard housing conditions as above, with ad libitum access to food (Pico Lab rodent diet 5053 for general maintenance or Pico Lab mouse diet 5058 for breeders; Lab Diet, St. Louis, MO) and water. Weanlings were supplemented with DietGel Recovery (ClearH2O, Westbrook, ME) for 2 weeks post-weaning. All animal experiments were conducted in accordance with policies of the NIH Guide for the Care and Use of Laboratory Animals and the Institutional Animal Care and Use Committee (IACUC) of Columbia University.

### METHOD DETAILS

#### Brain dissociation and single cell suspension preparation

Freshly perfused adult C57BL/6 mouse brains were gently homogenized using the gentleMACS Octodissociator and Miltenyi Biotec adult brain dissociation kit (Miltenyi #130-107-677). Briefly, whole brains were added to gentleMACS C tubes (Miltenyi #130-096-334) and cut into 4-8 pieces with scissors. 1950 uL of enzyme 1 mix (50 uL enzyme P and 1.9 mL buffer Z) and 30 uL of enzyme 2 mix (10 uL enzyme A and 20 uL buffer Y) were added to the C tubes, and the samples were run on the Octodissociator with the heater program “37C_ABDK_01” for 30 min at 37 deg. C tubes were detached and centrifuged for 2 min at 200 g to collect material, which were then resuspended and filtered through 70 um MACS SmartStrainer filters (Miltenyi #138-098-462) into 50 mL tubes. 10 mL of cold DPBS were used to wash the C tube and transferred to the filters, which was subsequently washed 3 times with 4 mL DPBS each and discarded. The cell suspension was centrifuged at 1000 g for 10 min at 4 deg and the supernatant was aspirated. For ABPP experiments, the remaining cell pellets were resuspended in DMEM and combined, then seeded into 6-well suspension plates with approximately ½ brain in 2 mL per well for *in cellulo* compound treatment. For flow cytometry analysis, cell suspension was spun at 300 g for 10 min and the pellets were prepared for debris and myelin removal by adding 900 uL of cold debris removal solution followed by gentle overlay with 3100 uL of cold DPBS without mixing phases. The cell suspension was centrifuged for 10 min at 3000 g at 4 deg. As three phases form, the top two phases were aspirated and discarded. Another 5 mL of cold DPBS was added and mixed with the cell suspension by inverting the tube three times. The samples were centrifuged at 4 deg and 1000 g for 10 min and the supernatant was aspirated. Red blood cells were removed by resuspending the cell pellets with 1 mL of diluted RBC removal solution (1:10 with cold ddH2O) and incubating at 4 deg for 10 min. 10 mL cold PB buffer (0.5% BSA in DPBS) was added, and the suspension was centrifuged at 4 deg and 300 g for 10 min. The remaining cell pellets were processed for flow cytometry.

#### Flow cytometry analysis of brainocytes

Cells were stained with LIVE/DEAD fixable violet dead cell stain kit (Thermo Fisher #L34964) at 1 million cells per mL with for 30 min at room temperature, protected from light. 150 uL DPBS were added on top of each sample to a total volume of 200 uL, then washed 2x at 500 g for 5 min. The cells were resuspended and fixed with 200 uL 2% paraformaldehyde in DPBS incubated at 4 deg for 15 min, then centrifuged at 1000 g for 10 min prior to running samples on the flow cytometer. Data were acquired on the NovoCyte Quanteon Agilent analyzer.

#### In situ stereoprobe treatment of brainocytes

Brainocytes were resuspended with DMEM supplemented with pen-strep and glutamine and seeded into 6-well suspension plates with approximately ½ brain in 2 mL per well for in situ compound treatment. Each well was treated with DMSO or 20 μM non-alkyne competitor stereoprobes for 2 h followed by treatment with 5 μM stereochemically matched alkyne probes for 1 h, then the cells were washed two times with chilled DPBS, and immediately processed or stored at −80 °C. For non-competitive protein-directed ABPP, cells were treated with 5 μM alkyne stereoprobe for 1 h.

#### In vitro stereoprobe treatment of brain lysates

Unperfused whole brains from adult C57BL/6 mice were harvested and homogenized using a Dounce homogenizer in cold DPBS. The homogenate was collected and further lysed via probe sonication (2x15, 10% output) and spun down at 3000 g for 5 min to remove debris. The supernatant was collected and normalized to 500 uL 2 mg/mL using a Pierce BCA protein assay kit, and each sample was treated with 5 uM 100x stereoprobe for 1 h at room temperature. Immediately after treatment 60 uL of 1 mg/mL lysate was obtained from each sample and proceeded for gel-ABPP for proteome-wide reactivity as described below.

#### Ex vivo stereoprobe treatment of acute brain slices

Unperfused whole brains from adult C57BL/6 mice were harvested and sliced using a vibratome in oxygenated slicing buffer (2.5 mM KCl, 26 mM NaHCO3, 1 mM NaH2PO4, 0.5 mM CaCl2, 7 mM MgSO4, 11 mM glucose, 228 mM sucrose). Six 200 micron brain slices were seeded in 6-well plates in 2 mL DMEM per well for alkyne stereoprobe treatment for 1 h at 37 deg. Slices were collected and Dounce homogenized in cold DPBS, then further lysed by probe sonication (2x15, 10% output). Lysate was spun down at 3000 g for 5 min to remove debris and the lysate normalized to 500 uL 2 mg/mL for protein-directed ABPP or 60 uL 1 mg/mL for gel-ABPP.

#### Gel-ABPP for proteome-wide reactivity

Following in situ or in vitro stereoprobe treatment and lysis (see earlier sections), lysate samples were normalized to 60 uL of 1 mg/mL using a Pierce BCA protein assay kit. Each sample were treated with 6 μl of click mix (45 µl of 1.7 mM tris((1-benzyl-4-triazolyl)methyl)amine (TBTA) in 4:1 *t*-BuOH:DMSO, 15 µl of 50 mM CuSO_4_ in H_2_O, 15 µl of 1.25 mM rhodamine–polyethylene glycol–azide in DMSO, 15 µl of freshly prepared 50 mM tris(2-carboxyethyl)phosphine in DPBS) for 1 h at room temperature, vortexing every 20 min. The click reaction was quenched by the addition of 22 µl of 4× SDS gel loading buffer. The samples were resolved with 10% SDS– PAGE and imaged by in-gel fluorescent scanning using a BioRad imager with Image Lab software version 6.1.

#### Protein-directed ABPP

Protein-directed ABPP was carried out as previously reported^13^ with slight modifications. Following in situ stereoprobe treatment and lysis (see earlier sections), the total protein content of the whole-cell lysates was measured using a Pierce BCA protein assay kit. The samples were normalized to 500 uL of 2 mg/mL (1 mg of proteome) and treated with 55 μl of click mix (30 µl of 1.7 mM TBTA in 4:1 *t*-BuOH:DMSO, 10 µl of 50 mM CuSO_4_ in H_2_O, 5 µl of 10 mM biotin-PEG4-azide in DMSO, 10 µl of freshly prepared 50 mM tris(2-carboxyethyl)phosphine in DPBS) for 1 h at room temperature with vortexing every 20 min. For in vitro treatments, 500 μl (1 mg) of proteome was first treated with 5 µl of 100× probe for 1 h at room temperature with gentle rotation prior to click reaction. Proteins were precipitated out of solution by the addition of chilled HPLC-grade methanol (600 μl), chloroform (200 μl) and water (100 μl), followed by vigorous vortexing and centrifugation at 16,000*g* for 10 min, to create a disk. Without disrupting the protein disk, both top and bottom layers were aspirated, and the protein disk resuspended in 600 µl of cold methanol, probe sonicated (1x15, 10% output), and centrifuged at 16,000*g* for 10 min. After complete aspiration of the methanol, protein pellets were frozen at −80 °C or immediately resuspended in 500 µl of freshly made 8 M urea in DPBS, followed by the addition of 10 µl of 10% SD. The pellets were probe sonicated to clarity (2x15, 10% output), then reduced with 25 µl of 200 mM dithiothreitol (DTT) at 65 °C for 15 min, followed by alkylation with 25 µl of 400 mM iodoacetamide at 37 °C for 30 min. The samples were quenched with 130 µl of 10% SDS, transferred to a 15-ml tube, and the total volume was brought to 6 ml with DPBS (0.2% final SDS). 100 uL of 50% slurry per sample of washed streptavidin beads (Thermo Fisher #20353) was then added and the probed labelled protein enriched for 1.5 h at room temperature with rotation. After incubation, the beads were pelleted (2 min × 2,000 g) and washed with 0.2% SDS in DPBS (2 × 10 ml), DPBS (1 × 5 ml, then transferred to a protein low-bind Eppendorf safe-lock tube), HPLC grade water (2 × 1 ml) and 200 mM pH 8 4-(2-hydroxyethyl)-1-piperazinepropanesulfonic acid (EPPS; 1 × 1 ml) at room temperature. Enriched proteins were digested on-bead overnight with 200 μl of trypsin mix (2 M urea, 1 mM CaCl_2_, 10 μg/mL trypsin, 200 mM EPPS, pH 8.0).

The beads were spun down, supernatant was collected and 100 μl of acetonitrile (30% final) was added, followed by 6 μl of 20 mg/mL (in dry acetonitrile) of the corresponding TMT^16plex^ tag (for competitive protein-directed ABPP or TMT^10plex^ for non-competitive protein-directed ABPP) for 1.5 h at room temperature with vortexing every 30 min. TMT labelling was quenched by the addition of hydroxylamine (6 μl 5% solution in HPLC grade H_2_O) and incubated for 15 min at room temperature. Samples were then acidified with 20 μl formic acid, combined and SpeedVac-ed to dryness.

Combined samples were desalted with a Sep-Pak column (Waters #WAT054955) then high pH fractionated into ten fractions using peptide desalting spin columns as previously reported^13^ using peptide desalting spin columns (Thermo #89852). Briefly, desalted samples were resuspended in 300 µl of buffer A (95% water, 5% acetonitrile, 0.1% formic acid) by 5 min water bath sonication and bound to the spin columns. Bound peptides were then washed twice with water, once with 5% acetonitrile in 10 mM NH_4_HCO_3_, and eluted into 30 fractions with an increasing gradient of acetonitrile. Every tenth fraction was combined (for example, 1, 10 and 30) and dried with a SpeedVac. Each of the resulting ten fractions was resuspended in buffer A (5% acetonitrile, 0.1% formic acid) and analyzed by MS.

#### Cysteine-directed ABPP

Cysteine-directed ABPP was carried out as previously reported^13^ with slight modifications. Following in situ stereoprobe treatment and lysis (see earlier sections), the total protein content of the whole-cell lysates was measured using a Pierce BCA protein assay kit. The samples were normalized to 500 uL of 2 mg/mL (1 mg of proteome) and treated with 5 μl of 10 mM IA-DTB (in DMSO) for 1 h at room temperature with vortexing every 20 min. Proteins were precipitated out of solution by the addition of chilled HPLC-grade methanol (600 μl), chloroform (200 μl) and water (100 μl), followed by vigorous vortexing and centrifugation at 16,000 g for 10 min, to create a disk. Without disrupting the protein disk, both top and bottom layers were aspirated, and the protein disk washed with 1 ml of cold methanol and centrifuged at 16,000 g for 10 min. The pellets were allowed to air-dry just enough get rid of methanol droplets, then resuspended in 90 μl of denaturing/reducing buffer (9 M urea, 10 mM DTT, 50 mM triethylammonium bicarbonate (TEAB) pH 8.5). The samples were reduced by heating at 65 °C for 20 min, followed by the addition of 10 μl (500 mM) iodoacetamide for 30 min, at 37 °C to alkylate free cysteines. The samples were then centrifuged at 16,000 g or 2 min to pellet any insoluble precipitate and probe-sonicated (1x15, 10% output) and diluted with 300 μl of 50 mM TEAB pH 8.5 to reach a final urea concentration of 2 M. Trypsin (4 μl of 0.25 μg/μl in trypsin resuspension buffer with 25 mM CaCl_2_) was added to each sample and digested at 37 °C overnight with shaking. Digested samples were then diluted with 300 μl of wash buffer (50 mM TEAB pH 8.5, 150 mM NaCl, 0.2% NP-40) containing streptavidin-agarose beads (50 μl of 50% slurry/sample) and were rotated at room temperature for 2 h. The samples were centrifuged (2,000 g, 2 min) and the entire tube (beads and solution) transferred to BioSpin columns and washed (3 × 1 ml wash buffer, 3 × 1 ml DPBS, 3 × 1 ml water). Enriched peptides were eluted from the beads with 300 μl of 50% acetonitrile with 0.1% formic acid and SpeedVac-ed to dryness. Eluted peptides were resuspended in 100 μl 70% EPPS buffer (200 mM, pH 8.0) with 30% acetonitrile, vortexed and water bath-sonicated. The samples were TMT-labelled by the addition of 3 μl of 20 mg/mL (in dry acetonitrile) of corresponding TMT^10plex^ tag, vortexed and incubated at room temperature for 1.5 h with vortexing every 30 min. TMT labelling was quenched with the addition of 3 uL 5% hydroxylamine in H_2_O and incubated for 15 min at room temperature. The samples were then each acidified with 5 μl of formic acid, combined and SpeedVac-ed to dryness. The samples were desalted with Sep-Pak and dried with SpeedVac. The desalted samples were resuspended in 500 μl of buffer A and fractionated with an Agilent HPLC system into a 96-deep-well plate containing 20 μl of 20% formic acid to acidify the eluting peptides, as previously reported.^13^ The peptides were eluted onto a capillary column (ZORBAX 300Extend-C18, 3.5 μm) and separated at a flow rate of 0.5 ml/min using the following gradient: 100% buffer A from 0 min to 2 min, 0–13% buffer B from 2 min to 3 min, 13–42% buffer B from 3 min to 60 min, 42–100% buffer B from 60 min to 61 min, 100% buffer B from 61 min to 65 min, 100–0% buffer B from 65 min to 66 min, 100% buffer A from 66 min to 75 min, 0–13% buffer B from 75 min to 78 min, 13–80% buffer B from 78 min to 80 min, 80% buffer B from 80 min to 85 min, 100% buffer A from 86 min to 91 min, 0–13% buffer B from 91 min to 94 min, 13–80% buffer B from 94 min to 96 min, 80% buffer B from 96 min to 101 min, and 80–0% buffer B from 101 min to 102 min (buffer A, 10 mM aqueous NH_4_HCO_3_; buffer B, acetonitrile). The plates were evaporated to dryness using a SpeedVac and peptides resuspended in 80% acetonitrile, with 0.1% formic acid, and combined to a total of 12 fractions. Samples were dried with a SpeedVac, and the resulting 12 fractions were resuspended in buffer A (5% acetonitrile, 0.1% formic acid in water) and analyzed by MS.

#### TMT LC–MS analysis

The samples were analyzed by LC tandem MS using an Orbitrap Fusion mass spectrometer (Thermo Scientific) coupled to an UltiMate 3000 Series Rapid Separation LC system and autosampler (Thermo Scientific Dionex), as previously reported^13^ and data were acquired with Thermo Scientific Xcalibur software version 2.2. The peptides were eluted onto a capillary column (75-μm-inner-diameter fused silica, packed with C18 (Waters, Acquity BEH C18, 1.7 μm, 25 cm)) or an EASY-Spray HPLC column (Thermo ES902, ES903) using an Acclaim PepMap 100 (Thermo 164535) loading column, and separated at a flow rate of 0.25 μl min^−1^. Data were acquired using an MS3-based TMT method on Orbitrap Fusion or Orbitrap Eclipse Tribrid mass spectrometers. Briefly, the scan sequence began with an MS1 master scan (Orbitrap analysis, resolution 120,000, 400–1,700 *m*/*z*, RF lens 60%, automatic gain control (AGC) target 2E5, maximum injection time 50 ms, centroid mode) with dynamic exclusion enabled (repeat count 1, duration 15 s). The top ten precursors were then selected for MS2/MS3 analysis. MS2 analysis consisted of quadrupole isolation (isolation window 0.7) of precursor ion followed by collision-induced dissociation in the ion trap (AGC 1.8E4, normalized collision energy 35%, maximum injection time 120 ms). Following the acquisition of each MS2 spectrum, synchronous precursor selection enabled the selection of up to 10 MS2 fragment ions for MS3 analysis. MS3 precursors were fragmented by HCD and analyzed using the Orbitrap (collision energy 55%, AGC 1.5E5, maximum injection time 120 ms, resolution 50,000). For MS3 analysis, we used charge state-dependent isolation windows. For charge state *z* = 2, the MS isolation window was set at 1.2; for *z* = 3–6, the MS isolation window was set at 0.7. Raw files were uploaded to the Integrated Proteomics Pipeline (IP2, version 6.0.2) available at http://ip2.scripps.edu/ip2/mainMenu.html, and MS2 and MS3 files were extracted from the raw files using RAW Converter (version 1.1.0.22, available at http://fields.scripps.edu/rawconv/) and searched using the ProLuCID algorithm using a reverse concatenated, non-redundant variant of the Human UniProt database or mouse UniProt database. Cysteine residues were searched with a static modification for carboxyamidomethylation (+57.02146 Da). A dynamic modification for IA-DTB labelling (+398.25292 Da) was included with a maximum number of two differential modifications per peptide. N termini and lysine residues were also searched with a static modification corresponding to the TMT tag (+229.1629 Da for 10plex and +304.2071 Da for 16plex). Peptides were required to be at least six amino acids long. ProLuCID data were filtered through DTASelect (version 2.0) to achieve a peptide false-positive rate below 1%. The MS3-based peptide quantification was performed with reporter ion mass tolerance set to 20 ppm with the Integrated Proteomics Pipeline (IP2).

#### Proteomic data processing

Enrichment ratios (stereoprobe versus stereoprobe) were calculated for each peptide–spectra match by dividing each TMT reporter ion intensity by the sum intensity for all the channels. Peptide–spectra matches were then grouped based on protein ID and (excluding peptides with summed reporter ion intensities < 10,000) coefficient of variation of > 0.5, and < 2 distinct peptides. A protein was considered stereoselectively liganded if the variability corresponding to the probe leading to the highest blockade of IA-DTB did not exceed 20%, and the average IA-DTB blockade by the probe was >33.3% and >2.5-fold that of its enantiomer in at least two biological replicates.

Cysteine engagement ratios (DMSO versus stereoprobe) were calculated for each peptide– spectra match by dividing each TMT reporter ion intensity by the average intensity for the DMSO channels. Peptide–spectra matches were then grouped based on protein ID and residue number, excluding peptides with summed reporter ion intensities for the DMSO channels of 0.5. Replicate channels were grouped across each experiment, and average values were computed for each cysteine site. A variability metric was also computed across replicate channels, which equaled the ratio of the median absolute deviation to the average and was expressed as a percentage. A cysteine site was considered stereoselectively liganded if the variability corresponding to the probe leading to the highest blockade of IA-DTB did not exceed 20%, and the average IA-DTB blockade by the probe was >33.3% and >2.5-fold that of its enantiomer in at least four biological replicates.

#### Generation of CNS enriched protein list

Human bioGPS^40^, mouse bioGPS^40^, and GTEx^41^ (human RNAseq) data were analyzed and aggregated to classify proteins for tissue-enrichment. For each of the three datasets, related samples were grouped into “tissues”, e.g. the amygdala, hippocampus, and spinal cord were all classified as ‘brain’. For each tissue Z-scores were calculated using the maximum signal. All genes that had a maximum tissue Z score > 4 in at least two independent datasets and a maximum absolute signal > 150 for bioGPS and > 10 for GTEx in the brain were classified as “CNS-enriched”.

#### Meta-analysis of stereoselectively liganded CNS-enriched proteins

##### Function class analysis

Panther Classification System (ver. PANTHER 18.0). and KEGG BRITE databases were used to analyze protein functional classes as described previously.^13,129,130^

##### Brain cell type enrichment analysis

Brain RNA-Seq data (Mus musculus) from https://BrainRNASeq.org were analyzed to classify proteins for brain cell type enrichment.^126^ For each brain cell type (newly formed oligodendrocyte, myelinating oligodendrocyte, and OPC were combined under ‘Oligodendrocyte’), Z scores were calculated and genes with a brain cell type Z score of >1.5 were classified as enriched for that brain cell type. Otherwise “non-specific” was used to denote the brain cell type enrichment for that gene.

##### Protein interactome analysis

BIOPLEX (ver. BioPlex 3.0 HEK293T cells) was used to analyze protein interactomes.^131^

#### Cloning and Mutagenesis

All full-length plasmids were obtained from either GenScript in pcDNA3.1-C-(k) DYK (FLAG) or amplified from gBlocks synthesized by Twist Bioscience and cloned into destination vectors (pRK5, pLEX307, etc) using Gateway Cloning (Thermo Fisher #11791019, #11789013). Site-directed mutagenesis was carried out using a Q5 site-directed mutagenesis kit (New England BioLabs #E0554S), using the primers shown below. All plasmids were amplified in One Shot Stbl3 chemically competent E. coli (Invitrogen #C737303) or NEB 5-alpha competent E. coli cells (New England BioLabs #C2987H) using vendor protocols and purified by miniprep (Zymo Research #D4019), except for the Gateway destination vectors containing ccdB, which were amplified in One Shot ccdB Survival T1 cells (Invitrogen #A10460).

DPYSL2_C504A_fwd: 5′-GACGCTTATGAGAAGTGCCG - 3’

DPYSL2_C504A_rev: 5’-TCATACAGGCCACGAGGA - 3’

PLP1_C6A_fwd: 5’-CTTGTTAGAGgccTGTGCAAGATGTC - 3’

PLP1_C6A_rev: 5’-CCCATTAAGCCTGCTTTTTTG - 3’

PLP1_C7A_fwd: 5’-GTTAGAGTGCgccGCAAGATGTCTGGTAG - 3’

PLP1_C7A_rev: 5’-AAGCCCATTAAGCCTGCT - 3’

PLP1_C6_7A_fwd: 5’-CTTGTTAGAGgccgccGCAAGATGTCTGGTAGG - 3’

PLP1_C6_7A_rev: 5’-CCCATTAAGCCTGCTTTTTTG - 3’

PDE7B_C136A_fwd: 5’-AACACTGTTGgccCACCTCTTCAATAC - 3’

PDE7B_C136A_rev: 5’-ACCAGGCTGTTTCCATTTG - 3’

HCN1_C542A_fwd: 5’-TGGAGAGATTgccCTGCTGACCAAAG - 3’

HCN1_C542A_rev: 5’-AAGTAAGAGCCATCTGTC - 3’

HCN1_C542S_fwd: 5’-TGGAGAGATTagcCTGCTGACCA - 3’

HCN1_C542S_rev: 5’-AAGTAAGAGCCATCTGTCAGC - 3’

HCN2_C611A_fwd: 5’-CGGGGAGATCgccCTGCTCACCC - 3’

HCN2_C611A_rev: 5’-AAGTAGGAGCCATCGGAC - 3’

HCN4_C662A_fwd: 5’-TGGAGAGATCgccCTGCTGACCCGGG - 3’

HCN4_C662A_rev: 5’-AAGTAGGAGCCGTCGGCC - 3’

#### Generation of DPYSL2/DPYSL5, PDE7B, HCN1 stable cell lines

Full-length DPYSL2, DPYSL5, PDE7B, or HCN1 (WT and C-to-A/S mutants) carrying C-terminal epitope tags (FLAG or HA) were generated using Gateway cloning into lentiviral vectors using primers shown in ‘Cloning and Mutagenesis’ section. To generate lentivirus, 3.5 × 10^5^ HEK293T cells were seeded in 6-well plates overnight, in 2 mL of DMEM. 1000 ng protein-encoding Lentiviral vector, 1000 ng lentiviral packaging vector psPAX2, and 100 ng envelope vector VSV-G were mixed in Optimem, and 10 uL of PEI (1 mg/mL, Polysciences) was added. The DNA:PEI complex was incubated at room temperature for 20 min and the complex added dropwise to the HEK293T cells. Media were replaced with fresh DMEM with 30% FBS plus pen-strep and 2 mM glutamine, 9 h post transfection. The virus was collected at 48 h post transfection and filtered through a 0.45-μm syringe filter to eliminate floating cells. For transduction, 0.1-0.5 million cells (HCT116, SH-SY5Y, Neuro2a) were mixed with 100-500 µL of viral supernatant in 1-2 mL supplemented with 6-8 μg/mL polybrene. Cells were spin-infected at 930 × g at 30 °C for 1 h and incubated for 24 h at 37 °C. Selection was initiated with 2 μg/mL puromycin or 1 mg/mL geneticin for 1-2 weeks (sequential transduction and selection with both antibiotics for stable cell lines expressing both DPYSL2 and DPYSL5). Selected pools were validated by immunoblots (FLAG or HA).

#### Generation of sgDPYSL5 CRISPR/Cas9 knockout cells

Stable knockout cells were generated by the transduction of cells with LentiCRISPR v2-Blast carrying sgDPYSL5. Briefly, sgRNA (sgDPYSL5-03_sense5′-GACGCTTATGAGAAGTGCCG-3’) were annealed and cloned into LentiCRISPR-v2 Blast. Viral supernatants were generated as described above, and parental SH-SY5Y cells or SH-SY5Y cells stably expressing DPYSL2-FLAG were transduced and selected with 5 μg/mL of blasticidin for one week. For parental SH-SY5Y cells, remaining DPYSL5 were further knocked out using sgRNA (sgDPYSL5-04_sense5′-GCACGCTTGCAAGGACATTG-3’) cloned into LentiCRISPR-v2 Puro. The selected stable pooled cell population were evaluated by immunoblot with anti-DPYSL5 antibody.

#### Gel-ABPP of transiently overexpressed recombinant proteins

HEK293T cells (2.5 × 10^5^) were seeded in TC-treated six-well plates overnight and transfected with 1 µg of FLAG-epitope tag plasmids in 250 Optimem (Thermo Fisher #31-985-062) with polyethylenimine (PEI) at a ratio of 1:3 (ug DNA:uL PEI) for 48 h, with a media exchange at 24 h. The cells were treated with alkyne probe only for 1 h or first pre-treated with competitor probe for 2 h followed by alkyne probe for a further 1 h. The cells were then collected, lysed, clicked with rhodamine azide, analyzed via in-gel fluorescent scanning as described above. Following in-gel fluorescence, gels were transferred to nitrocellulose at 60 V for 120 min and proceeded to standard western blotting procedures, described in the ‘Western blotting and antibodies’ section.

#### Gel-ABPP of recombinant proteins in stable cell lines

Parental and engineered cells stably expressing WT or cysteine mutant recombinant proteins (2.5 × 10^5^) were seeded in TC-treated six-well plates and cultured for 24-48 h. The cells were then treated with alkyne probe only for 1 h or first pre-treated with competitor probe for 2 h followed by alkyne probe for a further 1 h. The cells were collected, lysed, clicked with rhodamine azide, analyzed via in-gel fluorescent scanning as described above. Following in-gel fluorescence, gels were transferred and continued for standard western blotting procedures, described in the ‘Western blotting and antibodies’ section.

#### IP-gel ABPP of recombinant proteins in transient and stable cell lines

Parental and cells transiently or stably expressing WT or cysteine mutant recombinant proteins (2.5 × 10^5^) were seeded in TC-treated six-well plates and cultured for 24-48 h. The cells were then treated with alkyne probe only for 1 h or first pre-treated with competitor probe for 2 h followed by alkyne probe for a further 1 h. Treated cells were collected and washed three times with chilled DPBS. The cell pellets were resuspended in 250 μL of IP lysis buffer (1% NP-40 in DPBS), and cell pellets were lysed by sonication (2 × 15 pulses, 10% power output). The supernatant was assayed for total protein using BCA protein quantification kit. Protein concentrations for all the samples were adjusted to 1 mg/mL, 30 µL of proteome was taken for the input, and 10 µL of 4× SDS gel loading buffer was added. The remaining lysate (200-500 µg) was mixed with 16 μL of prewashed anti-FLAG or anti-Myc magnetic beads at 4 °C with rotation for 1-2 h or overnight, respectively. The samples were washed three times with IP wash buffer (0.2% NP-40 in DPBS) and once with DPBS. Beads were resuspended in 30 µL of DPBS for on-bead click reaction with 3 μL of click mix (1.5 μl of 1.7 mM tris((1-benzyl-4-triazolyl)methyl) amine (TBTA) in 4:1 t-BuOH:DMSO, 0.5 μL of 50 mM CuSO4 in H2O, 0.5 μL of 1.25 mM rhodamine–polyethylene glycol–azide in DMSO, 0.5 μL of freshly prepared 50 mM tris(2-carboxyethyl)phosphine in DPBS) for 1 h at room temperature with flicking of the tube to mix the bead mixture every 15 min. The click reaction was quenched and proteins were eluted off the beads by the addition of 11 μL of 4X SDS gel loading buffer and vortexing. The supernatant was collected with a magnetic stand into new tubes and input and IP samples were resolved on SDS–PAGE gels and imaged as described above.

#### Western blotting and antibodies

For protein expression analysis, cells pellets were lysed by probe sonication into ice cold DPBS (Gibco #14190144) supplemented with one protease inhibitor tablet per 10 mL (Roche #4693159001). Cell lysate concentrations were determined using DC Protein Assay (BioRad #5000112) and standardized to 1 mg/mL before the addition of 4x SDS loading dye. Western blots were performed using either pre-cast Novex™ WedgeWell™ 10%, Tris-Glycine, 1.0 mm, Mini Protein Gel (Invitrogen #XP00100PK2) or hand-poured 10% acrylamide gels and ran using tris-glycine running buffer (25 mM Tris pH 8.6, 192 mM glycine, 0.1% SDS) at 170 V for 1 hour (pre-cast) or 275 V for 3.5 hours (hand poured). Gels were transferred to nitrocellulose membranes in Towbins transfer buffer (25 mM Tris, 192 mM glycine, 20% MeOH) at 60 V for 2 h. The transferred blots were blocked in milk (5% w/v in TBST (20 mM Tris pH 7.5, 150 mM NaCl, 0.1% Tween-20)) for 1 h at room temperature. Primary antibodies were diluted into 5% w/v milk or BSA in TBST and incubated overnight at 4°C with gentle rotation. Blots were washed in TBST for 5 minutes 3x followed by the addition of secondary antibodies diluted into 5% milk in TBST for 1 hour at room temperature (dilution: 1:10,000). Blots were imaged using an Li-Cor Odyssey IR imager and quantitated using ImageStudio Lite software. Primary antibodies used in this study: mouse anti-FLAG M2 antibody (Sigma #F3165, 1:1000), rabbit anti-GAPDH (CST #2118, 1:1000), rabbit anti-FLAG antibody (CST #14793, 1:1000), rabbit anti-Myc antibody (CST #71D10, 1:1000), rabbit CRMP-2 (DPYSL2) antibody (CST #9393, 1:1000), rat anti-CRMP-5 (DPYSL5) antibody (Thermo Fisher #MA3-700, 1:1000). Secondary IRDye antibodies used in this study: 800CW Goat anti-Mouse IgG (#926-32210), 800CW Donkey anti-Rabbit IgG (#926-32213), 680LT Goat anti-Mouse IgG (#926-68020) and 680LT Donkey anti-Rabbit IgG (#926-68023) were purchased from Li-Cor.

#### Immunoprecipitation-mass spectrometry (IP-MS) experiments in stable cell lines

Cells stably expressing FLAG epitope-tagged protein of interest (WT and cysteine mutants of DPYSL2-FLAG or PDE7B-FLAG) were treated with DMSO or stereoprobes for 3 h. The cells were collected and washed twice with cold DPBS then resuspended in 500 uL of IP lysis buffer (50 mM EPPS pH 8.0, 150 mM NaCl, 1% NP-40, 10% glycerol) supplemented with EDTA-free complete protease inhibitor. Lysing was achieved by rotating the samples in IP lysis buffer at 4 deg for 1 h. The lysate was clarified by spinning at 16,000 × g for 5 min and the total protein content of the supernatant was quantified and normalized using Pierce BCA assay kit. Lysate (1-2 mg) was mixed with 40 μL of prewashed anti-FLAG magnetic beads (Thermo Fisher #A26797) for 2-4 h at 4 °C with rotation. The samples were washed three times with IP wash buffer (25 mM EPPS pH 8.0, 150 mM NaCl, 0.2% NP-40) and once with 50 mM EPPS pH 8.0. Enriched proteins were eluted off the beads by boiling with 40 μL of 8 M urea in 50 mM EPPS pH 8.0 at 65 °C for 10 min, and the supernatant was collected with a magnetic stand into new tubes. Sample was then reduced with 2 μL of 200 mM DTT for 15 min at 65 °C and alkylated with 2 μL of 400 mM iodoacetamide for 30 min at 37 °C. Samples were further diluted to 2 M urea with 115 μL of 50 mM EPPS pH 8.0 and trypsin (4 μL of 0.25 μg/µL in 50mM EPPS with 25 mM CaCl2) was added to each sample and digested at 37 °C overnight. Acetonitrile (75 μL) was added to each sample and TMT labeled and desalted as described in [Protein-directed ABPP] section. Combined samples were high-pH spin column fractionated and combined into three fractions.

#### PDE7B exogenous substrate enzyme assay

PDE7B enzymatic activity was assayed using a modified version of previously published LC/MS method for quantifying cAMP in cells and lysates.^132^ HEK293T cells expressing recombinant WT and C136A PDE7B were treated in situ with stereoprobes for 2h. After treatment and collection, the cells were washed 2x with cold DPBS then resuspended with enzyme assay buffer (50 mM pH 8 EPPS, 10 mM MgCL2, 100 mM NaCl) and normalized using the Pierce BCA assay kit. cAMP substrates diluted in enzyme assay buffer were added into Eppendorf tubes to create a total reaction volume of 100 uL 0.2 mg/mL lysate with 10 uM cAMP. The enzymatic reaction was allowed to incubate for 30 min at 37 deg and quenched by the addition of 300 uL ice cold MeOH containing 1 uM internal standard 13C5-cAMP. To precipitate proteins, the quenched samples were vortexed vigorously then chilled at -80 for 1-2 h. The samples were spun at 16,000 g for 10 min to pellet precipitated proteins and the supernatant were transferred to autosampler vials for metabolomic analysis. For in vitro compound treatment, 200 uL of 1 mg/mL lysate of HEK293T cells overexpressing WT or cysteine mutant PDE7B were treated with DMSO, stereoprobes, or PDE inhibitors IBMX and BRL-50481 for 2 h before the enzyme assay.

cAMP was measured using LC-MS/MS. Samples were injected into a Luna® Omega Polar (2.1 × 50 mm, 1.6 µm) C18 column used with a mobile phase consisting of 0.1% FA in HPLC H2O at a flow rate of 0.35 mL/ min. The injection volume was 5 µL. Eluted metabolites were detected using a triple quad mass spectrometer (Agilent 6470 MassHunter, Agilent) via multiple reaction monitoring (MRM) using an electrospray ionization (ESI) source in negative mode, with the following parameters: gas temperature: [350 °C]; gas flow: [11 L/min]; nebulizer: [45 psi]; sheath gas temperature: [450 °C]; sheath gas flow: [12 L/min]; capillary: [12 V]; nozzle voltage/charging: [1500 V]. The MRM transitions used were: AMP: 346.2 → 134.1; cAMP-^13^C_5_ (internal standard) m/z 333.2 → 134.1; cAMP: m/z 328.1 → 134.1. Metabolites were quantified by MassHunter quantitative analysis (version 10.0, Agilent) by integrating their peak area and normalizing relative to the peak area of the internal standard, 13C5-cAMP.

#### In vitro cAMP competition assay

HEK293T cells transiently overexpressing WT and cysteine mutants of HCN channels were collected and lysed in DPBS via probe sonication (2x15, 10% output). The lysate was normalized to 2 mg/mL using Pierce BCA assay kit, and added to 2x cAMP stock solution in DPBS to form a final concentration of 1 mg/mL and incubated for 30 min. Then, 100x stereoprobe stock solution was added to each sample and incubated at room temperature with rotation for 1 h. Immediately upon the end of compound treatment, 10% NP40 in DPBS was added to each sample to a final concentration of 1%, and mixed with 20 μL of prewashed anti-FLAG magnetic beads for 1 h at 4 °C with rotation. The samples were washed three times with IP wash buffer (0.2% NP-40 in DPBS) and once with DPBS. The beads were resuspended with 60 uL DPBS and treated on bead with 6 uL click mix, quenched, and prepared for gel-ABPP as previously described

#### Whole cell patch clamp electrophysiology in HEK293T cells

Whole cell patch clamp experiments were performed as previously described.^75,88,92^ Briefly, 24 h after transfection HEK293T cells were dispersed and single GFP+ cells were selected for patch-clamp recordings. Each set of experiments contains data from controls and mutants measured on the same day. Currents were recorded in whole-cell configuration at room temperature, either with an ePatch amplifier (Elements, Cesena, Italy) or with an Axopatch 200b amplifier (Molecular Devices); data acquired with the Axopatch 200b amplifier were digitized with an Axon Digidata 1550B (Molecular Devices) converter. Signals were acquired with a sampling rate of 5 kHz and low pass filtered at 2.5 kHz. Data analysis was performed using Clampfit 10.7 (Molecular devices). Patch pipettes were pulled from 1.5 mm O.D. and 0.86 mm I.D. borosilicate glass capillaries (Sutter, Novato, CA) and had resistances ranging from 3 to 6 MΩ. For HCN1 channels recordings, patch pipettes were filled with a solution containing 10 mM NaCl, 130 mM KCl, 1 mM egtazic acid (EGTA), 0.5 mM MgCl2, 2 mM ATP (magnesium salt), and 5 mM HEPES–KOH buffer (pH 7.2), while the extracellular bath solution contained 110 mM NaCl, 30 mM KCl, 1.8 mM CaCl2, 0.5 mM MgCl2, and 5 mM HEPES–KOH buffer (pH 7.4).

Where indicated, cAMP and WX-02-678/679 was added to the pipette solution to reach indicated final concentrations. Controls were treated with the vehicle (DMSO). To assess HCN1 and HCN2 channel activation curves, the holding potential was -20 mV (1 s), with steps from -30 mV to -120 mV (-10 mV 26 increments, 3.5 s) and tail currents recorded at -40 mV (3.5 s). For HCN4 the step protocol was extended to -140 mV and the step duration was 5s

Only cells in which a 1 GΩ seal or better was achieved were kept for analysis. For I/V plots currents were normalized to cell capacitance, indicated as ISS (pA/pF). Neither series resistance compensation nor leak correction were applied. Mean activation curves were obtained by fitting maximal tail current amplitude, plotted against the preconditioning voltage step, with the Boltzmann equation: y= 1/[1+exp((V−V1/2)/k)], where V is voltage, y the fractional activation, V1/2 the half-activation voltage, and k the inverse slope factor in mV (k = -RT/zF). Mean activation curves were obtained by fitting individual curves from each cell to the Boltzmann equation and then averaging all the obtained values. Data were analyzed with Clampfit (Molecular devices) and Origin (OriginLab) softwares and are presented as mean ± SEM. Statistical analysis was performed with the Student’s t-test for unpaired data or One-way ANOVA with Fisher’s test, as indicated for each experiment. Significance level was set to p = 0.05.

#### Inside-out patch clamp electrophysiology in HEK293T cells recombinantly expressing rbHCN4

The cells were grown as previously described.^92^ Briefly, HEK-293T cells were transfected with 1 ug rbHCN4 and 0.3 ug GFP-containing plasmid. After 24 h cells were dispersed and single GFP^+^ cells were selected for patch-clamp experiments at room temperature. HCN4 currents were recorded using an ePatch amplifier and PULSE acquisition software (E-Zpatch, Elements, Cesena, Italy). Patch pipettes had a resistance of approximately 1 MΩ after fire polishing. The pipette (external) solution contained (in mM): 70 KCl, 70 NaCl, 1 MgCl₂, 1.8 CaCl₂, 1 BaCl₂, 2 MnCl₂, and 5 HEPES, adjusted to pH 7.4 with KOH. The bath (internal) solution contained (in mM): 130 KCl, 10 NaCl, 2 CaCl₂, 10 HEPES, and 5 EGTA, adjusted to pH 7.2.

Whole-cell configuration was first established, after which the pipette was rapidly withdrawn to excise the membrane patch and obtain the inside-out configuration.^133^ The preparation was allowed to stabilize until the rundown of HCN4 currents had subsided, then record the HCN4 currents. Voltage-clamp protocols were initiated from a holding potential of −20 mV. Hyperpolarizing voltage steps from −60 to −175 mV (–15 mV increments, 5 s), followed by a 1-s step to −120 mV to record tail currents and a subsequent 6-s return to the holding potential of −20 mV.

WX-02-679 was dissolved in DMSO to prepare a 10 mM stock solution, whereas cAMP was dissolved in Milli-Q water to prepare a 2 mM stock solution. Following acquisition of baseline currents, cAMP was added to the bath solution to a final concentration of 100 μM. Currents were recorded again approximately 2 min after cAMP application. WX-02-679 was then added to the bath solution to a final concentration of 20 μM, and currents were recorded again approximately 2 min after drug application. The effect of WX-02-679 on HCN4 currents was evaluated by comparing the current amplitude at −105 mV under the respective experimental conditions. Current traces were plotted using Origin 2016, and data were analyzed using Clampfit 10.4 and Origin 2016.

#### Whole cell patch clamp electrophysiology in brain slices

Hippocampal slices were obtained from C57BL/6J mice aged 2-5 months as previously described.^76^ Recordings were performed in current clamp mode using a Multiclamp 700B amplifier and digitized using a Digidata 1322A A/D interface (Molecular Devices, CA), at a sampling rate of 10 or 50 kHz (low pass filtered at 10 kHz). Healthy somas of CA1 pyramidal neurons were identified visually and patched using borosilicate glass pipettes (I.D. 0.75 mm, O.D. 1.5 mm, Sutter Instruments, UK) with a tip resistance of 4 - 6 MΩ. The pipette intracellular solution contains: 125 mM K-gluconate, 10 mM phosphocreatine (di-tris), 1.5 mM NaCl, 3 mM KCl, 10 mM HEPES, 0.1 mM EGTA, 5 mM ATP (magnesium salt), 0.4 mM Na_3_-GTP, adjusted to a pH of 7.25 with KOH. The extracellular solution (ACSF) contains: 22.5 mM glucose, 125 mM NaCl, 1 mM MgCl_2_, 2 mM CaCl_2_, 25 mM NaHCO_3_, 2.5 mM KCl, 1.25 mM NaH_2_PO_4_, 3 mM Na-pyruvate, 1 mM ascorbic acid, pH 7.2. Recordings were kept for analysis only if the series resistance after establishing the whole-cell configuration did not exceed 15 MΩ and did not change by more than 20% of the initial value during the course of the experiment.

WX-02-678 and W-02-679 compounds were dissolved in DMSO to make 10 mM stock solutions and then diluted in ACSF to the desired concentration. Slices were incubated with oxygenated ACSF at 33-34°C containing the compounds for 5-6 minutes prior to recording and then perfused with clean ACSF throughout the patch clamp session. Controls were vehicle (DMSO)-treated.

CA1 pyramidal neurons were patched at the soma in whole-cell current-clamp mode. Resting membrane potential (RMP) was measured during 10 sec of gap free recording; for all other measurements neurons were held at - 70 mV by bias current injection. Input-output curves were obtained using current steps from 0 to + 400 pA (25 pA increments, 1 sec), while input resistance (R_input_) was calculated from voltage responses to -20 and -40 pA current injections. Rheobase and action potential (AP) properties were determined from the first AP elicited by a 1 sec ramp from 0 to +1000 pA. AP threshold was defined using the first derivative of the voltage trace (50x sampling interval). Fast and medium afterhyperpolarization (fAHP and mAHP) and afterdepolarization peak (ADP) were quantified relative to AP threshold or baseline, respectively. Voltage sag (V_sag_) ratio was calculated as (V_peak_-V_steady state_)/V_peak_ following a hyperpolarizing current injection adjusted to reach -105/-110 mV in each cell. Rebound depolarization was measured as the peak voltage following termination of the hyperpolarizing pulse, relative to baseline.

Data were analyzed using Clampfit (Molecular devices), MATLAB® (Mathworks, MA, USA) and Origin (OriginLab) softwares and are presented as mean ± SEM. Statistical analysis was performed with One way ANOVA with Fisher’s test. Significance level was set to p = 0.05.

## References

1. Wang, J.Y., and Doudna, J.A. (2023). CRISPR technology: A decade of genome editing is only the beginning. Science 379, eadd8643. 10.1126/science.add8643.

2. Shendure, J., Balasubramanian, S., Church, G.M., Gilbert, W., Rogers, J., Schloss, J.A., and Waterston, R.H. (2017). DNA sequencing at 40: past, present and future. Nature 550, 345–353. 10.1038/nature24286.

3. Dang, C.V., Reddy, E.P., Shokat, K.M., and Soucek, L. (2017). Drugging the “undruggable” cancer targets. Nat Rev Cancer 17, 502–508. 10.1038/nrc.2017.36.

4. Spradlin, J.N., Zhang, E., and Nomura, D.K. (2021). Reimagining Druggability Using Chemoproteomic Platforms. Accounts of Chemical Research. 10.1021/acs.accounts.1c00065.

5. Niphakis, M.J., and Cravatt, B.F. (2024). Ligand Discovery by Activity-Based Protein Profiling. Cell Chem Biol 31, 1636–1651. 10.1016/j.chembiol.2024.08.006.

6. Denis, J.D.S., Hall, R.J., Murray, C.W., Heightman, T.D., and Rees, D.C. (2021). Fragment-based drug discovery: opportunities for organic synthesis. RSC Med. Chem. 12, 321–329. 10.1039/D0MD00375A.

7. Edfeldt, F.N.B., Folmer, R.H.A., and Breeze, A.L. (2011). Fragment screening to predict druggability (ligandability) and lead discovery success. Drug Discovery Today 16, 284–287. 10.1016/j.drudis.2011.02.002.

8. Kolb, P., Ferreira, R.S., Irwin, J.J., and Shoichet, B.K. (2009). Docking and chemoinformatic screens for new ligands and targets. Current Opinion in Biotechnology 20, 429–436. 10.1016/j.copbio.2009.08.003.

9. Prudent, R., Annis, D.A., Dandliker, P.J., Ortholand, J.-Y., and Roche, D. (2021). Exploring new targets and chemical space with affinity selection-mass spectrometry. Nat Rev Chem 5, 62–71. 10.1038/s41570-020-00229-2.

10. Wen, X., Wu, X., Jin, R., and Lu, X. (2023). Privileged heterocycles for DNA-encoded library design and hit-to-lead optimization. European Journal of Medicinal Chemistry 248, 115079. 10.1016/j.ejmech.2022.115079.

11. Won, S.J., Zhang, Y., Reinhardt, C.J., Hargis, L.M., MacRae, N.S., DeMeester, K.E., Njomen, E., Remsberg, J.R., Melillo, B., Cravatt, B.F., et al. (2024). Redirecting the pioneering function of FOXA1 with covalent small molecules. Molecular Cell 84, 4125–4141.e10. 10.1016/j.molcel.2024.09.024.

12. Kathman, S.G., Koo, S.J., Lindsey, G.L., Her, H.-L., Blue, S.M., Li, H., Jaensch, S., Remsberg, J.R., Ahn, K., Yeo, G.W., et al. (2023). Remodeling oncogenic transcriptomes by small molecules targeting NONO. Nat Chem Biol 19, 825–836. 10.1038/s41589-023-01270-0.

13. Njomen, E., Hayward, R.E., DeMeester, K.E., Ogasawara, D., Dix, M.M., Nguyen, T., Ashby, P., Simon, G.M., Schreiber, S.L., Melillo, B., et al. (2024). Multi-tiered chemical proteomic maps of tryptoline acrylamide-protein interactions in cancer cells. Nat Chem 16, 1592–1604. 10.1038/s41557-024-01601-1.

14. Hayward, R.E., Berkeley, R.F., Gao, Z., Garhammer, M., Morizono, M.A., Njomen, E., Li, H., DeMeester, K.E., Cociorva, V., Herzik, M.A., et al. (2025). Tryptoline Stereoprobe Elaboration Identifies Inhibitors of the GRPEL1-HSPA9 Chaperone Complex. Preprint at bioRxiv, 10.1101/2025.10.20.683548 https://doi.org/10.1101/2025.10.20.683548.

15. Goetzke, F.W., Bernard, S.M., Ju, C.-W., Pollock, J., DeMeester, K.E., Gross, J., Simon, G.M., He, C., Melillo, B., and Cravatt, B.F. (2026). Complexoform-restricted covalent TRMT112 ligands that allosterically agonize METTL5. Nat Chem Biol 22, 770–782. 10.1038/s41589-025-02099-5.

16. Niessen, S., Dix, M.M., Barbas, S., Potter, Z.E., Lu, S., Brodsky, O., Planken, S., Behenna, D., Almaden, C., Gajiwala, K.S., et al. (2017). Proteome-wide Map of Targets of T790M-EGFR-Directed Covalent Inhibitors. Cell Chem Biol 24, 1388–1400.e7. 10.1016/j.chembiol.2017.08.017.

17. Jiang, M., and van der Stelt, M. (2018). Activity-Based Protein Profiling Delivers Selective Drug Candidate ABX-1431, a Monoacylglycerol Lipase Inhibitor, To Control Lipid Metabolism in Neurological Disorders. J. Med. Chem. 61, 9059–9061. 10.1021/acs.jmedchem.8b01405.

18. Mitchell, D.C., Menon, A., and Garner, A.L. (2019). Chemoproteomic Profiling Uncovers CDK4-Mediated Phosphorylation of the Translational Suppressor 4E-BP1. Cell Chemical Biology 26, 980–990.e8. 10.1016/j.chembiol.2019.03.012.

19. Capes-Davis, A., Bairoch, A., Barrett, T., Burnett, E.C., Dirks, W.G., Hall, E.M., Healy, L., Kniss, D.A., Korch, C., Liu, Y., et al. (2019). Cell Lines as Biological Models: Practical Steps for More Reliable Research. Chem. Res. Toxicol. 32, 1733–1736. 10.1021/acs.chemrestox.9b00215.

20. Simon, G.M., and Cravatt, B.F. (2010). Activity-based Proteomics of Enzyme Superfamilies: Serine Hydrolases as a Case Study*. Journal of Biological Chemistry 285, 11051–11055. 10.1074/jbc.R109.097600.

21. Liu, Y., Patricelli, M.P., and Cravatt, B.F. (1999). Activity-based protein profiling: the serine hydrolases. Proc Natl Acad Sci U S A 96, 14694–14699. 10.1073/pnas.96.26.14694.

22. Leung, D., Hardouin, C., Boger, D.L., and Cravatt, B.F. (2003). Discovering potent and selective reversible inhibitors of enzymes in complex proteomes. Nat Biotechnol 21, 687– 691. 10.1038/nbt826.

23. Long, J.Z., Nomura, D.K., Vann, R.E., Walentiny, D.M., Booker, L., Jin, X., Burston, J.J., Sim-Selley, L.J., Lichtman, A.H., Wiley, J.L., et al. (2009). Dual blockade of FAAH and MAGL identifies behavioral processes regulated by endocannabinoid crosstalk in vivo. Proceedings of the National Academy of Sciences 106, 20270–20275. 10.1073/pnas.0909411106.

24. Long, J.Z., Li, W., Booker, L., Burston, J.J., Kinsey, S.G., Schlosburg, J.E., Pavón, F.J., Serrano, A.M., Selley, D.E., Parsons, L.H., et al. (2009). Selective blockade of 2-arachidonoylglycerol hydrolysis produces cannabinoid behavioral effects. Nat Chem Biol 5, 37–44. 10.1038/nchembio.129.

25. Baggelaar, M.P., Janssen, F.J., van Esbroeck, A.C.M., den Dulk, H., Allarà, M., Hoogendoorn, S., McGuire, R., Florea, B.I., Meeuwenoord, N., van den Elst, H., et al. (2013). Development of an Activity-Based Probe and In Silico Design Reveal Highly Selective Inhibitors for Diacylglycerol Lipase-α in Brain. Angewandte Chemie International Edition 52, 12081–12085. 10.1002/anie.201306295.

26. Hsu, K.-L., Tsuboi, K., Chang, J.W., Whitby, L.R., Speers, A.E., Pugh, H., and Cravatt, B.F. (2013). Discovery and Optimization of Piperidyl-1,2,3-Triazole Ureas as Potent, Selective, and in Vivo-Active Inhibitors of α/β-Hydrolase Domain Containing 6 (ABHD6). J. Med. Chem. 56, 8270–8279. 10.1021/jm400899c.

27. Baggelaar, M.P., Chameau, P.J.P., Kantae, V., Hummel, J., Hsu, K.-L., Janssen, F., van der Wel, T., Soethoudt, M., Deng, H., den Dulk, H., et al. (2015). Highly Selective, Reversible Inhibitor Identified by Comparative Chemoproteomics Modulates Diacylglycerol Lipase Activity in Neurons. J. Am. Chem. Soc. 137, 8851–8857. 10.1021/jacs.5b04883.

28. Baggelaar, M.P., and Van der Stelt, M. (2017). Competitive ABPP of Serine Hydrolases: A Case Study on DAGL-Alpha. In Activity-Based Proteomics: Methods and Protocols, H. S. Overkleeft and B. I. Florea, eds. (Springer), pp. 161–169. 10.1007/978-1-4939-6439-0_12.

29. Ogasawara, D., Ichu, T.-A., Vartabedian, V.F., Benthuysen, J., Jing, H., Reed, A., Ulanovskaya, O.A., Hulce, J.J., Roberts, A., Brown, S., et al. (2018). Selective blockade of the lyso-PS lipase ABHD12 stimulates immune responses in vivo. Nat Chem Biol 14, 1099–1108. 10.1038/s41589-018-0155-8.

30. Ogasawara, D., Ichu, T.-A., Jing, H., Hulce, J.J., Reed, A., Ulanovskaya, O.A., and Cravatt, B.F. (2019). Discovery and Optimization of Selective and in Vivo Active Inhibitors of the Lysophosphatidylserine Lipase α/β-Hydrolase Domain-Containing 12 (ABHD12). J. Med. Chem. 62, 1643–1656. 10.1021/acs.jmedchem.8b01958.

31. Vinogradova, E.V., Zhang, X., Remillard, D., Lazar, D.C., Suciu, R.M., Wang, Y., Bianco, G., Yamashita, Y., Crowley, V.M., Schafroth, M.A., et al. (2020). An Activity-Guided Map of Electrophile-Cysteine Interactions in Primary Human T Cells. Cell 182, 1009–1026.e29. 10.1016/j.cell.2020.07.001.

32. Backus, K.M., Correia, B.E., Lum, K.M., Forli, S., Horning, B.D., González-Páez, G.E., Chatterjee, S., Lanning, B.R., Teijaro, J.R., Olson, A.J., et al. (2016). Proteome-wide covalent ligand discovery in native biological systems. Nature 534, 570–574. 10.1038/nature18002.

33. Weerapana, E., Wang, C., Simon, G.M., Richter, F., Khare, S., Dillon, M.B.D., Bachovchin, D.A., Mowen, K., Baker, D., and Cravatt, B.F. (2010). Quantitative reactivity profiling predicts functional cysteines in proteomes. Nature 468, 790–795. 10.1038/nature09472.

34. Pagel, O., Kollipara, L., and Sickmann, A. (2021). Tandem Mass Tags for Comparative and Discovery Proteomics. Methods Mol Biol 2228, 117–131. 10.1007/978-1-0716-1024-4_9.

35. Tao, Y., Remillard, D., Vinogradova, E.V., Yokoyama, M., Banchenko, S., Schwefel, D., Melillo, B., Schreiber, S.L., Zhang, X., and Cravatt, B.F. (2022). Targeted Protein Degradation by Electrophilic PROTACs that Stereoselectively and Site-Specifically Engage DCAF1. J. Am. Chem. Soc. 144, 18688–18699. 10.1021/jacs.2c08964.

36. Sharma, H.A., Bielecki, M., Holm, M.A., Thompson, T.M., Yin, Y., Cravatt, J.B., Ware, T.B., Reed, A., Nassir, M., Ewing, T.E.-H., et al. (2025). Proteomic Ligandability Maps of Phosphorus(V) Stereoprobes Identify Covalent TLCD1 Inhibitors. J. Am. Chem. Soc. 147, 15554–15566. 10.1021/jacs.5c01944.

37. Liu, Z., Remsberg, J.R., Li, H., Njomen, E., DeMeester, K.E., Tao, Y., Xia, G., Hayward, R.E., Yoo, M., Nguyen, T., et al. (2024). Proteomic Ligandability Maps of Spirocycle Acrylamide Stereoprobes Identify Covalent ERCC3 Degraders. J. Am. Chem. Soc. 146, 10393–10406. 10.1021/jacs.3c13448.

38. Lazear, M.R., Remsberg, J.R., Jaeger, M.G., Rothamel, K., Her, H.-L., DeMeester, K.E., Njomen, E., Hogg, S.J., Rahman, J., Whitby, L.R., et al. (2023). Proteomic discovery of chemical probes that perturb protein complexes in human cells. Mol Cell 83, 1725–1742.e12. 10.1016/j.molcel.2023.03.026.

39. Xiong, Y., Reinhardt, C.J., Nguyen, T., Hoffman, M.A., Simon, G.M., Melillo, B., and Cravatt, B.F. (2026). A Global Ligandability Map of Tryptoline Butynamide Stereoprobes Identifies Covalent Inhibitors of the Actin Maturation Protease. Journal American Chemical Society 148, 22077–22090. 10.1021/jacs.6c03985.

40. Wu, C., Macleod, I., and Su, A.I. (2013). BioGPS and MyGene.info: organizing online, gene-centric information. Nucleic Acids Res 41, D561–565. 10.1093/nar/gks1114.

41. GTEx Consortium (2013). The Genotype-Tissue Expression (GTEx) project. Nat Genet 45, 580–585. 10.1038/ng.2653.

42. Inoue, K. (2017). Cellular Pathology of Pelizaeus-Merzbacher Disease Involving Chaperones Associated with Endoplasmic Reticulum Stress. Front Mol Biosci 4, 7. 10.3389/fmolb.2017.00007.

43. Khalaf, G., Mattern, C., Begou, M., Boespflug-Tanguy, O., Massaad, C., and Massaad-Massade, L. (2022). Mutation of Proteolipid Protein 1 Gene: From Severe Hypomyelinating Leukodystrophy to Inherited Spastic Paraplegia. Biomedicines 10, 1709. 10.3390/biomedicines10071709.

44. Inoue, K. (2025). Molecular pathologies and therapies for Pelizaeus-Merzbacher disease. Brain and Development 47, 104383. 10.1016/j.braindev.2025.104383.

45. Schneider, A., Länder, H., Schulz, G., Wolburg, H., Nave, K.-A., Schulz, J.B., and Simons, M. (2005). Palmitoylation is a sorting determinant for transport to the myelin membrane. J Cell Sci 118, 2415–2423. 10.1242/jcs.02365.

46. Dhaunchak, A.-S., and Nave, K.-A. (2007). A common mechanism of PLP/DM20 misfolding causes cysteine-mediated endoplasmic reticulum retention in oligodendrocytes and Pelizaeus-Merzbacher disease. Proc Natl Acad Sci U S A 104, 17813–17818. 10.1073/pnas.0704975104.

47. Collins, M.O., Woodley, K.T., and Choudhary, J.S. (2017). Global, site-specific analysis of neuronal protein S-acylation. Sci Rep 7, 4683. 10.1038/s41598-017-04580-1.

48. Hetman, J.M., Soderling, S.H., Glavas, N.A., and Beavo, J.A. (2000). Cloning and characterization of PDE7B, a cAMP-specific phosphodiesterase. Proc Natl Acad Sci U S A 97, 472–476. 10.1073/pnas.97.1.472.

49. Chen, Y., Li, S., Zhong, X., Kang, Z., and Chen, R. (2020). PDE-7 Inhibitor BRL-50481 Reduces Neurodegeneration and Long-Term Memory Deficits in Mice Following Sevoflurane Exposure. ACS Chem. Neurosci. 11, 1353–1358. 10.1021/acschemneuro.0c00106.

50. Morales-Garcia, J.A., Aguilar-Morante, D., Hernandez-Encinas, E., Alonso-Gil, S., Gil, C., Martinez, A., Santos, A., and Perez-Castillo, A. (2015). Silencing phosphodiesterase 7B gene by lentiviral-shRNA interference attenuates neurodegeneration and motor deficits in hemiparkinsonian mice. Neurobiology of Aging 36, 1160–1173. 10.1016/j.neurobiolaging.2014.10.008.

51. Zhao, T., and Liang, S.H. PDE7 as a Precision Target: Bridging Disease Modulation and Potential PET Imaging for Translational Medicine. ACS Med Chem Lett 16, 711–714. 10.1021/acsmedchemlett.5c00160.

52. Fernández-Araujo, A., Tobío, A., Alfonso, A., and Botana, L.M. (2014). Role of AKAP 149– PKA–PDE4A complex in cell survival and cell differentiation processes. The International Journal of Biochemistry & Cell Biology 53, 89–101. 10.1016/j.biocel.2014.04.028.

53. Moleschi, K., and Melacini, G. (2014). Signaling at Crossroads: The Dialogue between PDEs and PKA is Spoken in Multiple Languages. Biophys J 107, 1259–1260. 10.1016/j.bpj.2014.07.051.

54. Baillie, G.S., Scott, J.D., and Houslay, M.D. (2005). Compartmentalisation of phosphodiesterases and protein kinase A: opposites attract. FEBS Letters 579, 3264– 3270. 10.1016/j.febslet.2005.03.089.

55. Pham, X., Song, G., Lao, S., Goff, L., Zhu, H., Valle, D., and Avramopoulos, D. (2016). The DPYSL2 gene connects mTOR and schizophrenia. Transl Psychiatry 6, e933. 10.1038/tp.2016.204.

56. Suzuki, H., Li, S., Tokutomi, T., Takeuchi, C., Takahashi, M., Yamada, M., Okuno, H., Miya, F., Takenouchi, T., Numabe, H., et al. (2022). De novo non-synonymous DPYSL2 (CRMP2) variants in two patients with intellectual disabilities and documentation of functional relevance through zebrafish rescue and cellular transfection experiments. Hum Mol Genet 31, 4173–4182. 10.1093/hmg/ddac166.

57. Tang, Y., Ye, Z., Wei, Y., Lin, C., Wang, Y., and Qin, C. (2015). Vertebrate Paralogous CRMPs in Nervous System: Evolutionary, Structural, and Functional Interplay. J Mol Neurosci 55, 324–334. 10.1007/s12031-014-0327-2.

58. Stenmark, P., Ogg, D., Flodin, S., Flores, A., Kotenyova, T., Nyman, T., Nordlund, P., and Kursula, P. (2007). The structure of human collapsin response mediator protein 2, a regulator of axonal growth. Journal of Neurochemistry 101, 906–917. 10.1111/j.1471-4159.2006.04401.x.

59. Myllykoski, M., Baumann, A., Hensley, K., and Kursula, P. (2017). Collapsin response mediator protein 2: high-resolution crystal structure sheds light on small-molecule binding, post-translational modifications, and conformational flexibility. Amino Acids 49, 747–759. 10.1007/s00726-016-2376-z.

60. Desprez, F., Ung, D.C., Vourc’h, P., Jeanne, M., and Laumonnier, F. (2023). Contribution of the dihydropyrimidinase-like proteins family in synaptic physiology and in neurodevelopmental disorders. Front Neurosci 17, 1154446. 10.3389/fnins.2023.1154446.

61. DiFrancesco, J.C., and DiFrancesco, D. (2015). Dysfunctional HCN ion channels in neurological diseases. Front. Cell. Neurosci. 9. 10.3389/fncel.2015.00071.

62. Xu, X., Vysotskaya, Z.V., Liu, Q., and Zhou, L. (2010). Structural Basis for the cAMP-dependent Gating in the Human HCN4 Channel*. Journal of Biological Chemistry 285, 37082–37091. 10.1074/jbc.M110.152033.

63. Lee, C.-H., and MacKinnon, R. (2017). Structures of the human HCN1 hyperpolarization-activated channel. Cell 168, 111–120.e11. 10.1016/j.cell.2016.12.023.

64. Burtscher, V., Mount, J., Huang, J., Cowgill, J., Chang, Y., Bickel, K., Chen, J., Yuan, P., and Chanda, B. (2024). Structural basis for hyperpolarization-dependent opening of human HCN1 channel. Nat Commun 15, 5216. 10.1038/s41467-024-49599-x.

65. Wainger, B.J., DeGennaro, M., Santoro, B., Siegelbaum, S.A., and Tibbs, G.R. (2001). Molecular mechanism of cAMP modulation of HCN pacemaker channels. Nature 411, 805–810. 10.1038/35081088.

66. Novella Romanelli, M., Sartiani, L., Masi, A., Mannaioni, G., Manetti, D., Mugelli, A., and Cerbai, E. (2016). HCN Channels Modulators: The Need for Selectivity. Curr Top Med Chem 16, 1764–1791. 10.2174/1568026616999160315130832.

67. Fala, L. (2016). Corlanor (Ivabradine), First HCN Channel Blocker, FDA Approved for the Treatment of Patients with Heart Failure. Am Health Drug Benefits 9, 56–59.

68. Saponaro, A., Krumbach, J.H., Chaves-Sanjuan, A., Sharifzadeh, A.S., Porro, A., Castelli, R., Hamacher, K., Bolognesi, M., DiFrancesco, D., Clarke, O.B., et al. (2024). Structural determinants of ivabradine block of the open pore of HCN4. Proc Natl Acad Sci U S A 121, e2402259121. 10.1073/pnas.2402259121.

69. Harde, E., Hierl, M., Weber, M., Waiz, D., Wyler, R., Wach, J.-Y., Haab, R., Gundlfinger, A., He, W., Schnider, P., et al. (2024). Selective and brain-penetrant HCN1 inhibitors reveal links between synaptic integration, cortical function, and working memory. Cell Chemical Biology 31, 577–592.e23. 10.1016/j.chembiol.2023.11.004.

70. Mistrík, P., Mader, R., Michalakis, S., Weidinger, M., Pfeifer, A., and Biel, M. (2005). The murine HCN3 gene encodes a hyperpolarization-activated cation channel with slow kinetics and unique response to cyclic nucleotides. J Biol Chem 280, 27056–27061. 10.1074/jbc.M502696200.

71. Stieber, J., Stöckl, G., Herrmann, S., Hassfurth, B., and Hofmann, F. (2005). Functional expression of the human HCN3 channel. J Biol Chem 280, 34635–34643. 10.1074/jbc.M502508200.

72. He, C., Chen, F., Li, B., and Hu, Z. (2014). Neurophysiology of HCN channels: from cellular functions to multiple regulations. Prog Neurobiol 112, 1–23. 10.1016/j.pneurobio.2013.10.001.

73. Proenza, C., Tran, N., Angoli, D., Zahynacz, K., Balcar, P., and Accili, E.A. (2002). Different roles for the cyclic nucleotide binding domain and amino terminus in assembly and expression of hyperpolarization-activated, cyclic nucleotide-gated channels. J Biol Chem 277, 29634–29642. 10.1074/jbc.M200504200.

74. Marini, C., Porro, A., Rastetter, A., Dalle, C., Rivolta, I., Bauer, D., Oegema, R., Nava, C., Parrini, E., Mei, D., et al. (2018). HCN1 mutation spectrum: from neonatal epileptic encephalopathy to benign generalized epilepsy and beyond. Brain 141, 3160–3178. 10.1093/brain/awy263.

75. Castelli, R., Marini, C., Porro, A., Castellini, A., Fontana, G., Saponaro, A., Cavalleri, G., Rizzi, S., Fusco, C., Parida, A., et al. (2026). Comprehensive classification of HCN1 variants linked to neurodevelopmental disorders with and without epilepsy. Preprint at bioRxiv, 10.64898/2026.03.18.712601 https://doi.org/10.64898/2026.03.18.712601.

76. Merseburg, A., Kasemir, J., Buss, E.W., Leroy, F., Bock, T., Porro, A., Barnett, A., Tröder, S.E., Engeland, B., Stockebrand, M., et al. (2022). Seizures, behavioral deficits, and adverse drug responses in two new genetic mouse models of HCN1 epileptic encephalopathy. eLife 11, e70826. 10.7554/eLife.70826.

77. Pape, H.C. (1996). Queer current and pacemaker: the hyperpolarization-activated cation current in neurons. Annu Rev Physiol 58, 299–327. 10.1146/annurev.ph.58.030196.001503.

78. Kodirov, S.A. (2025). Delineation and functions of HCN channels in neurons. Progress in Biophysics and Molecular Biology 198, 21–31. 10.1016/j.pbiomolbio.2025.09.002.

79. Bender, R.A., and Baram, T.Z. (2008). HCN channels in developing neuronal networks. Prog Neurobiol 86, 129–140. 10.1016/j.pneurobio.2008.09.007.

80. Combe, C.L., and Gasparini, S. (2021). Ih from synapses to networks: HCN channel functions and modulation in neurons. Prog Biophys Mol Biol 166, 119–132. 10.1016/j.pbiomolbio.2021.06.002.

81. Gasparini, S., and DiFrancesco, D. (1997). Action of the hyperpolarization-activated current (Ih) blocker ZD 7288 in hippocampal CA1 neurons. Pflügers Arch 435, 99–106. 10.1007/s004240050488.

82. Schultz, P.G. (2024). Synthesis at the Interface of Chemistry and Biology. Acc Chem Res 57, 2631–2642. 10.1021/acs.accounts.4c00320.

83. Schreiber, S.L., Kotz, J.D., Li, M., Aubé, J., Austin, C.P., Reed, J.C., Rosen, H., White, E.L., Sklar, L.A., Lindsley, C.W., et al. (2015). Advancing Biological Understanding and Therapeutics Discovery with Small-Molecule Probes. Cell 161, 1252–1265. 10.1016/j.cell.2015.05.023.

84. Falkenburger, B.H., and Schulz, J.B. (2006). Limitations of cellular models in Parkinson’s disease research. J Neural Transm Suppl, 261–268. 10.1007/978-3-211-45295-0_40.

85. Dolmetsch, R., and Geschwind, D.H. (2011). The human brain in a dish: the promise of iPSC-derived neurons. Cell 145, 831–834. 10.1016/j.cell.2011.05.034.

86. Cerneckis, J., Cai, H., and Shi, Y. (2024). Induced pluripotent stem cells (iPSCs): molecular mechanisms of induction and applications. Sig Transduct Target Ther 9, 112. 10.1038/s41392-024-01809-0.

87. Birtele, M., Lancaster, M., and Quadrato, G. (2025). Modelling human brain development and disease with organoids. Nat Rev Mol Cell Biol 26, 389–412. 10.1038/s41580-024-00804-1.

88. Loya-Lopez, S.I., Gomez, K., Porro, A., Thiel, G., Moroni, A., Allen, H.N., Khanna, R., and Saponaro, A. TRIP8bnano peptide prevents cAMP binding to HCN2 channels alleviating pain-like behaviors in rats with neuropathic pain. The Journal of Physiology n/a. 10.1113/JP290260.

89. Dini, L., Del Lungo, M., Resta, F., Melchiorre, M., Spinelli, V., Di Cesare Mannelli, L., Ghelardini, C., Laurino, A., Sartiani, L., Coppini, R., et al. (2018). Selective Blockade of HCN1/HCN2 Channels as a Potential Pharmacological Strategy Against Pain. Front. Pharmacol. 9. 10.3389/fphar.2018.01252.

90. Chen, S., Xu, Y., Liang, Y., Cao, Y., Lv, J., Pang, J., and Zhou, P. (2019). Identification and characterization of a series of novel HCN channel inhibitors. Acta Pharmacol Sin 40, 746– 754. 10.1038/s41401-018-0162-z.

91. Patberg, M., Oniani, T., Disse, P., Peischard, S., Vinnenberg, L., Zobeiri, M., Romanelli, M.N., Epping, L., Wiendl, H., Meuth, S.G., et al. (2023). Optimized synthesis and pharmacological evaluation of HCN channel inhibitor EC18. Archiv der Pharmazie 356, 2200665. 10.1002/ardp.202200665.

92. Sharifzadeh, A.S., Castelli, R., Porro, A., Mesirca, P., Perrier, R., Gómez, A.M., Mekrane, N., Benoit, H., Meli, A.C., Palloni, L.M.G., et al. (2025). Extracellular activation of HCN4 by a subtype-specific nanobody. Nat Commun 16, 10804. 10.1038/s41467-025-65852-3.

93. Shi, P., Melillo, B., McHenry, M.W., Camara, C.M., Yang, K., Njomen, E., Godes, M., Pazyra-Murphy, M.F., Branch, M.R., Tesar, B., Segal, R.A., Rubin, L.L., Cameron, M.D., Bird, G.H., Wales, T.E., Gygi, S.P., Cravatt, B.F., Walensky, L.D. (2026). An enantioselective covalent inhibitor of BAX confers cytoprotection in vivo. Nat Chem Bio. In press.

94. Zhang, Y., Liu, Z., Hirschi, M., Brodsky, O., Johnson, E., Won, S.J., Nagata, A., Bezwada, D., Petroski, M.D., Majmudar, J.D., et al. (2025). An allosteric cyclin E-CDK2 site mapped by paralog hopping with covalent probes. Nat Chem Biol 21, 420–431. 10.1038/s41589-024-01738-7.

95. Changeux, J.-P. (2012). Allostery and the Monod-Wyman-Changeux model after 50 years. Annu Rev Biophys 41, 103–133. 10.1146/annurev-biophys-050511-102222.

96. Cecchini, M., and Changeux, J.-P. (2022). Nicotinic receptors: From protein allostery to computational neuropharmacology. Mol Aspects Med 84, 101044. 10.1016/j.mam.2021.101044.

97. VanSchouwen, B., and Melacini, G. (2017). Regulation of HCN Ion Channels by Non-canonical Cyclic Nucleotides. Handb Exp Pharmacol 238, 123–133. 10.1007/164_2016_5006.

98. Krůsek, J. (2004). Allostery and cooperativity in the interaction of drugs with ionic channel receptors. Physiol Res 53, 569–579.

99. Krone, M.W., and Crews, C.M. (2025). Next steps for targeted protein degradation. Cell Chemical Biology 32, 219–226. 10.1016/j.chembiol.2024.10.004.

100. Hillebrand, L., Liang, X.J., Serafim, R.A.M., and Gehringer, M. (2024). Emerging and Re-emerging Warheads for Targeted Covalent Inhibitors: An Update. J. Med. Chem. 67, 7668–7758. 10.1021/acs.jmedchem.3c01825.

101. Hacker, S.M., Backus, K.M., Lazear, M.R., Forli, S., Correia, B.E., and Cravatt, B.F. (2017). Global profiling of lysine reactivity and ligandability in the human proteome. Nature Chem 9, 1181–1190. 10.1038/nchem.2826.

102. Ward, C.C., Kleinman, J.I., and Nomura, D.K. (2017). NHS-Esters As Versatile Reactivity-Based Probes for Mapping Proteome-Wide Ligandable Hotspots. ACS Chem. Biol. 12, 1478–1483. 10.1021/acschembio.7b00125.

103. Bach, K., Beerkens, B.L.H., Zanon, P.R.A., and Hacker, S.M. (2020). Light-Activatable, 2,5-Disubstituted Tetrazoles for the Proteome-wide Profiling of Aspartates and Glutamates in Living Bacteria. ACS Cent. Sci. 6, 546–554. 10.1021/acscentsci.9b01268.

104. Brulet, J.W., Borne, A.L., Yuan, K., Libby, A.H., and Hsu, K.-L. (2020). Liganding Functional Tyrosine Sites on Proteins Using Sulfur–Triazole Exchange Chemistry. Journal American Chemical Society 142, 8270–8280. 10.1021/jacs.0c00648.

105. Hahm, H.S., Toroitich, E.K., Borne, A.L., Brulet, J.W., Libby, A.H., Yuan, K., Ware, T.B., McCloud, R.L., Ciancone, A.M., and Hsu, K. L. (2020). Global targeting of functional tyrosines using sulfur-triazole exchange chemistry. Nat Chem Biol 16, 150–159. 10.1038/s41589-019-0404-5.

106. Abbasov, M.E., Kavanagh, M.E., Ichu, T.-A., Lazear, M.R., Tao, Y., Crowley, V.M., am Ende, C.W., Hacker, S.M., Ho, J., Dix, M.M., et al. (2021). A proteome-wide atlas of lysine-reactive chemistry. Nat. Chem. 13, 1081–1092. 10.1038/s41557-021-00765-4.

107. Shindo, N., and Ojida, A. (2021). Recent progress in covalent warheads for in vivo targeting of endogenous proteins. Bioorganic & Medicinal Chemistry 47, 116386. 10.1016/j.bmc.2021.116386.

108. Zhang, Z., Morstein, J., Ecker, A.K., Guiley, K.Z., and Shokat, K.M. (2022). Chemoselective Covalent Modification of K-Ras(G12R) with a Small Molecule Electrophile. Journal American Chemical Society 144, 15916–15921. 10.1021/jacs.2c05377.

109. Chen, Y., Craven, G.B., Kamber, R.A., Cuesta, A., Zhersh, S., Moroz, Y.S., Bassik, M.C., and Taunton, J. (2023). Direct mapping of ligandable tyrosines and lysines in cells with chiral sulfonyl fluoride probes. Nat. Chem. 15, 1616–1625. 10.1038/s41557-023-01281-3.

110. Zheng, Q., Zhang, Z., Guiley, K.Z., and Shokat, K.M. (2024). Strain-release alkylation of Asp12 enables mutant selective targeting of K-Ras-G12D. Nat Chem Biol 20, 1114–1122. 10.1038/s41589-024-01565-w.

111. Craven, G.B., Chu, H., Sun, J.D., Carelli, J.D., Coyne, B., Chen, H., Chen, Y., Ma, X., Das, S., Kong, W., et al. (2025). Mutant-selective AKT inhibition through lysine targeting and neo-zinc chelation. Nature 637, 205–214. 10.1038/s41586-024-08176-4.

112. Jones, L.H. (2025). Advances in sulfonyl exchange chemical biology: expanding druggable target space. Chem. Sci. 16, 10119–10140. 10.1039/d5sc02647d.

113. Kim, G., Grams, R.J., and Hsu, K.-L. (2025). Advancing Covalent Ligand and Drug Discovery beyond Cysteine. Chem. Rev. 125, 6653–6684. 10.1021/acs.chemrev.5c00001.

114. Upadhyay, T., Woods, E.C., Dela Ahator, S., Julin, K., Faucher, F.F., Uddin, M.J., Hollander, M.J., Pedowitz, N.J., Abegg, D., Hammond, I., et al. (2025). Identification of covalent inhibitors of Staphylococcus aureus serine hydrolases important for virulence and biofilm formation. Nat Commun 16, 5046. 10.1038/s41467-025-60367-3.

115. Zhao, W., Tang, Y., Gao, Y., Ding, Q., Li, Q., Li, W., and Lei, X. (2025). Quantitative Reactivity Profiling of Functional Arginine Residues in Human Cancer Cell Line Proteomes. Angewandte Chemie International Edition 64, e202515603. 10.1002/anie.202515603.

116. Wang, Y., Hu, T., Zhu, L., Xie, S., Yang, X., Xu, C., Zhai, Y., Li, Y., Huang, X., Yang, B., et al. (2026). Global profiling of arginine reactivity and ligandability in the human proteome. Nat. Chem. 18, 374–385. 10.1038/s41557-025-02012-6.

117. Wang, S., Wang, L., Hadisurya, M., Nia, S.S., Tao, W.A., Dykhuizen, E.C., and Krusemark, C.J. (2026). Covalent Protein Inhibitors via Tyrosine and Tryptophan Conjugation with Cyclic Imine Mannich Electrophiles. Angewandte Chemie International Edition 65, e16630. 10.1002/anie.202516630.

118. Nuber, C.M., Milton, A.V., Nissl, B., Isaza Alvarez, M.C., Bissinger, B.R.G., Sathian, M.B., Pignot, C.D., Haberhauer, A., Wu, D., Douat, C., et al. A Highly Reactive Cysteine-Targeted Acrylophenone Chemical Probe That Enables Peptide/Protein Bioconjugation and Chemoproteomics Analysis. JACS Au 5, 5908–5916. 10.1021/jacsau.5c00692.

119. Wozniak, J.M., Li, W., Governa, P., Chen, L.-Y., Jadhav, A., Dongre, A., Forli, S., and Parker, C.G. (2024). Enhanced mapping of small-molecule binding sites in cells. Nat Chem Biol 20, 823–834. 10.1038/s41589-023-01514-z.

120. Ogasawara, D., Konrad, D.B., Tan, Z.Y., Carey, K.L., Luo, J., Won, S.J., Li, H., Carter, T.R., DeMeester, K.E., Njomen, E., et al. (2024). Chemical tools to expand the ligandable proteome: Diversity-oriented synthesis-based photoreactive stereoprobes. Cell Chemical Biology 31, 2138–2155.e32. 10.1016/j.chembiol.2024.10.005.

121. Deutsch, E.W., Bandeira, N., Perez-Riverol, Y., Sharma, V., Carver, J.J., Mendoza, L., Kundu, D.J., Bandla, C., Kamatchinathan, S., Hewapathirana, S., et al. (2026). The ProteomeXchange consortium in 2026: making proteomics data FAIR. Nucleic Acids Res 54, D459–D469. 10.1093/nar/gkaf1146.

122. Perez-Riverol, Y., Bai, J., Bandla, C., García-Seisdedos, D., Hewapathirana, S., Kamatchinathan, S., Kundu, D.J., Prakash, A., Frericks-Zipper, A., Eisenacher, M., et al. (2022). The PRIDE database resources in 2022: a hub for mass spectrometry-based proteomics evidences. Nucleic Acids Res 50, D543–D552. 10.1093/nar/gkab1038.

123. Rostovtsev, V.V., Green, L.G., Fokin, V.V., and Sharpless, K.B. (2002). A Stepwise Huisgen Cycloaddition Process: Copper(I)-Catalyzed Regioselective “Ligation” of Azides and Terminal Alkynes. Angewandte Chemie International Edition 41, 2596–2599. 10.1002/1521-3773(20020715)41:14%3C2596::AID-ANIE2596%3E3.0.CO;2-4.

124. Tornøe, C.W., Christensen, C., and Meldal, M. (2002). Peptidotriazoles on Solid Phase: [1,2,3]-Triazoles by Regiospecific Copper(I)-Catalyzed 1,3-Dipolar Cycloadditions of Terminal Alkynes to Azides. Journal of Organic Chemistry 67, 3057–3064. 10.1021/jo011148j.

125. Speers, A.E., Adam, G.C., and Cravatt, B.F. (2003). Activity-Based Protein Profiling in Vivo Using a Copper(I)-Catalyzed Azide-Alkyne [3 + 2] Cycloaddition. Journal American Chemical Society 125, 4686–4687. 10.1021/ja034490h.

126. Zhang, Y., Chen, K., Sloan, S.A., Bennett, M.L., Scholze, A.R., O’Keeffe, S., Phatnani, H.P., Guarnieri, P., Caneda, C., Ruderisch, N., et al. (2014). An RNA-Sequencing Transcriptome and Splicing Database of Glia, Neurons, and Vascular Cells of the Cerebral Cortex. J. Neurosci. 34, 11929–11947. 10.1523/JNEUROSCI.1860-14.2014.

127. Niwa, S., Nakamura, F., Tomabechi, Y., Aoki, M., Shigematsu, H., Matsumoto, T., Yamagata, A., Fukai, S., Hirokawa, N., Goshima, Y., et al. (2017). Structural basis for CRMP2-induced axonal microtubule formation. Sci Rep 7, 10681. 10.1038/s41598-017-11031-4.

128. Wang, H., Liu, Y., Chen, Y., Robinson, H., and Ke, H. (2005). Multiple Elements Jointly Determine Inhibitor Selectivity of Cyclic Nucleotide Phosphodiesterases 4 and 7*. Journal of Biological Chemistry 280, 30949–30955. 10.1074/jbc.M504398200.

129. Ashburner, M., Ball, C.A., Blake, J.A., Botstein, D., Butler, H., Cherry, J.M., Davis, A.P., Dolinski, K., Dwight, S.S., Eppig, J.T., et al. (2000). Gene Ontology: tool for the unification of biology. Nat Genet 25, 25–29. 10.1038/75556.

130. Thomas, P.D., Ebert, D., Muruganujan, A., Mushayahama, T., Albou, L., and Mi, H. (2022). PANTHER: Making genome-scale phylogenetics accessible to all. Protein Sci 31, 8–22. 10.1002/pro.4218.

131. Huttlin, E.L., Bruckner, R.J., Navarrete-Perea, J., Cannon, J.R., Baltier, K., Gebreab, F., Gygi, M.P., Thornock, A., Zarraga, G., Tam, S., et al. (2021). Dual proteome-scale networks reveal cell-specific remodeling of the human interactome. Cell 184, 3022–3040.e28. 10.1016/j.cell.2021.04.011.

132. Tsjokajev, A., Røberg-Larsen, H., Wilson, S.R., Dyve Lingelem, A.-B., Skotland, T., Sandvig, K., and Lundanes, E. (2020). Mass spectrometry-based measurements of cyclic adenosine monophosphate in cells, simplified using reversed phase liquid chromatography with a polar characterized stationary phase. Journal of Chromatography B 1160, 122384. 10.1016/j.jchromb.2020.122384.

133. Hu, L., Santoro, B., Saponaro, A., Liu, H., Moroni, A., and Siegelbaum, S. (2013). Binding of the auxiliary subunit TRIP8b to HCN channels shifts the mode of action of cAMP. J Gen Physiol 142, 599–612. 10.1085/jgp.201311013.

