## Supplementary Chemistry Information for "Activity-based profiling of primary brain cells identifies covalent allosteric modulators of HCN channels"

### Supplementary Information: Synthesis and Characterization of Novel Compounds

#### General considerations

All NMR spectra were recorded at 298 K unless otherwise noted. $^1$H NMR spectra were recorded on Bruker Avance series spectrometers ($^1$H, 400 MHz). $^1$H NMR data are reported as follows: chemical shift (δ), multiplicity (s = singlet, d = doublet, t = triplet, q = quartet, m = multiplet; br. = broad), coupling constants, and integration. Chemical shifts are reported in parts per million (ppm) using the appropriate solvent as reference.[1] Analytical tandem liquid chromatography-mass spectrometry (LC-MS) was performed on Agilent 1200 series LC-MSD systems equipped with Agilent G6110A or G6125A mass detectors. Analytical supercritical fluid chromatography (SFC) was performed on a Shimadzu LC system (flow rate: 3 mL/min, back pressure: 100 bar, column temperature: 35 °C) equipped with a photodiode array detector. Mass measurements for high-resolution mass spectrometry (HRMS) were performed on a Waters Xevo G2-XS TOF calibrated against sodium formate clusters and using a LeuEnk lockmass. Expected monoisotopic masses were calculated using MassLynx 4.1 and the m/z values for calibrant and lockmass were MassLynx-default values.

#### Experimental procedures and analytical data

New stereoprobes used in this study were synthesized by adapting previously reported protocols.[2,3] Experimental procedures as well as analytical data for all new stereoprobes are provided.

*Synthesis of WX-02-45 and WX-02-25*

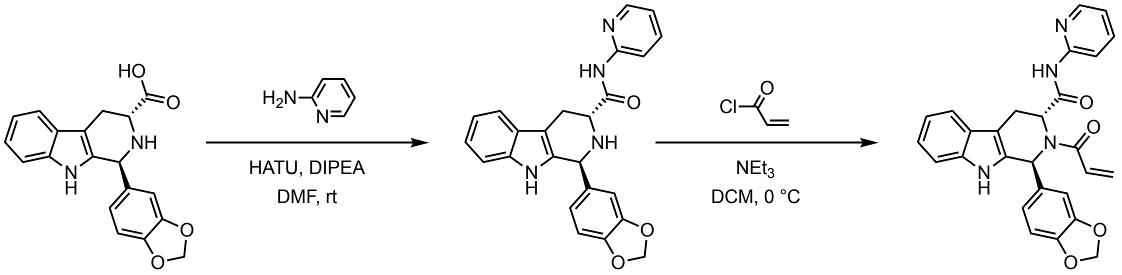


**S1** **S2** **WX-02-45**

(1*S*,3*R*)-1-(benzo[*d*][1,3]dioxol-5-yl)-*N*-(pyridin-2-yl)-2,3,4,9-tetrahydro-1*H*-pyrido[3,4-*b*]indole-3-carboxamide (S2)

To a solution of **S1** (Li salt, 70.0 mg, 204 µmol, 1 equiv) and 2-aminopyridine (29.4 mg, 312 µmol, 1.5 equiv) in DMF (2 mL) were added HATU (119 mg, 312 µmol, 1.5 equiv) and DIPEA (80.7 mg, 624 µmol, 109 µL, 3.1 equiv). The mixture was stirred at 25 °C for 1 hour. Upon reaction completion, the mixture was diluted with water (20 mL) and extracted with ethyl acetate (20 mL × 3). The combined organic layers were dried over anhydrous sodium sulfate, filtered and

concentrated under reduced pressure. The resulting residue was purified by prep-TLC (petroleum ether/EtOAc = 1:1) to obtain **S2** (50.0 mg, 59% yield) as a yellow solid.
LC-MS m/z calc. for $C_{24}H_{21}N_4O_3$ $[M+H]^+$ 413.2 found 413.1.
(1*S*,3*R*)-2-acryloyl-1-(benzo[*d*][1,3]dioxol-5-yl)-*N*-(pyridin-2-yl)-2,3,4,9-tetrahydro-1*H*-pyrido[3,4-*b*]indole-3-carboxamide (WX-02-45)
To a precooled (0 °C) solution of **S2** (50.0 mg, 121 µmol, 1 equiv) in DCM (2 mL) were added triethylamine (36.8 mg, 364 µmol, 50.6 µL, 3 equiv) and acryloyl chloride (13.2 mg, 145 µmol, 11.9 µL, 1.2 equiv). The mixture was stirred at 0 °C for 0.5 hours. Upon reaction completion, the mixture was concentrated under reduced pressure. The resulting residue was purified by prep-HPLC (mobile phase: [A: water (0.05% $NH_4OH$), B: acetonitrile]; gradient elution: 34%-64% B over 10 min) to obtain **WX-02-45** (10.0 mg, 18% yield) as an off-white solid.
$^1H$ NMR (400 MHz, $CD_3OD$) δ 8.27 – 8.20 (m, 1H), 7.97 – 7.79 (m, 1H), 7.65 (t, *J* = 7.9 Hz, 1H), 7.42 (d, *J* = 7.8 Hz, 1H), 7.26 (d, *J* = 8.1 Hz, 1H), 7.09 – 7.01 (m, 2H), 7.00 – 6.92 (m, 3H), 6.79 (dd, *J* = 16.7, 10.6 Hz, 2H), 6.36 (s, 1H), 6.19 (br. d, *J* = 16.7 Hz, 1H), 5.91 (br. s, 2H), 5.68 (dd, *J* = 10.5, 1.9 Hz, 1H), 5.52 – 5.40 (m, 1H), 3.65 – 3.50 (m, 1H), 3.46 – 3.35 (m, 1H); 2 exchangeable protons not observed.
HRMS m/z calc. for $C_{27}H_{23}N_4O_4$ $[M+H]^+$ 467.1719 found 467.1715.

(1*R*,3*S*)-2-acryloyl-1-(benzo[*d*][1,3]dioxol-5-yl)-*N*-(pyridin-2-yl)-2,3,4,9-tetrahydro-1*H*-pyrido[3,4-*b*]indole-3-carboxamide (WX-02-25)
Prepared in similar fashion from *ent*-**S1** (two-step sequence).
$^1H$ NMR (400 MHz, $CD_3OD$) δ 8.23 (ddd, *J* = 5.1, 2.0, 0.9 Hz, 1H), 7.95 – 7.78 (m, 1H), 7.63 (t, *J* = 7.8 Hz, 1H), 7.41 (d, *J* = 8.0 Hz, 1H), 7.25 (d, *J* = 8.1 Hz, 1H), 7.10 – 7.00 (m, 2H), 6.99 – 6.90 (m, 3H), 6.78 (dd, *J* = 16.7, 10.6 Hz, 2H), 6.36 (s, 1H), 6.19 (br. d, *J* = 16.6 Hz, 1H), 5.90 (br. s, 2H), 5.67 (dd, *J* = 10.6, 1.8 Hz, 1H), 5.53 – 5.38 (m, 1H), 3.67 – 3.50 (m, 1H), 3.49 – 3.34 (m, 1H); 2 exchangeable protons not observed.
HRMS m/z calc. for $C_{27}H_{23}N_4O_4$ $[M+H]^+$ 467.1719 found 467.1714.

*Synthesis of WX-02-278 and WX-02-260*

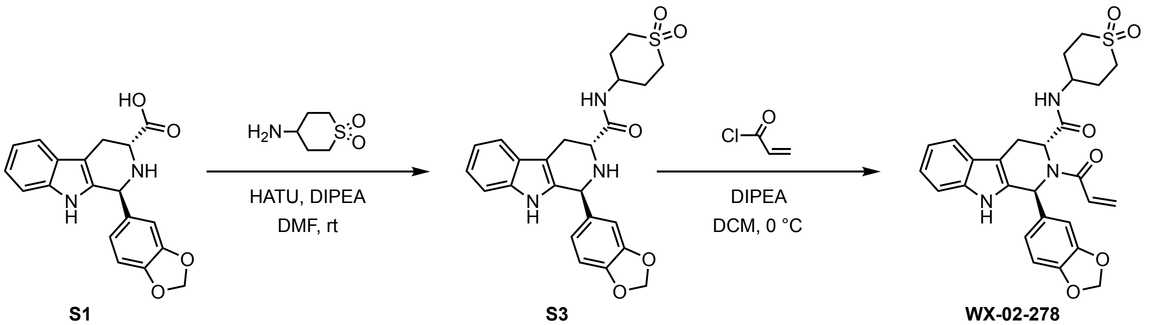

**S1** **S3** **WX-02-278**

(1*S*,3*R*)-1-(benzo[*d*][1,3]dioxol-5-yl)-*N*-(1,1-dioxidotetrahydro-2*H*-thiopyran-4-yl)-2,3,4,9-tetrahydro-1*H*-pyrido[3,4-*b*]indole-3-carboxamide (S3)
To a solution of **S1** (Li salt, 70.0 mg, 204 µmol, 1 equiv) and 4-aminotetrahydro-2*H*-thiopyran 1,1-dioxide (HCl salt, 46.4 mg, 250 µmol, 1.2 equiv) in DMF (2 mL) were added DIPEA (108 mg, 833

µmol, 4.1 equiv) and HATU (159 mg, 416 µmol, 2 equiv). The mixture was stirred at 20 °C for 2 hours. Upon reaction completion, the reaction mixture was filtered and purified by reverse-phase chromatography (mobile phase: [A: water (10 mM $NH_3{\cdot}H_2O$), B: acetonitrile]; gradient elution: 0%-100% B over 20 min) to give **S3** (60.0 mg, 96% purity [LC-UV], 60% yield) as a yellow solid.
LC-MS m/z calc. for $C_{24}H_{26}N_3O_5S$ $[M+H]^+$ 468.2 found 468.1.

(1*S*,3*R*)-2-acryloyl-1-(benzo[*d*][1,3]dioxol-5-yl)-*N*-(1,1-dioxidotetrahydro-2*H*-thiopyran-4-yl)-2,3,4,9-tetrahydro-1*H*-pyrido[3,4-*b*]indole-3-carboxamide (WX-02-278)
To a precooled (0 °C) solution of **S3** (60.0 mg, 96% purity [LC-UV], 123 µmol, 1 equiv) and DIPEA (33.2 mg, 257 µmol, 2.1 equiv) in DCM (2 mL) was added acryloyl chloride (17.4 mg, 193 µmol, 1.6 equiv). The mixture was stirred at 0 °C for 1 hour. Upon reaction completion, the mixture was concentrated under reduced pressure. The resulting residue was purified by prep-HPLC (mobile phase: [A: water (0.05% $NH_4OH$), B: acetonitrile]; gradient elution: 20%-50% B over 9 min) to obtain **WX-02-278** (29.0 mg, 45% yield) as a white solid.
$^1H$ NMR (400 MHz, $CD_3OD$) δ 7.41 (d, *J* = 7.8 Hz, 1H), 7.25 (d, *J* = 8.0 Hz, 1H), 7.10 – 6.87 (m, 4H), 6.81 – 6.73 (m, 1H), 6.70 (dd, *J* = 16.7, 10.6 Hz, 1H), 6.29 (s, 1H), 6.17 (br. d, *J* = 16.5 Hz, 1H), 5.88 (br. s, 2H), 5.69 – 5.57 (m, 1H), 5.41 – 5.25 (m, 1H), 3.89 – 3.72 (m, 1H), 3.58 – 3.36 (m, 2H), 3.18 – 2.75 (m, 4H), 2.13 – 1.69 (m, 4H); 2 exchangeable protons not observed.
HRMS m/z calc. for $C_{27}H_{28}N_3O_6S$ $[M+H]^+$ 522.1699 found 522.1688.

(1*R*,3*S*)-2-acryloyl-1-(benzo[*d*][1,3]dioxol-5-yl)-*N*-(1,1-dioxidotetrahydro-2*H*-thiopyran-4-yl)-2,3,4,9-tetrahydro-1*H*-pyrido[3,4-*b*]indole-3-carboxamide (WX-02-260)
Prepared in similar fashion from *ent*-**S1** (two-step sequence).
$^1H$ NMR (400 MHz, $CD_3OD$) δ 7.41 (d, *J* = 7.8 Hz, 1H), 7.25 (d, *J* = 8.0 Hz, 1H), 7.11 – 6.88 (m, 4H), 6.82 – 6.73 (m, 1H), 6.70 (dd, *J* = 16.7, 10.6 Hz, 1H), 6.29 (s, 1H), 6.17 (br. d, *J* = 16.7 Hz, 1H), 5.88 (br. s, 2H), 5.71 – 5.55 (m, 1H), 5.40 – 5.26 (m, 1H), 4.58 (s, 1H), 3.89 – 3.73 (m, 1H), 3.55 – 3.35 (m, 2H), 3.20 – 2.76 (m, 4H), 2.15 – 1.69 (m, 4H); 1 exchangeable proton not observed.
HRMS m/z calc. for $C_{27}H_{28}N_3O_6S$ $[M+H]^+$ 522.1699 found 522.1705.

*Synthesis of WX-02-280 and WX-02-262*

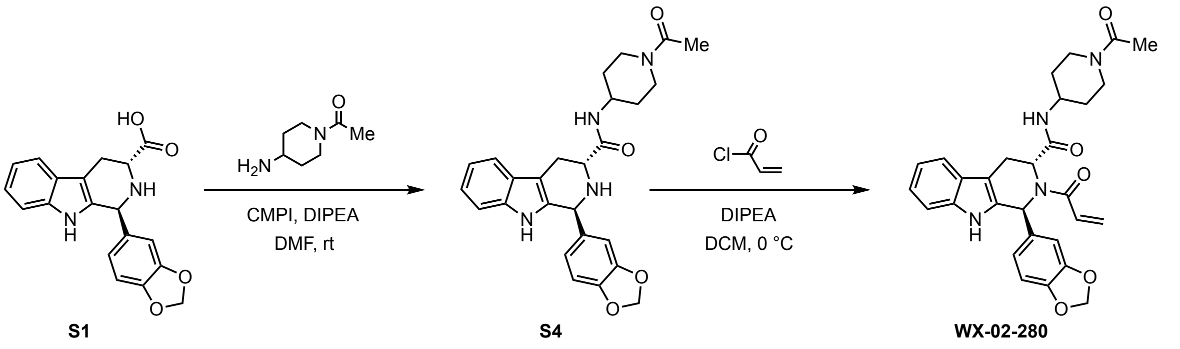

(1*S*,3*R*)-*N*-(1-acetylpiperidin-4-yl)-1-(benzo[*d*][1,3]dioxol-5-yl)-2,3,4,9-tetrahydro-1*H*-pyrido[3,4-*b*]indole-3-carboxamide (S4)

To a solution of **S1** (Li salt, 60.0 mg, 175 μmol, 1 equiv) and 1-acetyl-4-aminopiperidine (30.4 mg, 214 μmol, 1.2 equiv) in DMF (2 mL) were added DIPEA (92.2 mg, 714 μmol, 4.1 equiv) and CMPI (91.2 mg, 357 μmol, 2 equiv). The mixture was stirred at 25 °C for 2 hours. Upon reaction completion, the reaction mixture was filtered and purified by reverse-phase chromatography (mobile phase: [A: water (10 mM $NH_3$•$H_2O$), B: acetonitrile]; gradient elution: 0%-100% B over 20 min) to give **S4** (60.0 mg, 96% purity [LC-UV], 71% yield) as a yellow solid.

LC-MS m/z calc. for $C_{26}H_{29}N_4O_4$ $[M+H]^+$ 461.2 found 461.1.

(1*S*,3*R*)-*N*-(1-acetylpiperidin-4-yl)-2-acryloyl-1-(benzo[*d*][1,3]dioxol-5-yl)-2,3,4,9-tetrahydro-1*H*-pyrido[3,4-*b*]indole-3-carboxamide (WX-02-280)

To a precooled (0 °C) solution of **S4** (60.0 mg, 96% purity [LC-UV], 125 μmol, 1 equiv) and DIPEA (33.7 mg, 260 μmol, 2.1 equiv) in DCM (2 mL) was added acryloyl chloride (17.7 mg, 195 μmol, 1.6 equiv). The mixture was stirred at 0 °C for 1 hour. Upon reaction completion, the mixture was concentrated under reduced pressure. The resulting residue was purified by prep-HPLC (mobile phase: [A: water (0.05% $NH_4OH$), B: acetonitrile]; gradient elution: 19%-49% B over 9 min) to obtain **WX-02-280** (27.0 mg, 42% yield) as a white solid.

$^1H$ NMR (400 MHz, $CD_3OD$) δ 7.40 (d, *J* = 7.8 Hz, 1H), 7.25 (d, *J* = 8.0 Hz, 1H), 7.04 (ddd, *J* = 8.1, 7.0, 1.3 Hz, 1H), 7.00 – 6.90 (m, 3H), 6.76 (br. s, 1H), 6.70 (dd, *J* = 16.7, 10.6 Hz, 1H), 6.33 – 6.24 (m, 1H), 6.16 (br. d, *J* = 16.6 Hz, 1H), 5.88 (br. s, 2H), 5.74 – 5.54 (m, 1H), 5.49 – 5.21 (m, 1H), 4.33 – 4.10 (m, 1H), 3.85 – 3.56 (m, 2H), 3.54 – 3.38 (m, 2H), 3.16 – 2.88 (m, 1H), 2.79 – 2.54 (m, 1H), 2.12 – 1.92 (m, 3H), 1.84 – 1.64 (m, 1H), 1.63 – 1.40 (m, 1H), 1.40 – 1.10 (m, 2H); 2 exchangeable protons not observed; mixture of rotamers.

HRMS m/z calc. for $C_{29}H_{31}N_4O_5$ $[M+H]^+$ 515.2294 found 515.2300.

(1*R*,3*S*)-*N*-(1-acetylpiperidin-4-yl)-2-acryloyl-1-(benzo[*d*][1,3]dioxol-5-yl)-2,3,4,9-tetrahydro-1*H*-pyrido[3,4-*b*]indole-3-carboxamide (WX-02-262)

Prepared in similar fashion from *ent*-**S1** (two-step sequence).

$^1H$ NMR (400 MHz, $CD_3OD$) δ 7.40 (d, *J* = 7.8 Hz, 1H), 7.25 (d, *J* = 8.0 Hz, 1H), 7.04 (ddd, *J* = 8.2, 7.0, 1.3 Hz, 1H), 7.01 – 6.87 (m, 3H), 6.76 (br. s, 1H), 6.70 (dd, *J* = 16.7, 10.6 Hz, 1H), 6.32 – 6.23 (m, 1H), 6.17 (br. d, *J* = 16.6 Hz, 1H), 5.88 (br. s, 2H), 5.71 – 5.56 (m, 1H), 5.48 – 5.22 (m, 1H), 4.34 – 4.11 (m, 1H), 3.85 – 3.59 (m, 2H), 3.55 – 3.37 (m, 2H), 3.14 – 2.89 (m, 1H), 2.81 – 2.55 (m, 1H), 2.14 – 1.92 (m, 3H), 1.87 – 1.65 (m, 1H), 1.64 – 1.41 (m, 1H), 1.40 – 1.12 (m, 2H); 2 exchangeable protons not observed; mixture of rotamers.

HRMS m/z calc. for $C_{29}H_{31}N_4O_5$ $[M+H]^+$ 515.2294 found 515.2303.

*Synthesis of WX-02-677 and WX-02-676*

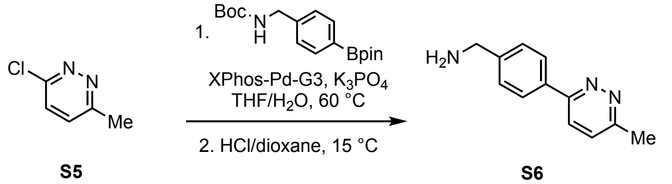


(4-(6-methylpyridazin-3-yl)phenyl)methanamine (S6)

A reactor containing a mixture of **S5** (1.00 g, 7.78 mmol, 1 equiv), 4-(*N*-Boc-aminomethyl)phenylboronic acid, pinacol ester (2.72 g, 8.17 mmol, 1.05 equiv), $K_3PO_4$ (4.95 g, 23.3 mmol, 3 equiv) and XPhos-Pd-G3 (658 mg, 778 µmol, 10 mol%) in THF (20 mL) and water (2 mL) was evacuated and backfilled with nitrogen (× 3). The mixture was then warmed to 60 °C and stirred for 2 hours under nitrogen atmosphere. Upon completion, the reaction mixture was concentrated under reduced pressure. The resulting residue was purified by silica gel column chromatography (petroleum ether/EtOAc = 1:0 to 0:1) to give a yellow solid (1.60 g), which was taken up in HCl/dioxane (4 M, 16 mL) and stirred at 15 °C for 1 hour to remove the amine protecting group (Boc). Upon reaction completion, the mixture was concentrated under reduced pressure to give **S6** (HCl salt, 1.20 g, 65% yield) as a pink solid.

$^1$H NMR (400 MHz, $CD_3OD$) δ 8.97 (d, *J* = 9.1 Hz, 1H), 8.50 (d, *J* = 9.0 Hz, 1H), 8.27 (d, *J* = 8.4 Hz, 2H), 7.75 (d, *J* = 8.3 Hz, 2H), 4.27 (s, 2H), 2.96 (s, 3H); 3 exchangeable protons not observed. LC-MS m/z calc. for $C_{12}H_{14}N_3$ $[M+H]^+$ 200.1 found 200.2.

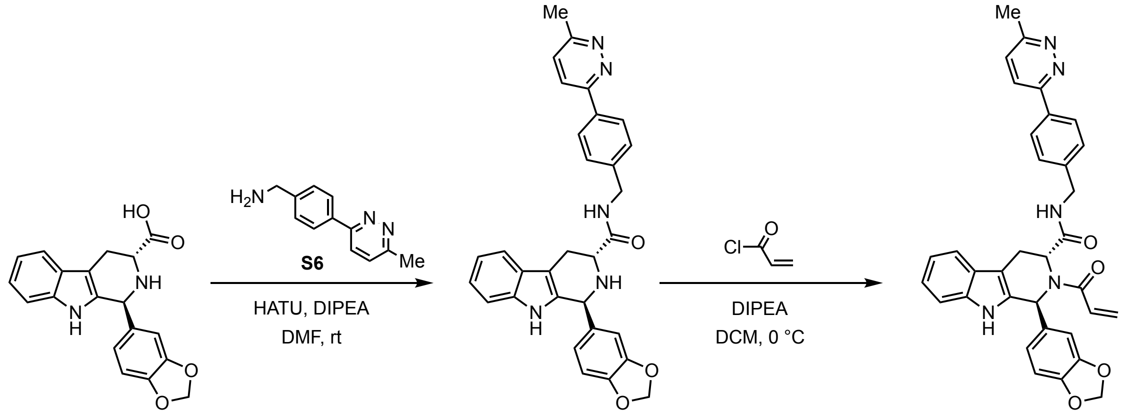


(1*S*,3*R*)-2-acryloyl-1-(benzo[*d*][1,3]dioxol-5-yl)-*N*-(4-(6-methylpyridazin-3-yl)benzyl)-2,3,4,9-tetrahydro-1*H*-pyrido[3,4-*b*]indole-3-carboxamide (WX-02-677)

To a solution of **S1** (Li salt, 100 mg, 292 µmol, 1 equiv) and **S6** (HCl salt, 80.0 mg, 339 µmol, 1.2 equiv) in DMF (10 mL) were added DIPEA (192 mg, 1.49 mmol, 259 µL, 5.1 equiv) and HATU (170 mg, 446 µmol, 1.5 equiv). The mixture was stirred at 15 °C for 1 hour. Upon reaction completion, the mixture was diluted with water (15 mL) and extracted with ethyl acetate (10 mL × 3). The combined organic layers were washed with brine (20 mL × 2), dried over anhydrous sodium sulfate, filtered and concentrated under reduced pressure to give **S7** (150 mg, crude) as a yellow oil.

To a precooled (0 °C) solution of **S7** (150 mg, crude) and DIPEA (112 mg, 869 μmol, 2.9 equiv) in DCM (8 mL) was added a solution of acryloyl chloride (31.5 mg, 348 μmol, 1.2 equiv) in DCM (2 mL). The mixture was stirred at 0 °C for 0.5 hours. Upon reaction completion, water (20 mL) was added and the mixture extracted with DCM (15 mL × 3). The combined organic layers were dried over anhydrous sodium sulfate, filtered and concentrated under reduced pressure. The resulting residue was purified by prep-TLC ($SiO_2$, MeOH/EtOAc = 10:1) and prep-HPLC (mobile phase: [A: water (0.05% $NH_4OH$), B: acetonitrile]; gradient elution: 32%-62% B over 10 min) to obtain **WX-02-677** (39.0 mg, 23% yield over two steps) as a white solid.
$^1H$ NMR (400 MHz, $CD_3OD$) δ 7.81 (d, *J* = 8.9 Hz, 1H), 7.64 (d, *J* = 8.8 Hz, 1H), 7.55 – 7.37 (m, 3H), 7.32 (d, *J* = 8.1 Hz, 1H), 7.11 (t, *J* = 7.6 Hz, 1H), 7.03 (t, *J* = 7.5 Hz, 1H), 7.00 – 6.91 (m, 2H), 6.90 – 6.62 (m, 4H), 6.36 (br. s, 1H), 6.22 (br. d, *J* = 16.4 Hz, 1H), 5.98 – 5.78 (m, 2H), 5.77 – 5.66 (m, 0.3H), 5.66 – 5.49 (m, 1.4H), 5.49 – 5.33 (m, 0.3H), 4.46 – 4.19 (m, 2H), 3.75 – 3.44 (m, 1.4H), 3.43 – 3.33 (m, 0.6H), 2.70 (s, 3H); 2 exchangeable protons not observed; 7:3 mixture of rotamers.
HRMS m/z calc. for $C_{34}H_{30}N_5O_4$ $[M+H]^+$ 572.2298 found 572.2306.

(1*R*,3*S*)-2-acryloyl-1-(benzo[*d*][1,3]dioxol-5-yl)-*N*-(4-(6-methylpyridazin-3-yl)benzyl)-2,3,4,9-tetrahydro-1*H*-pyrido[3,4-*b*]indole-3-carboxamide (WX-02-676)
Prepared in similar fashion from *ent*-**S1** (two-step sequence).
$^1H$ NMR (400 MHz, $CD_3OD$) δ 7.79 (d, *J* = 8.8 Hz, 1H), 7.63 (d, *J* = 8.8 Hz, 1H), 7.52 – 7.36 (m, 3H), 7.32 (d, *J* = 8.1 Hz, 1H), 7.11 (t, *J* = 7.5 Hz, 1H), 7.02 (d, *J* = 7.4 Hz, 1H), 7.00 – 6.91 (m, 2H), 6.90 – 6.60 (m, 4H), 6.36 (br. s, 1H), 6.22 (br. d, *J* = 16.5 Hz, 1H), 5.96 – 5.80 (m, 2H), 5.76 – 5.66 (m, 0.3H), 5.65 – 5.49 (m, 1.4H), 5.47 – 5.33 (m, 0.3H), 4.45 – 4.17 (m, 2H), 3.72 – 3.44 (m, 1.4H), 3.43 – 3.33 (m, 0.6H), 2.70 (s, 3H); 2 exchangeable protons not observed; 7:3 mixture of rotamers.
HRMS m/z calc. for $C_{34}H_{30}N_5O_4$ $[M+H]^+$ 572.2298 found 572.2313.

*Synthesis of WX-02-679 and WX-02-678*

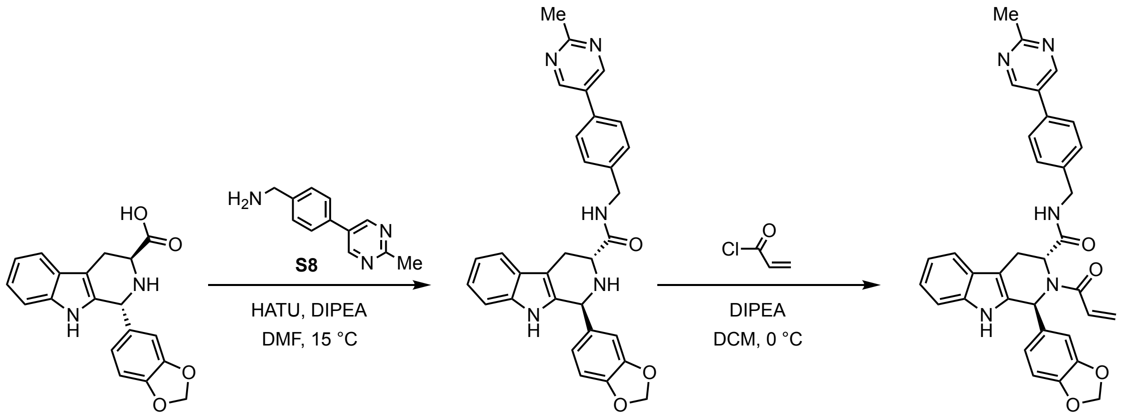

S1 S9 WX-02-679

(1*S*,3*R*)-1-(benzo[*d*][1,3]dioxol-5-yl)-*N*-(4-(2-methylpyrimidin-5-yl)benzyl)-2,3,4,9-tetrahydro-1*H*-pyrido[3,4-*b*]indole-3-carboxamide (S9)

To a precooled (0 °C) solution of **S1** (Li salt, 150 mg, 438 µmol, 1 equiv), **S8** (88.9 mg, 446 µmol, 1 equiv) and DIPEA (173 mg, 1.34 mmol, 3.1 equiv) in DMF (3 mL) was added HATU (339 mg, 892 µmol, 2 equiv). The mixture was stirred at 0 °C for 2 hours. Upon reaction completion, the mixture was filtered and concentrated under reduced pressure. The resulting residue was purified by prep-HPLC (mobile phase: [A: water (10 mM HCl), B: acetonitrile]; gradient elution: 0%-100% B over 20 min) to obtain **S9** (150 mg, 83% purity [LC-UV], 55% yield) as a white solid.

LC-MS m/z calc. for $C_{31}H_{28}N_5O_3$ $[M+H]^+$ 518.2 found 518.3.

(1*S*,3*R*)-2-acryloyl-1-(benzo[*d*][1,3]dioxol-5-yl)-*N*-(4-(2-methylpyrimidin-5-yl)benzyl)-2,3,4,9-tetrahydro-1*H*-pyrido[3,4-*b*]indole-3-carboxamide (WX-02-679)

To a precooled (0 °C) solution of **S9** (150 mg, 83% purity [LC-UV], 240 µmol, 1 equiv) and DIPEA (112 mg, 869 µmol, 3.6 equiv) in DCM (3 mL) was added acryloyl chloride (34.1 mg, 377 µmol, 1.6 equiv). The mixture was stirred at 0 °C for 1 hour. Upon reaction completion, the mixture was filtered and concentrated under reduced pressure. The resulting residue was purified by prep-TLC ($SiO_2$, petroleum ether/EtOAc = 1:1) and prep-HPLC (mobile phase: [A: water (0.05% $NH_4OH$), B: acetonitrile]; gradient elution: 30%-60% B over 9 min) to obtain **WX-02-679** (72 mg, 52% yield) as a white solid.

$^1H$ NMR (400 MHz, $CD_3OD$) δ 8.74 (s, 2H), 7.46 (d, *J* = 7.9 Hz, 1H), 7.34 (d, *J* = 8.1 Hz, 1H), 7.15 (t, *J* = 7.5 Hz, 1H), 7.05 (t, *J* = 7.4 Hz, 1H), 7.02 – 6.88 (m, 4H), 6.86 – 6.62 (m, 4H), 6.42 – 6.31 (m, 1H), 6.30 – 6.11 (m, 1H), 5.97 – 5.81 (m, 2H), 5.77 – 5.68 (m, 0.3H), 5.67 – 5.54 (m, 1.4H), 5.46 – 5.34 (m, 0.3H), 4.44 – 4.15 (m, 2H), 3.74 – 3.44 (m, 1.4H), 3.41 – 3.35 (m, 0.6H), 2.72 (s, 3H); 2 exchangeable protons not observed; 7:3 mixture of rotamers.

HRMS m/z calc. for $C_{34}H_{30}N_5O_4$ $[M+H]^+$ 572.2298 found 572.2302.

(1*R*,3*S*)-2-acryloyl-1-(benzo[*d*][1,3]dioxol-5-yl)-*N*-(4-(2-methylpyrimidin-5-yl)benzyl)-2,3,4,9-tetrahydro-1*H*-pyrido[3,4-*b*]indole-3-carboxamide (WX-02-678)

Prepared in similar fashion from *ent*-**S1** (two-step sequence).

$^1H$ NMR (400 MHz, $CD_3OD$) δ 8.74 (s, 2H), 7.46 (d, *J* = 7.8 Hz, 1H), 7.34 (d, *J* = 8.0 Hz, 1H), 7.14 (t, *J* = 7.5 Hz, 1H), 7.05 (t, *J* = 7.4 Hz, 1H), 7.03 – 6.89 (m, 4H), 6.87 – 6.61 (m, 4H), 6.38 – 6.34 (m, 1H), 6.28 – 6.12 (m, 1H), 5.97 – 5.81 (m, 2H), 5.77 – 5.68 (m, 0.3H), 5.66 – 5.53 (m, 1.4H), 5.47 – 5.33 (m, 0.3H), 4.45 – 4.16 (m, 2H), 3.73 – 3.45 (m, 1.4H), 3.42 – 3.35 (m, 0.6H), 2.72 (s, 3H); 2 exchangeable protons not observed; 7:3 mixture of rotamers.

HRMS m/z calc. for $C_{34}H_{30}N_5O_4$ $[M+H]^+$ 572.2298 found 572.2305.

*Synthesis of WX-02-922 and WX-02-921*

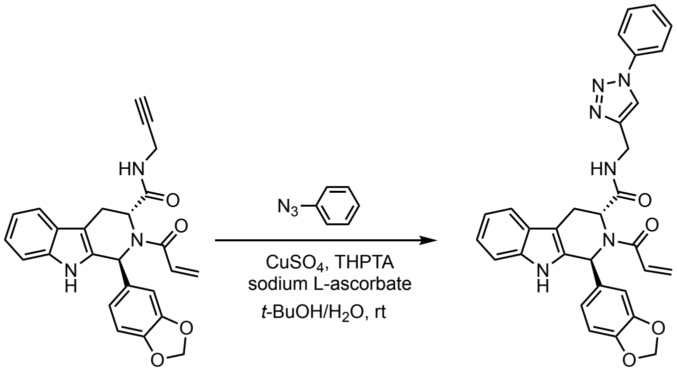


(1*S*,3*R*)-2-acryloyl-1-(benzo[*d*][1,3]dioxol-5-yl)-*N*-((1-phenyl-1*H*-1,2,3-triazol-4-yl)methyl)-2,3,4,9-tetrahydro-1*H*-pyrido[3,4-*b*]indole-3-carboxamide (WX-02-922)

To a solution of **S10** (prepared as described in ref.[3], 65.0 mg, 152 µmol, 1 equiv), phenyl azide (0.5 M in 2-MeTHF, 456 µL, 228 µmol, 1.5 equiv) and $CuSO_4$ (2.43 mg, 15.2 µmol, 10 mol%) in *t*-BuOH (1 mL) and $H_2O$ (1 mL) were added sodium L-ascorbate (45.2 mg, 228 µmol, 1.5 equiv) and THPTA (26.4 mg, 60.8 µmol, 40 mol%). The reactor was then evacuated and backfilled with nitrogen (× 3), and the mixture was stirred at 25 °C for 1 hour under nitrogen atmosphere. Upon reaction completion, the mixture was diluted with water (30 mL) and extracted with EtOAc (20 mL × 3). The combined organic layers were dried over $Na_2SO_4$, filtered and concentrated under reduced pressure. The resulting residue was purified by prep-HPLC (mobile phase: [A: water (0.05% $NH_4OH$), B: acetonitrile]; gradient elution: 28%-58% B over 15 min) to obtain **WX-02-922** (27.0 mg, 32% yield) as a yellow solid.

$^1H$ NMR (400 MHz, $CD_3OD$) δ 7.54 – 7.43 (m, 4H), 7.40 (d, *J* = 7.8 Hz, 2H), 7.30 – 7.13 (m, 2H), 7.06 – 6.84 (m, 4H), 6.82 – 6.63 (m, 2H), 6.33 (s, 1H), 6.28 – 6.10 (m, 1H), 5.88 (br. s, 2H), 5.69 – 5.56 (m, 1H), 5.50 – 5.35 (m, 1H), 4.43 (br. s, 2H), 3.66 – 3.44 (m, 1.4H), 3.37 – 3.33 (m, 0.6H); 2 exchangeable protons not observed; 7:3 mixture of rotamers.

HRMS m/z calc. for $C_{31}H_{27}N_6O_4$ $[M+H]^+$ 547.2094 found 547.2106.

(1*R*,3*S*)-2-acryloyl-1-(benzo[*d*][1,3]dioxol-5-yl)-*N*-((1-phenyl-1*H*-1,2,3-triazol-4-yl)methyl)-2,3,4,9-tetrahydro-1*H*-pyrido[3,4-*b*]indole-3-carboxamide (WX-02-921)

Prepared in similar fashion from *ent*-**S10** (prepared as described in ref.[3]).

$^1H$ NMR (400 MHz, $CD_3OD$) δ 7.55 – 7.43 (m, 4H), 7.40 (d, *J* = 7.7 Hz, 2H), 7.31 – 7.12 (m, 2H), 7.05 – 6.85 (m, 4H), 6.83 – 6.63 (m, 2H), 6.33 (s, 1H), 6.27 – 6.11 (m, 1H), 5.89 (br. s, 2H), 5.72 – 5.56 (m, 1H), 5.50 – 5.34 (m, 1H), 4.43 (br. s, 2H), 3.68 – 3.42 (m, 1.4H), 3.38 – 3.33 (m, 0.6H); 2 exchangeable protons not observed; 7:3 mixture of rotamers.

HRMS m/z calc. for $C_{31}H_{27}N_6O_4$ $[M+H]^+$ 547.2094 found 547.2099.

*Synthesis of WX-02-924 and WX-02-923*

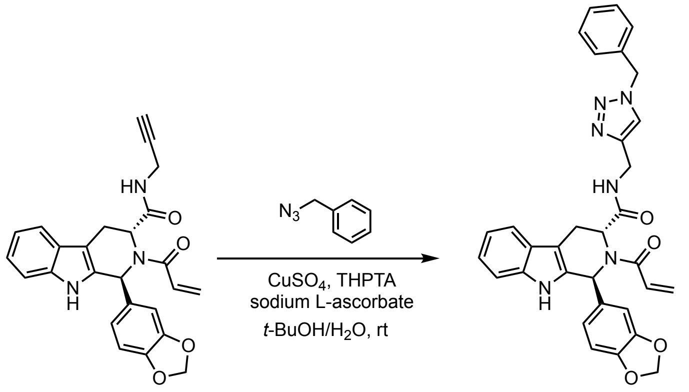


(1*S*,3*R*)-2-acryloyl-1-(benzo[*d*][1,3]dioxol-5-yl)-*N*-((1-benzyl-1*H*-1,2,3-triazol-4-yl)methyl)-2,3,4,9-tetrahydro-1*H*-pyrido[3,4-*b*]indole-3-carboxamide (WX-02-924)
Prepared in similar fashion to WX-02-922, using **S10** and benzyl azide.
$^1$H NMR (400 MHz, $CD_3OD$) δ 7.43 (d, *J* = 7.8 Hz, 1H), 7.29 (dt, *J* = 4.8, 2.4 Hz, 4H), 7.16 – 6.98 (m, 4H), 6.97 – 6.89 (m, 2H), 6.84 – 6.60 (m, 3H), 6.33 (s, 1H), 6.26 – 6.01 (m, 1H), 5.97 – 5.78 (m, 2H), 5.61 (dd, *J* = 10.4, 1.9 Hz, 1H), 5.50 – 5.27 (m, 1H), 5.15 – 4.92 (m, 2H), 4.40 – 4.19 (m, 2H), 3.65 – 3.42 (m, 1.4H), 3.36 – 3.31 (m, 0.6H + MeOH); 2 exchangeable protons not observed; 7:3 mixture of rotamers.
HRMS m/z calc. for $C_{32}H_{29}N_6O_4$ $[M+H]^+$ 561.2250 found 561.2270.

(1*R*,3*S*)-2-acryloyl-1-(benzo[*d*][1,3]dioxol-5-yl)-*N*-((1-benzyl-1*H*-1,2,3-triazol-4-yl)methyl)-2,3,4,9-tetrahydro-1*H*-pyrido[3,4-*b*]indole-3-carboxamide (WX-02-923)
Prepared in similar fashion to WX-02-921, using *ent*-**S10** and benzyl azide.
$^1$H NMR (400 MHz, $CD_3OD$) δ 7.43 (d, *J* = 7.8 Hz, 1H), 7.37 – 7.25 (m, 4H), 7.14 – 6.99 (m, 4H), 6.98 – 6.90 (m, 2H), 6.82 – 6.60 (m, 3H), 6.32 (s, 1H), 6.25 – 6.07 (m, 1H), 5.97 – 5.79 (m, 2H), 5.62 (d, *J* = 10.4 Hz, 1H), 5.51 – 5.29 (m, 1H), 5.17 – 4.93 (m, 2H), 4.39 – 4.21 (m, 2H), 3.63 – 3.45 (m, 1.4H), 3.35 – 3.31 (m, 0.6H + MeOH); 2 exchangeable protons not observed; 7:3 mixture of rotamers.
HRMS m/z calc. for $C_{32}H_{29}N_6O_4$ $[M+H]^+$ 561.2250 found 561.2255.

**Analytical data: NMR spectra**

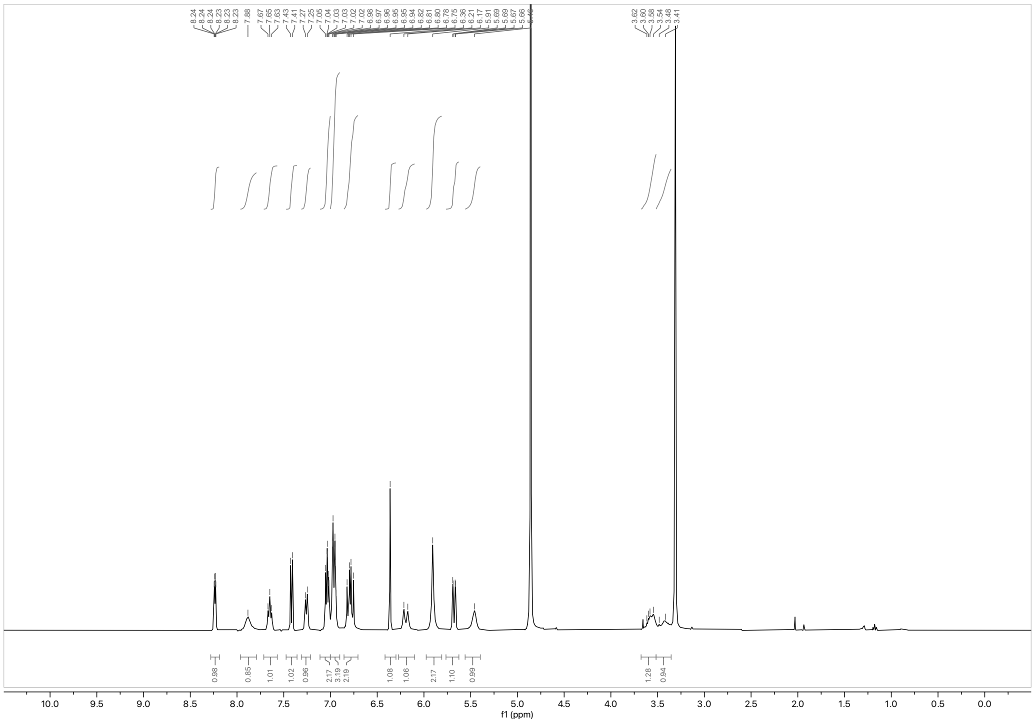

$^1H$ NMR spectrum of WX-02-45 (400 MHz, $CD_3OD$)

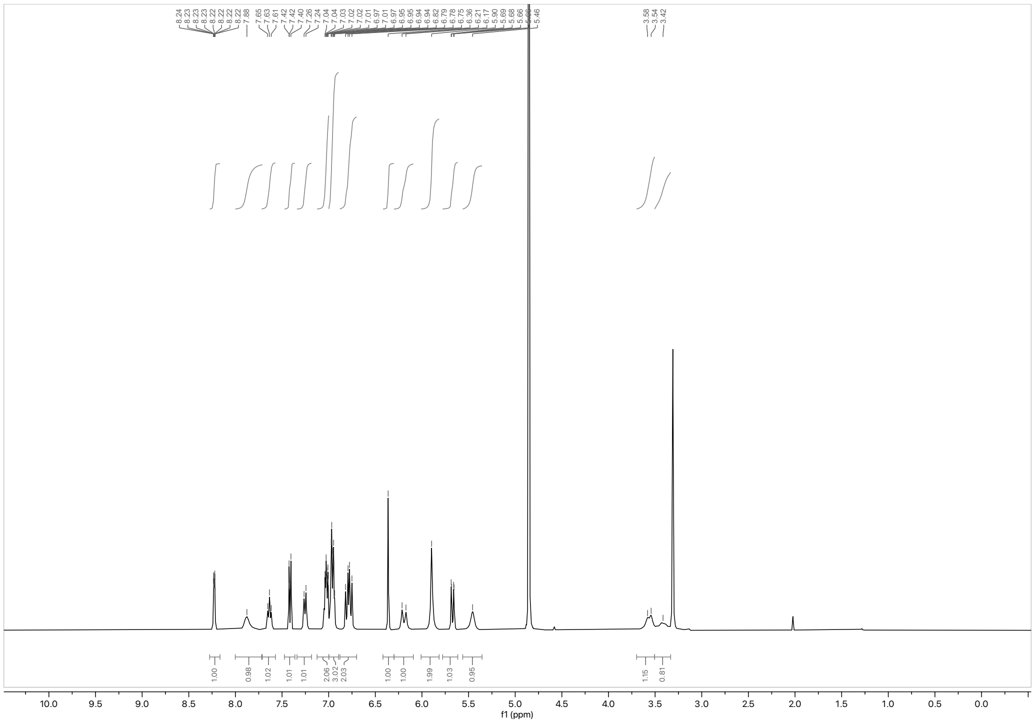

$^1H$ NMR spectrum of WX-02-25 (400 MHz, $CD_3OD$)

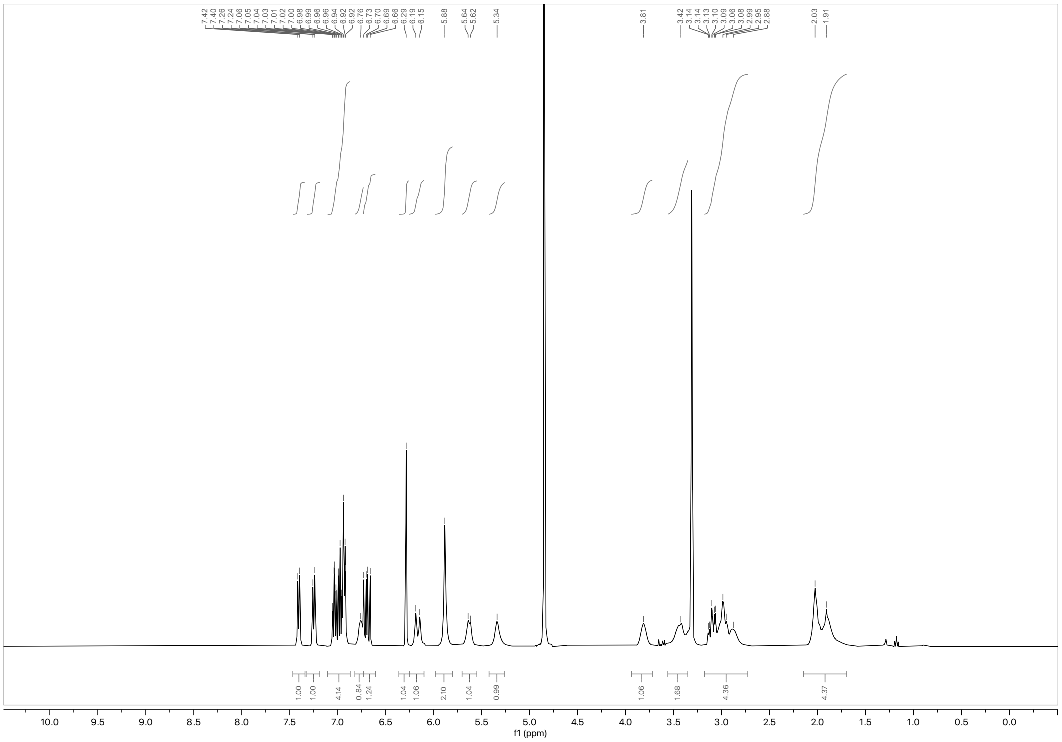

$^{1}H$ NMR spectrum of WX-02-278 (400 MHz, $CD_3OD$)

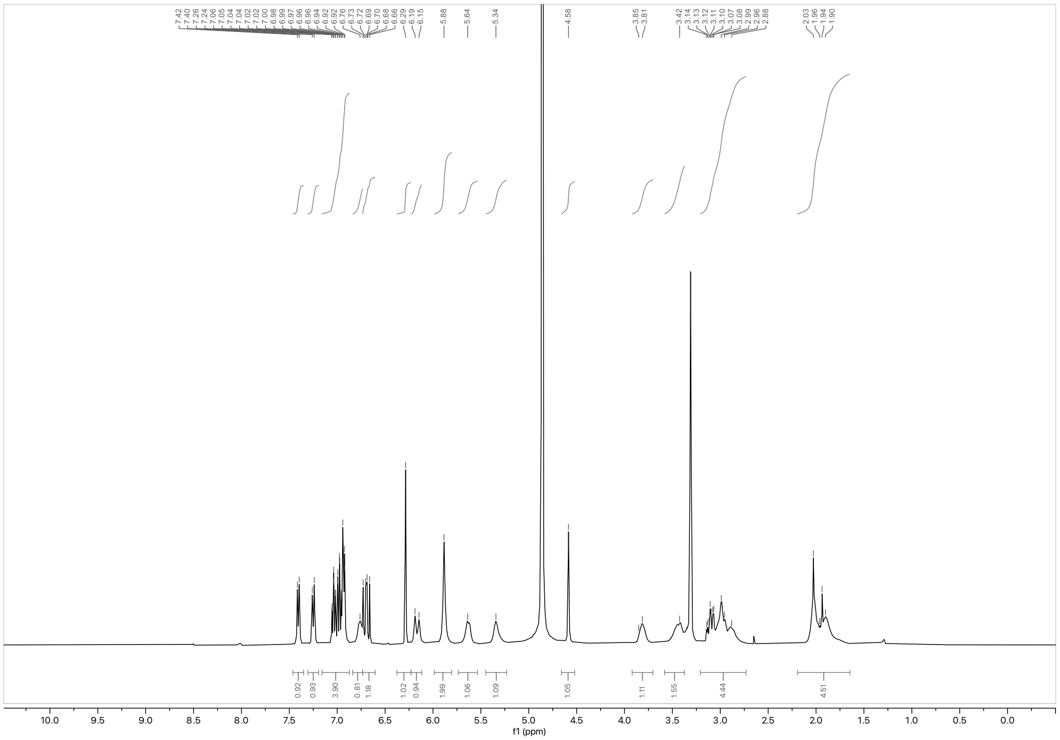

$^{1}H$ NMR spectrum of WX-02-260 (400 MHz, $CD_3OD$)

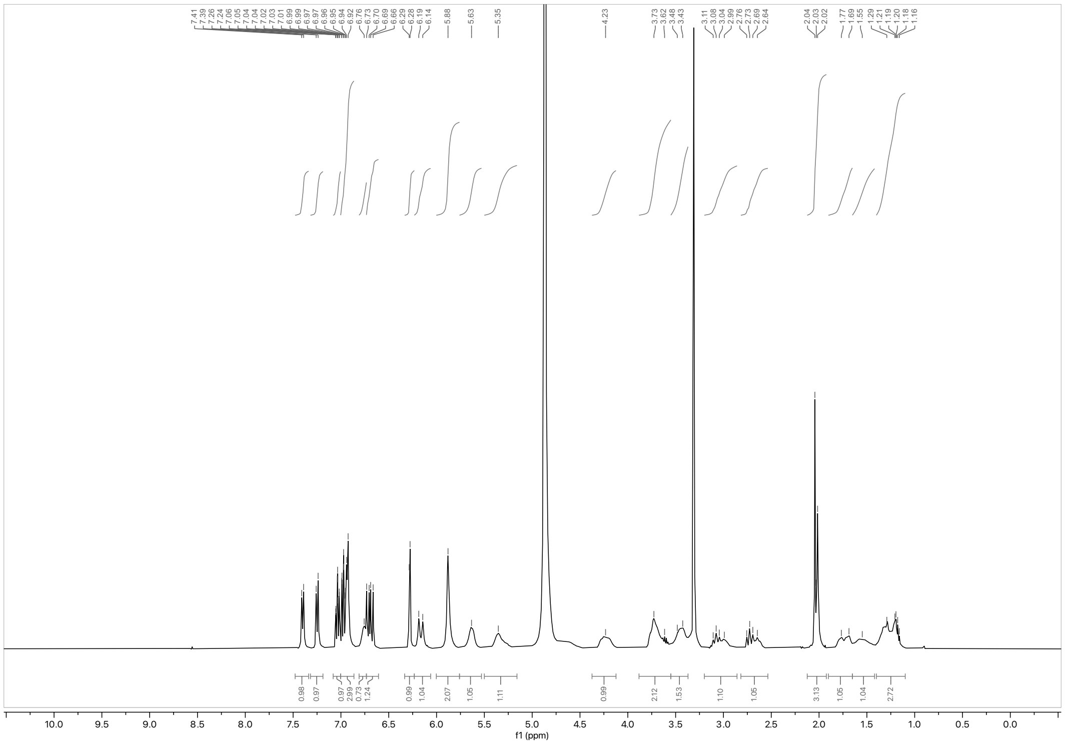

¹H NMR spectrum of WX-02-280 (400 MHz, $CD_3OD$)

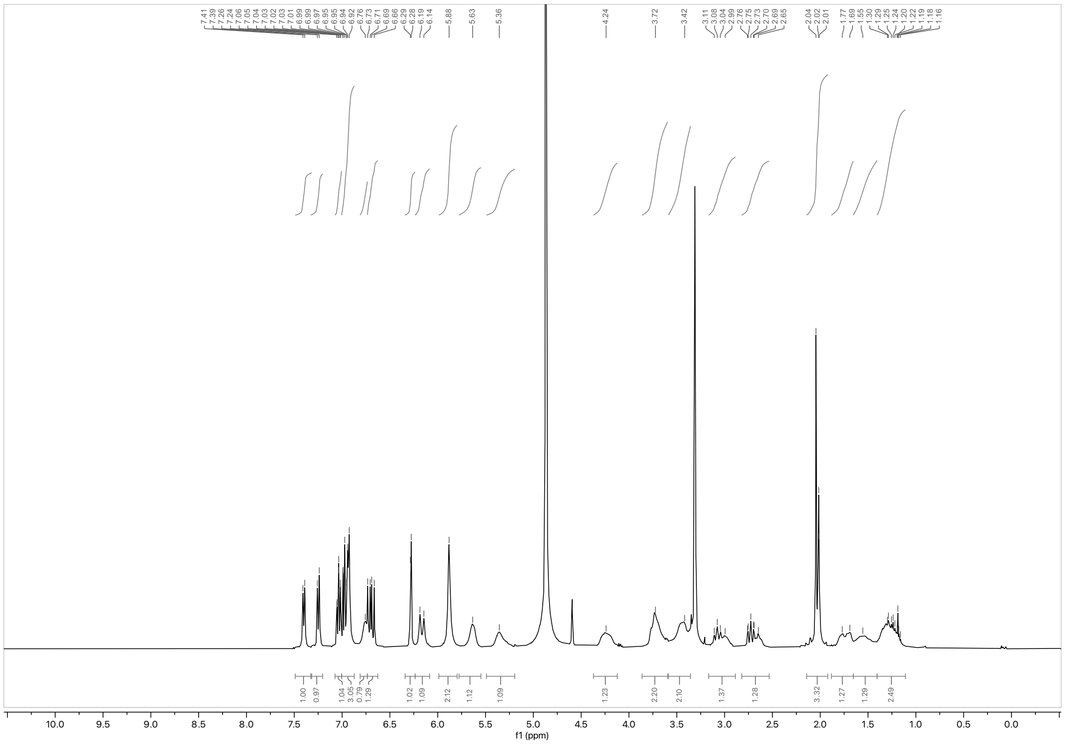

¹H NMR spectrum of WX-02-262 (400 MHz, $CD_3OD$)

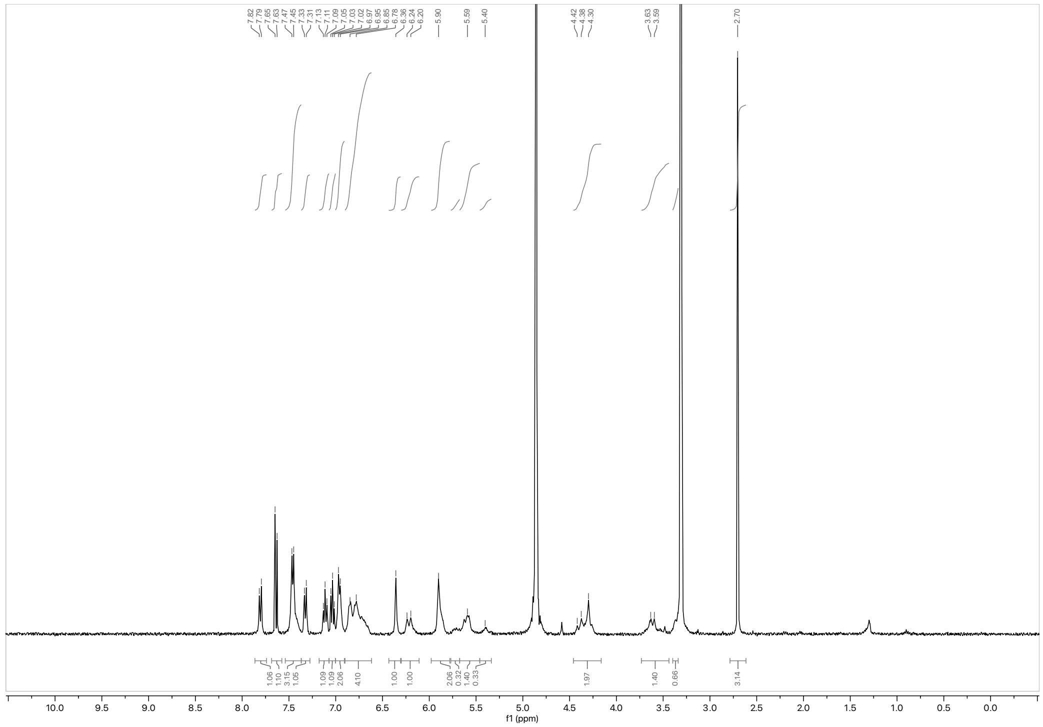

$^{1}H$ NMR spectrum of WX-02-677 (400 MHz, $CD_3OD$)

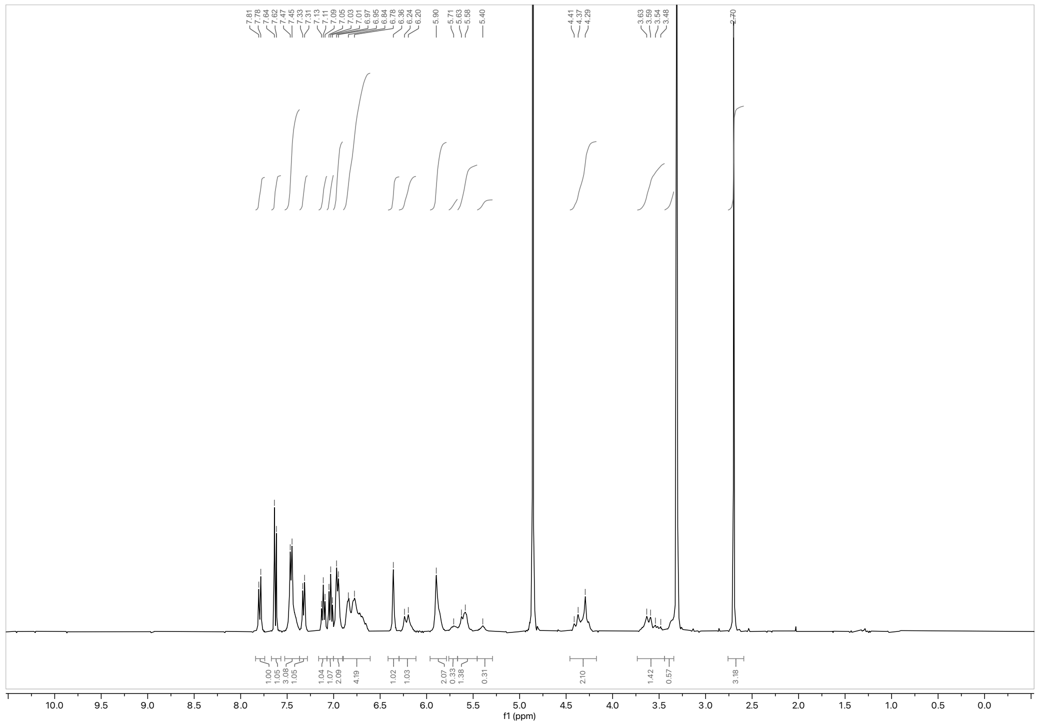

$^{1}H$ NMR spectrum of WX-02-676 (400 MHz, $CD_3OD$)

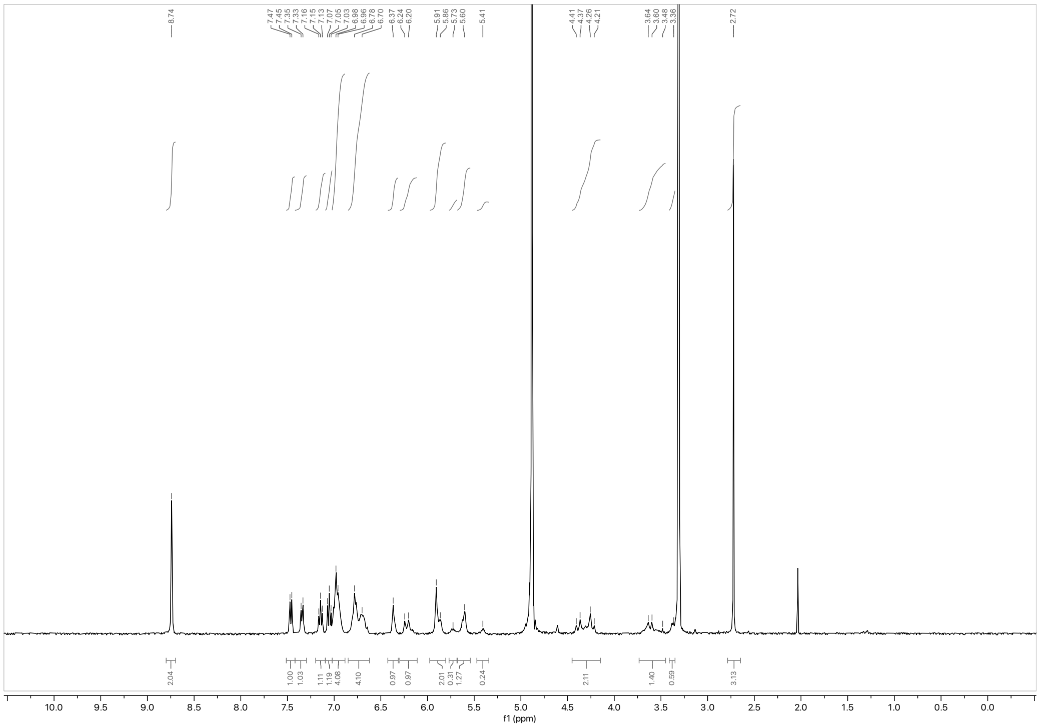

$^{1}H$ NMR spectrum of WX-02-679 (400 MHz, $CD_3OD$)

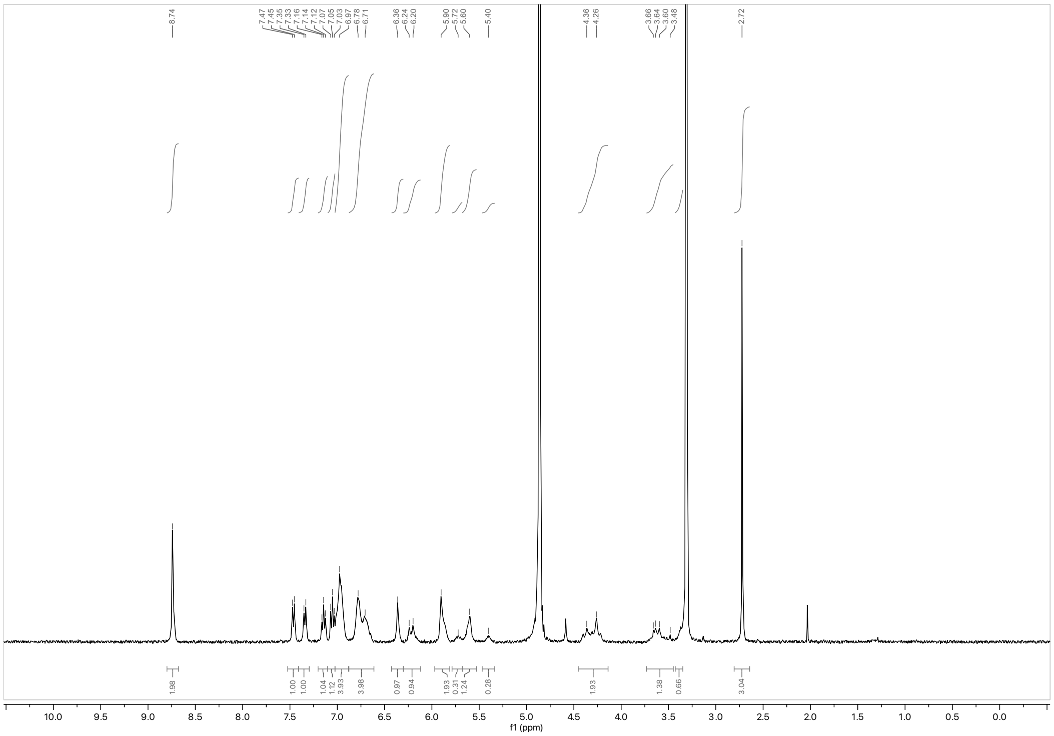

$^{1}H$ NMR spectrum of WX-02-678 (400 MHz, $CD_3OD$)

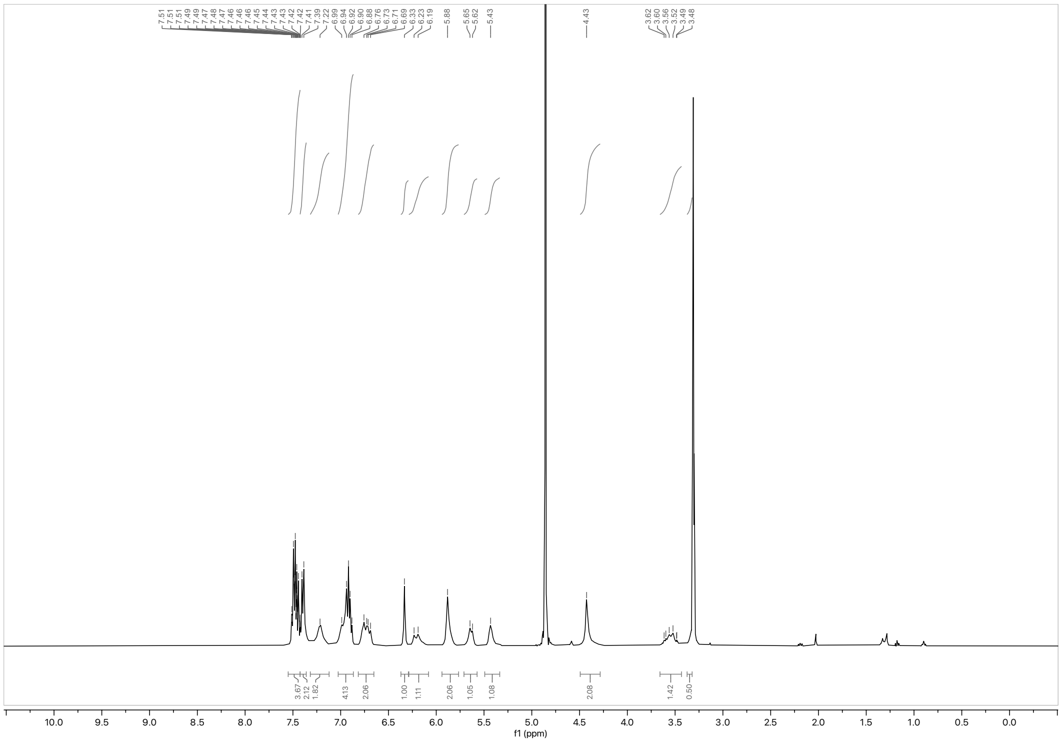

$^{1}H$ NMR spectrum of WX-02-922 (400 MHz, $CD_3OD$)

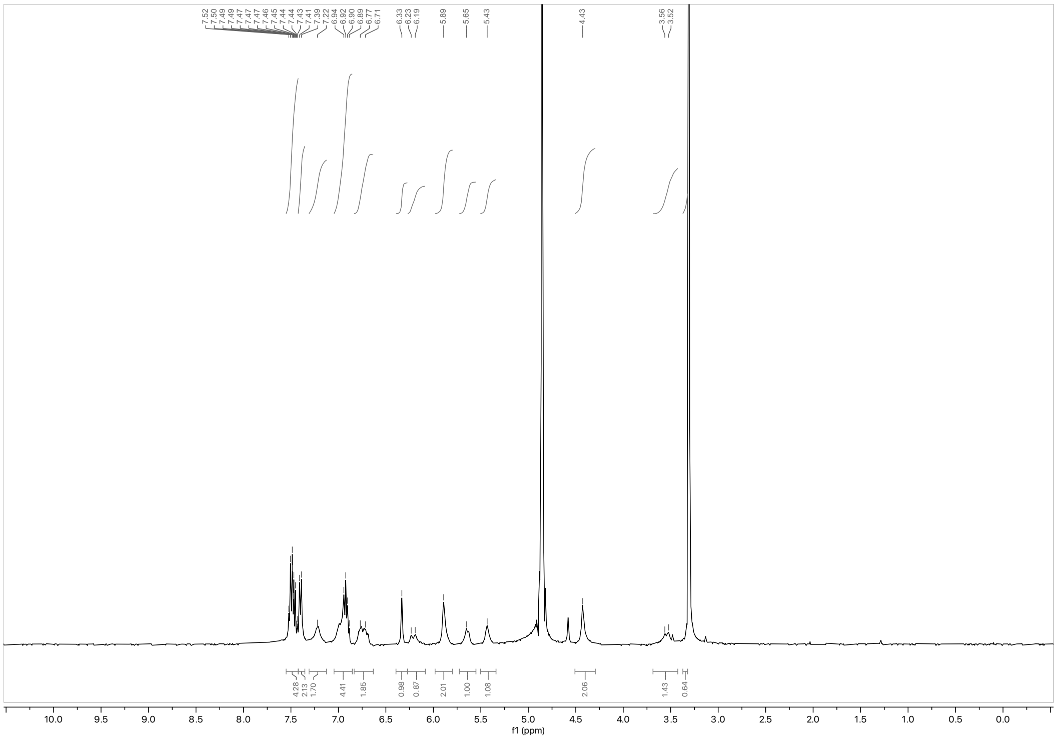

$^{1}H$ NMR spectrum of WX-02-921 (400 MHz, $CD_3OD$)

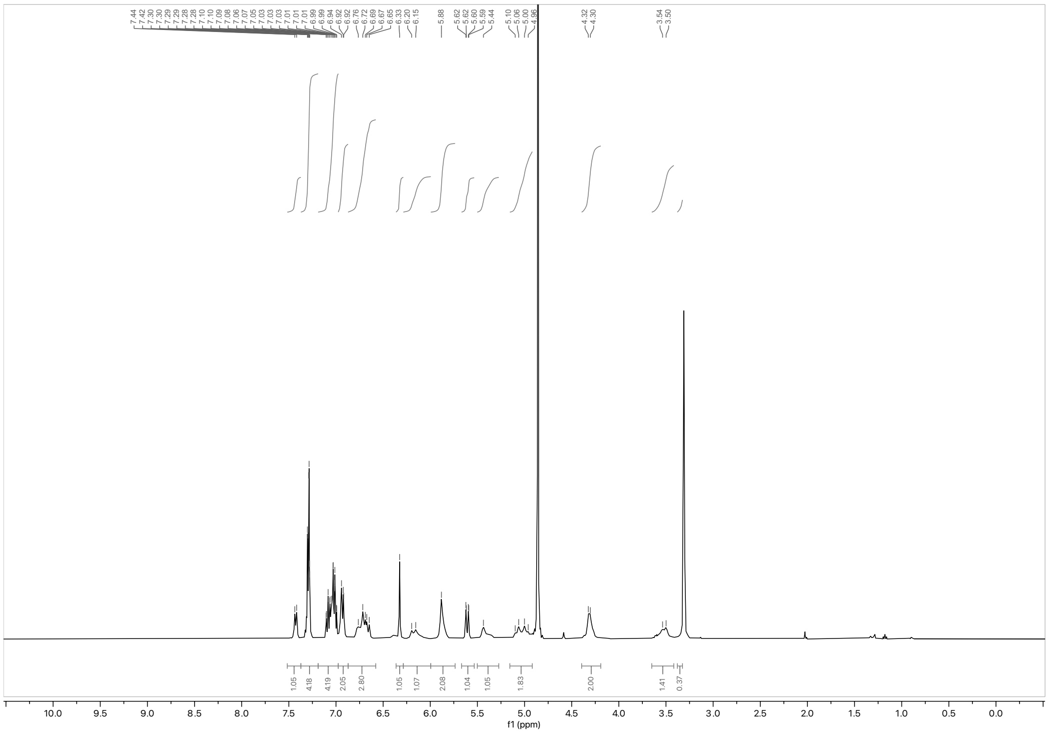

$^1$H NMR spectrum of WX-02-924 (400 MHz, $CD_3OD$)

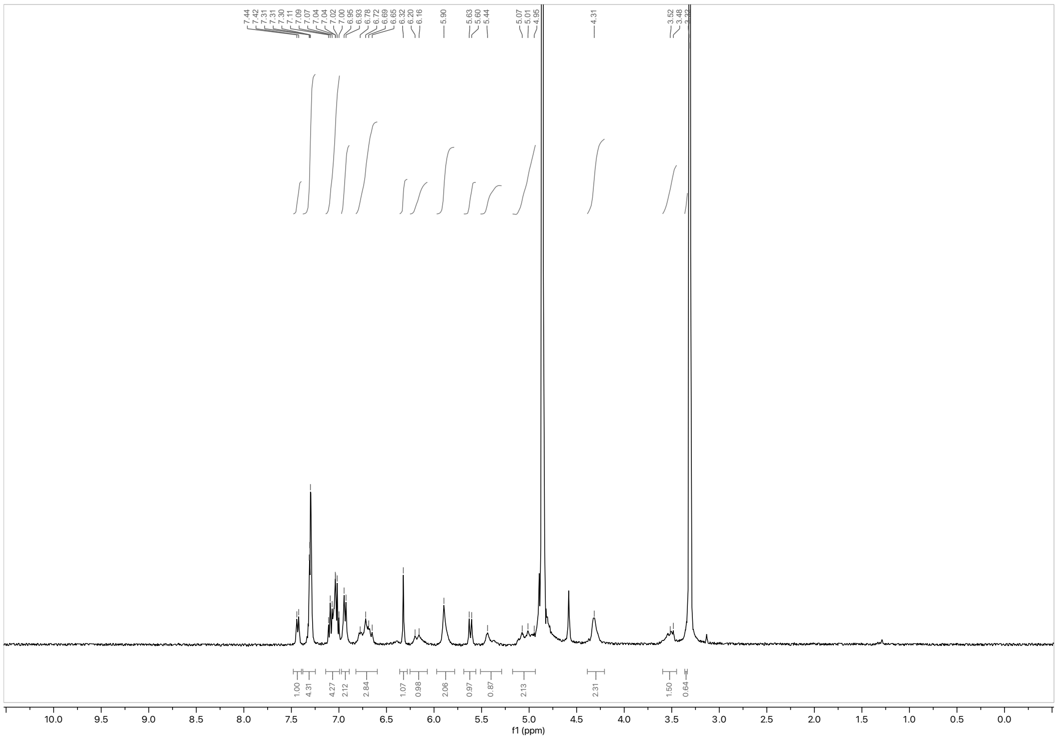

$^1$H NMR spectrum of WX-02-923 (400 MHz, $CD_3OD$)

### Analytical data: SFC

WX-02-45

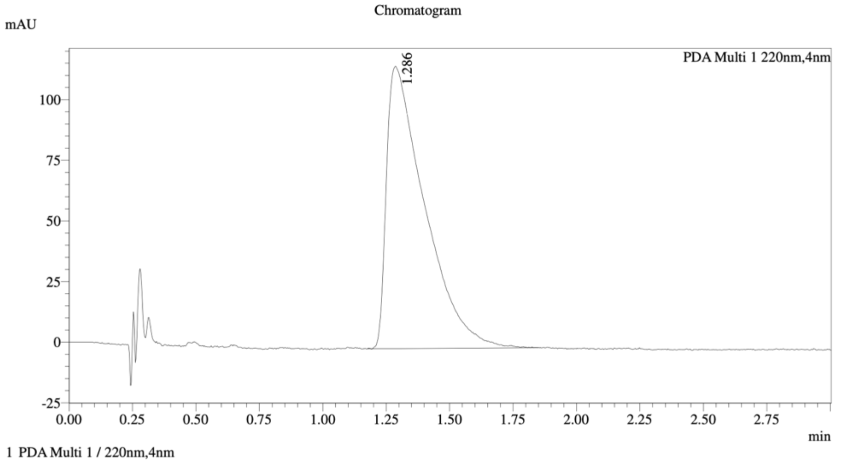


================================================================================
Integration Result
================================================================================

Peak Table

PDA Ch1 220nm

| Peak# | Ret. Time | Height | Height% | Resolution(USP) | Area | Area% |
|---|---|---|---|---|---|---|
| 1 | 1.286 | 116277 | 100.000 | -- | 1223475 | 100.000 |

WX-02-25

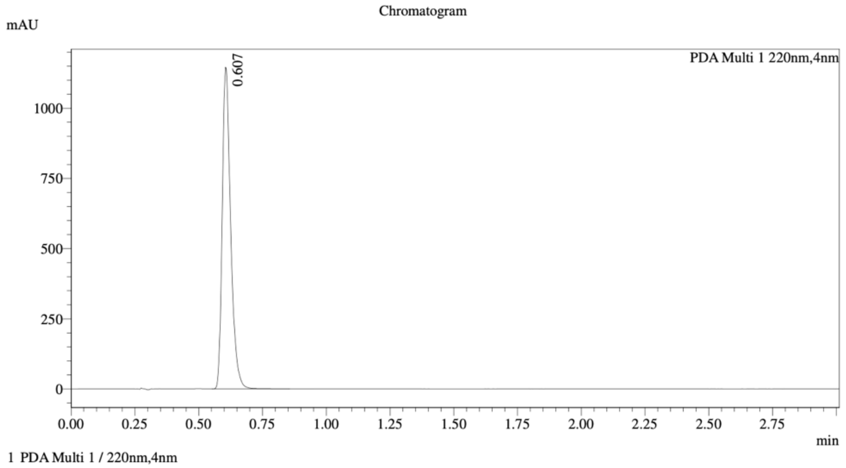


================================================================================
Integration Result
================================================================================

Peak Table

PDA Ch1 220nm

| Peak# | Ret. Time | Height | Height% | Resolution(USP) | Area | Area% |
|---|---|---|---|---|---|---|
| 1 | 0.607 | 1131883 | 100.000 | -- | 2636982 | 100.000 |

Mixture of WX-02-45 and WX-02-25

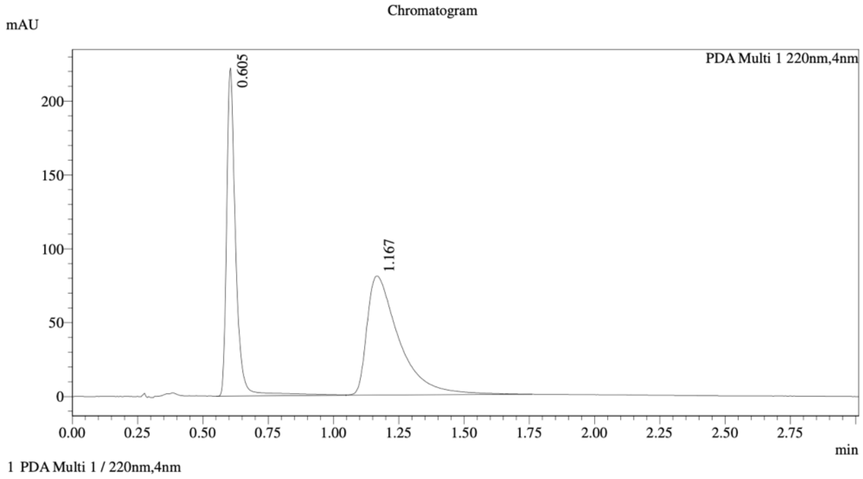


Integration Result

Peak Table

PDA Ch1 220nm

| Peak# | Ret. Time | Height | Height% | Resolution(USP) | Area | Area% |
|---|---|---|---|---|---|---|
| 1 | 0.605 | 219432 | 73.127 | -- | 539424 | 43.796 |
| 2 | 1.167 | 80637 | 26.873 | 4.134 | 692256 | 56.204 |

Method details

Column: Chiralcel OD-3 50×4.6mm I.D., 3 µm

Mobile phase: A: $CO_2$, B: MeOH (0.05% diethylamine)

Elution: 40% B

WX-02-278

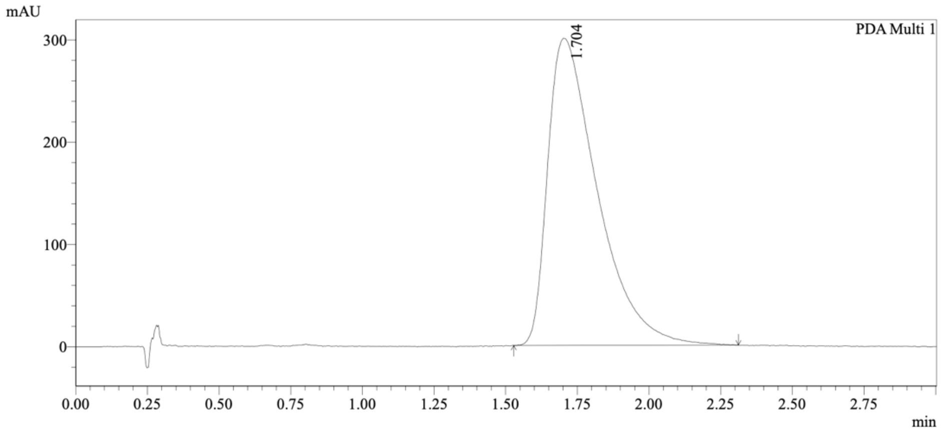


1 PDA Multi 1 / 220nm,4nm

Integration Results

PeakTable

PDA Ch1 220nm

| Peak# | Ret. Time | USP Width | Resolution | Height | Area | Area % |
|---|---|---|---|---|---|---|
| 1 | 1.704 | 0.311 | 0.000 | 300033 | 3623953 | 100.000 |
| Total | | | | 300033 | 3623953 | 100.000 |

WX-02-260

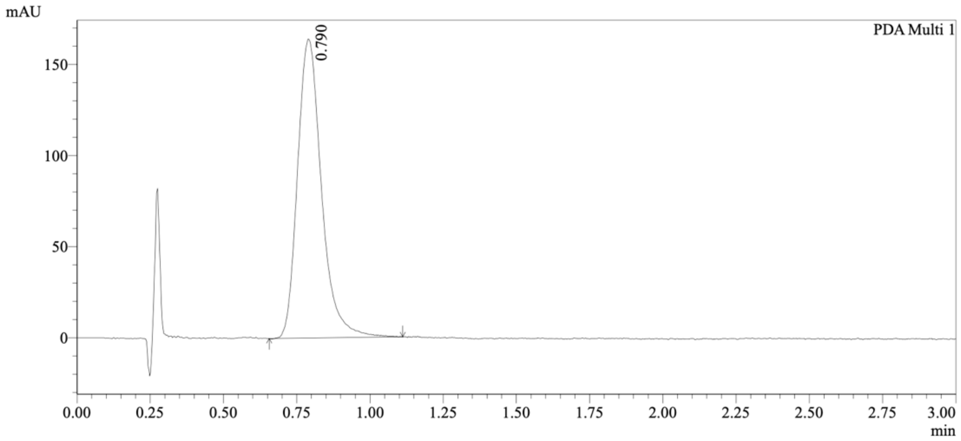


1 PDA Multi 1 / 220nm,4nm

Integration Results

PeakTable

PDA Ch1 220nm

| Peak# | Ret. Time | USP Width | Resolution | Height | Area | Area % |
|---|---|---|---|---|---|---|
| 1 | 0.790 | 0.146 | 0.000 | 163727 | 933686 | 100.000 |
| Total | | | | 163727 | 933686 | 100.000 |

Mixture of WX-02-278 and WX-02-260

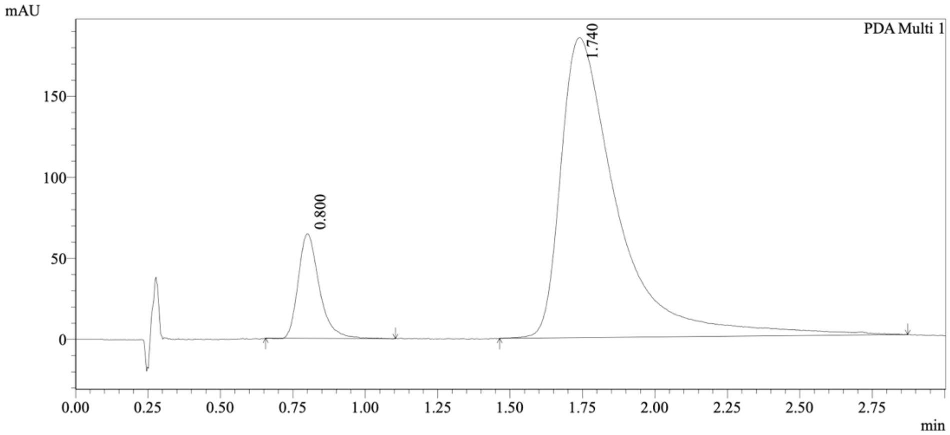


1 PDA Multi 1 / 220nm,4nm

Integration Results

PeakTable

PDA Ch1 220nm

| Peak# | Ret. Time | USP Width | Resolution | Height | Area | Area % |
|---|---|---|---|---|---|---|
| 1 | 0.800 | 0.130 | 0.000 | 64142 | 324236 | 11.381 |
| 2 | 1.740 | 0.317 | 4.205 | 185135 | 2524728 | 88.619 |
| Total | | | | 249278 | 2848964 | 100.000 |

Method details

Column: Chiralcel OD-3 50×4.6mm I.D., 3 µm
Mobile phase: A: $CO_2$, B: MeOH (0.05% diethylamine)
Elution: 40% B

WX-02-280

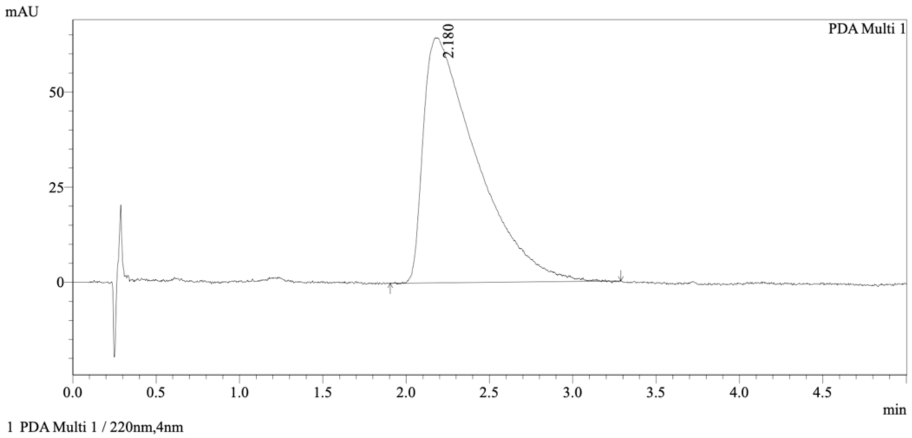


Integration Results

PeakTable

PDA Ch1 220nm

| Peak# | Ret. Time | USP Width | Resolution | Height | Area | Area % |
|---|---|---|---|---|---|---|
| 1 | 2.180 | 0.604 | 0.000 | 64310 | 1439805 | 100.000 |
| Total | | | | 64310 | 1439805 | 100.000 |

WX-02-262

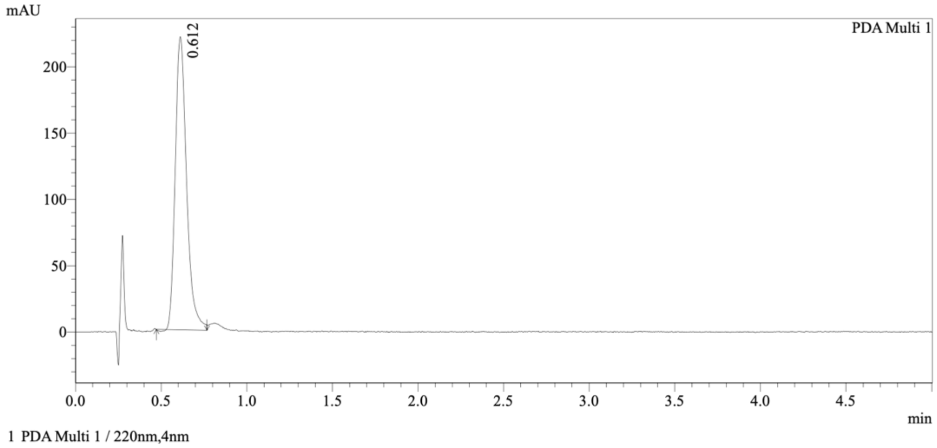


Integration Results

PeakTable

PDA Ch1 220nm

| Peak# | Ret. Time | USP Width | Resolution | Height | Area | Area % |
|---|---|---|---|---|---|---|
| 1 | 0.612 | 0.120 | 0.000 | 220343 | 1016545 | 100.000 |
| Total | | | | 220343 | 1016545 | 100.000 |

Mixture of WX-02-280 and WX-02-262

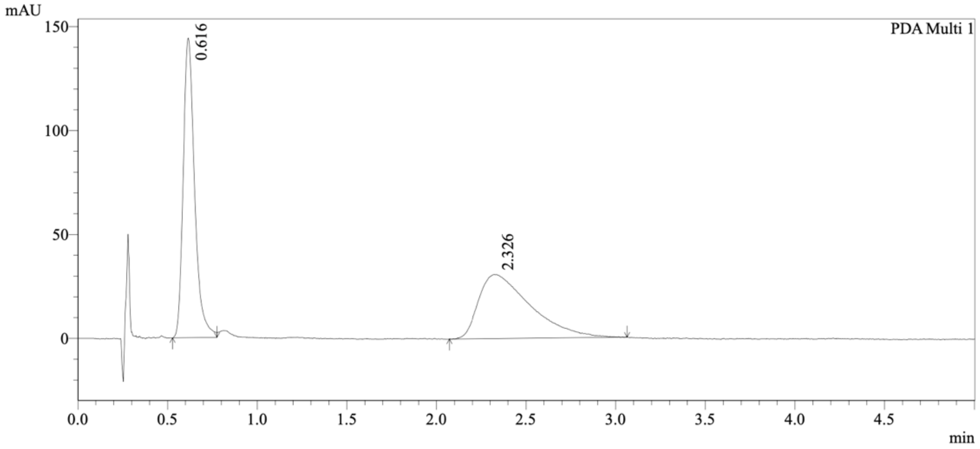


1 PDA Multi 1 / 220nm,4nm

Integration Results

PeakTable

PDA Ch1 220nm

| Peak# | Ret. Time | USP Width | Resolution | Height | Area | Area % |
|---|---|---|---|---|---|---|
| 1 | 0.616 | 0.114 | 0.000 | 143280 | 636899 | 50.805 |
| 2 | 2.326 | 0.519 | 5.409 | 30749 | 616725 | 49.195 |
| Total | | | | 174029 | 1253625 | 100.000 |

Method details

Column: Chiralcel OD-3 50×4.6mm I.D., 3 µm
Mobile phase: A: $CO_2$, B: MeOH (0.05% diethylamine)
Elution: 40% B

WX-02-677

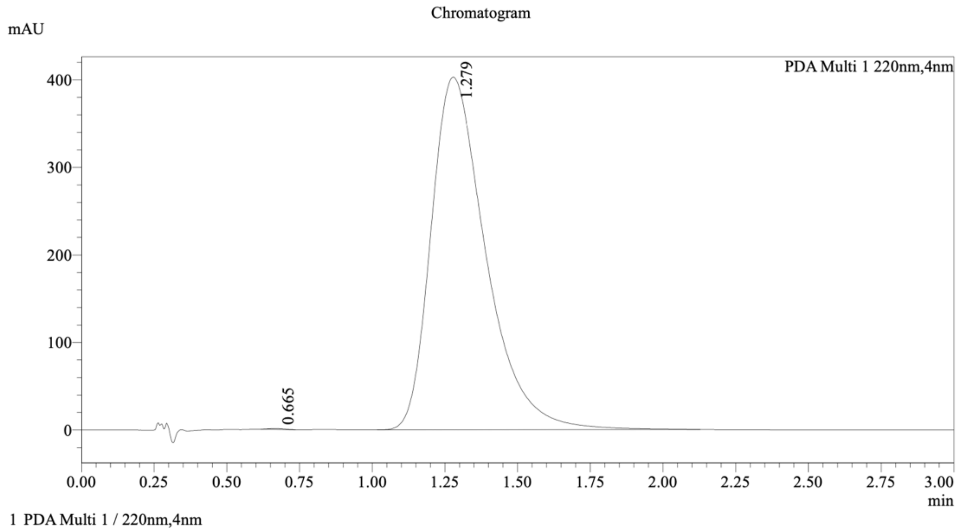


Integration Result

Peak Table

PDA Ch1 220nm

| Peak# | Ret. Time | Height | Height% | Resolution(USP) | Area | Area% |
|---|---|---|---|---|---|---|
| 1 | 0.665 | 1075 | 0.266 | -- | 3780 | 0.071 |
| 2 | 1.279 | 402591 | 99.734 | 2.832 | 5297355 | 99.929 |

WX-02-676

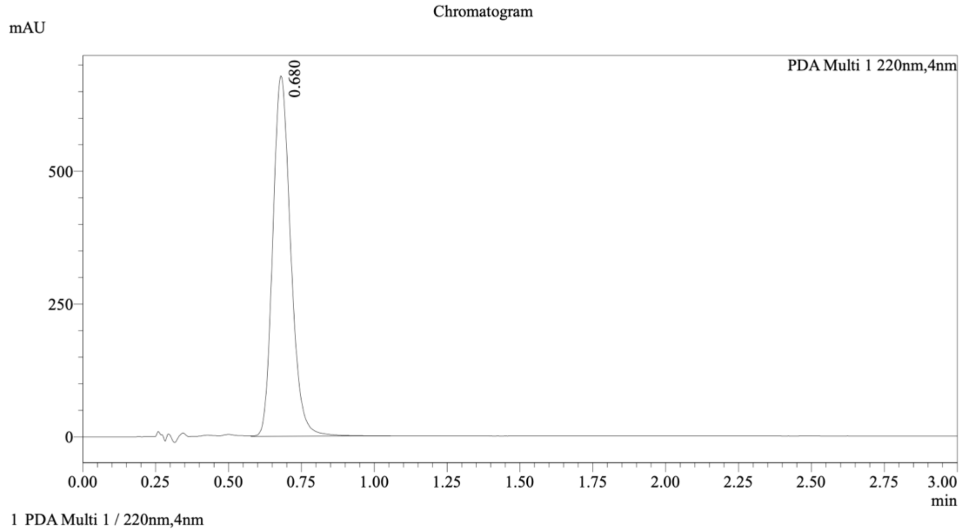


Integration Result

Peak Table

PDA Ch1 220nm

| Peak# | Ret. Time | Height | Height% | Resolution(USP) | Area | Area% |
|---|---|---|---|---|---|---|
| 1 | 0.680 | 672996 | 100.000 | -- | 2901667 | 100.000 |

Mixture of WX-02-677 and WX-02-676

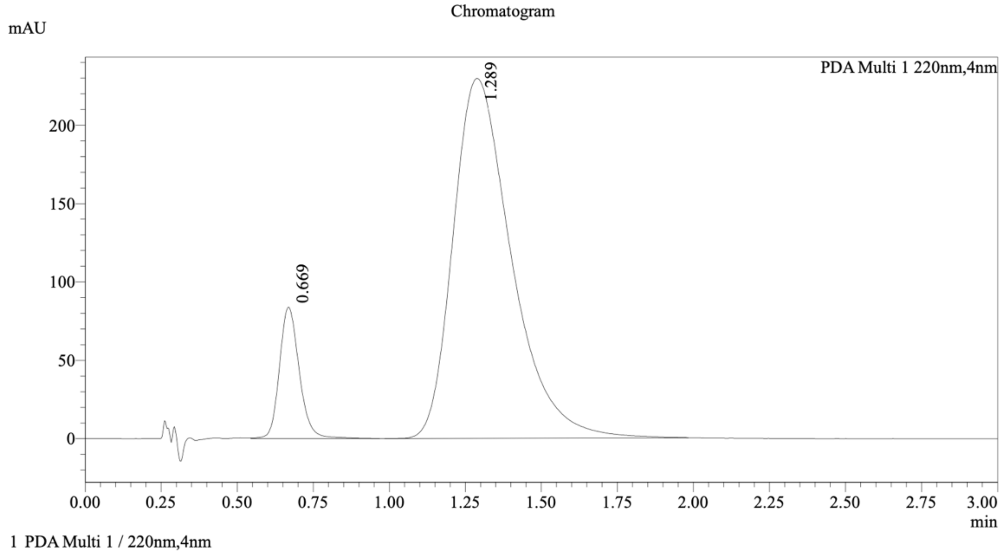

1 PDA Multi 1 / 220nm,4nm

Integration Result

Peak Table

PDA Ch1 220nm

| Peak# | Ret. Time | Height | Height% | Resolution(USP) | Area | Area% |
|---|---|---|---|---|---|---|
| 1 | 0.669 | 83657 | 26.727 | -- | 380813 | 11.027 |
| 2 | 1.289 | 229350 | 73.273 | 2.693 | 3072662 | 88.973 |

Method details

Column: Chiralpak IC-3 50×4.6mm I.D., 3 µm
Mobile phase: A: $CO_2$, B: EtOH (0.05% diethylamine)
Elution: 60% B

WX-02-679

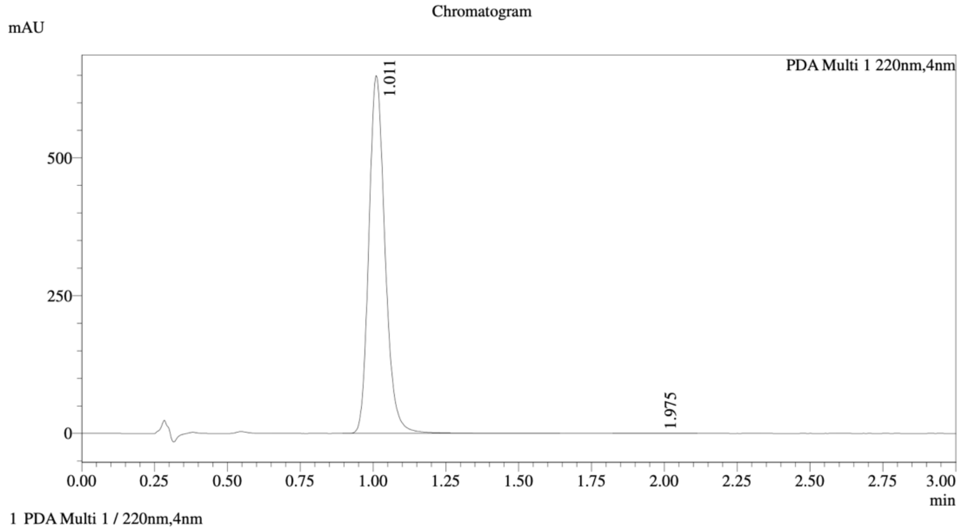


Integration Result

Peak Table

PDA Ch1 220nm

| Peak# | Ret. Time | Height | Height% | Resolution(USP) | Area | Area% |
|---|---|---|---|---|---|---|
| 1 | 1.011 | 643290 | 99.955 | -- | 2510858 | 99.905 |
| 2 | 1.975 | 288 | 0.045 | 6.270 | 2395 | 0.095 |

WX-02-678

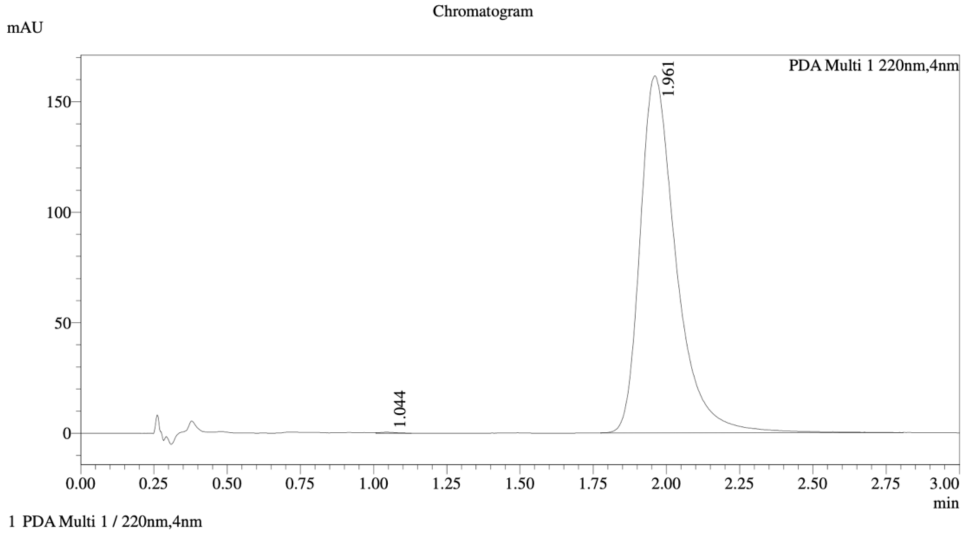


Integration Result

Peak Table

PDA Ch1 220nm

| Peak# | Ret. Time | Height | Height% | Resolution(USP) | Area | Area% |
|---|---|---|---|---|---|---|
| 1 | 1.044 | 397 | 0.246 | -- | 1450 | 0.105 |
| 2 | 1.961 | 161099 | 99.754 | 5.721 | 1382915 | 99.895 |

Mixture of WX-02-679 and WX-02-678

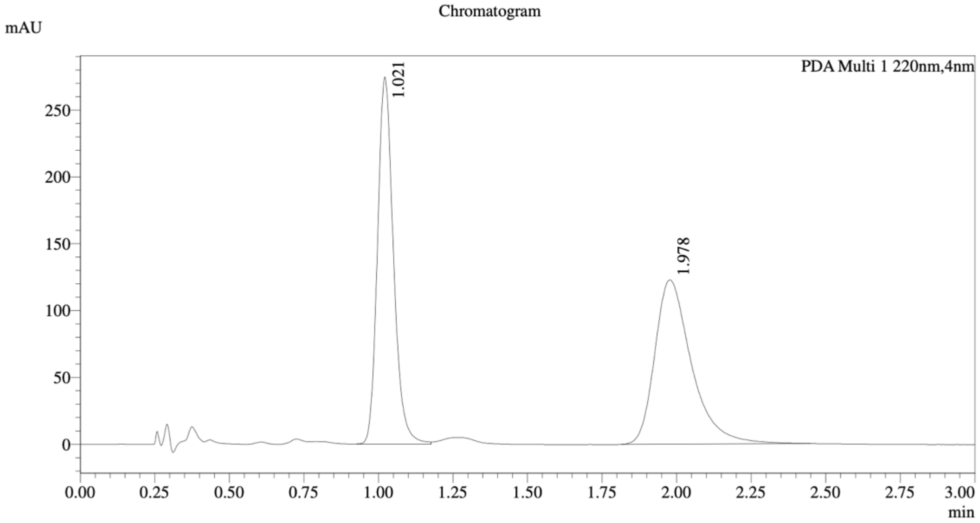


1 PDA Multi 1 / 220nm,4nm

Integration Result

Peak Table

PDA Ch1 220nm

| Peak# | Ret. Time | Height | Height% | Resolution(USP) | Area | Area% |
|---|---|---|---|---|---|---|
| 1 | 1.021 | 273451 | 69.062 | -- | 1021205 | 49.242 |
| 2 | 1.978 | 122497 | 30.938 | 6.085 | 1052628 | 50.758 |

Method details

Column: (S,S)Whelk-O1 50×4.6mm I.D., 3.5 µm

Mobile phase: A: $CO_2$, B: EtOH (0.05% diethylamine)

Elution: 60% B

WX-02-922

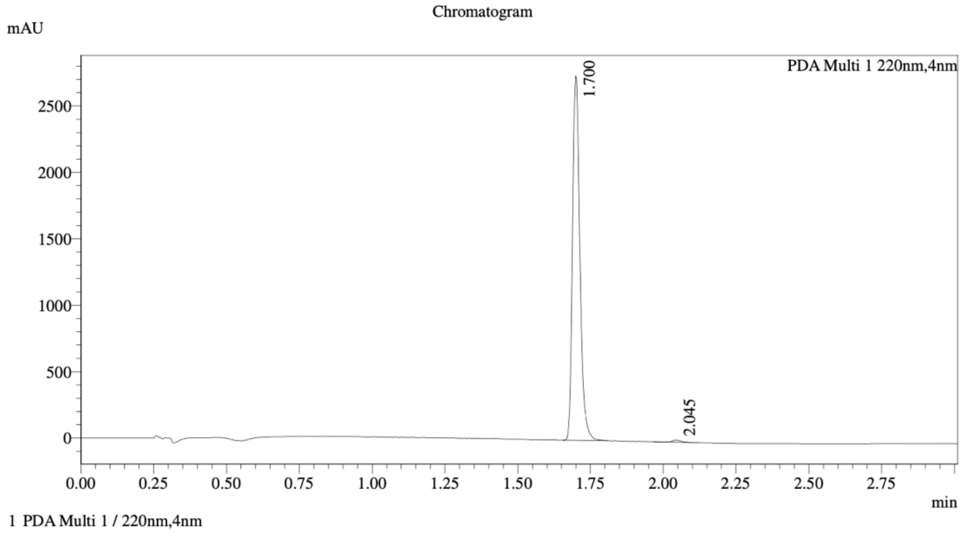


Integration Result

Peak Table

PDA Ch1 220nm

| Peak# | Ret. Time | Height | Height% | Resolution(USP) | Area | Area% |
|---|---|---|---|---|---|---|
| 1 | 1.700 | 2677155 | 99.443 | -- | 4861250 | 99.401 |
| 2 | 2.045 | 15005 | 0.557 | 6.526 | 29310 | 0.599 |

WX-02-921

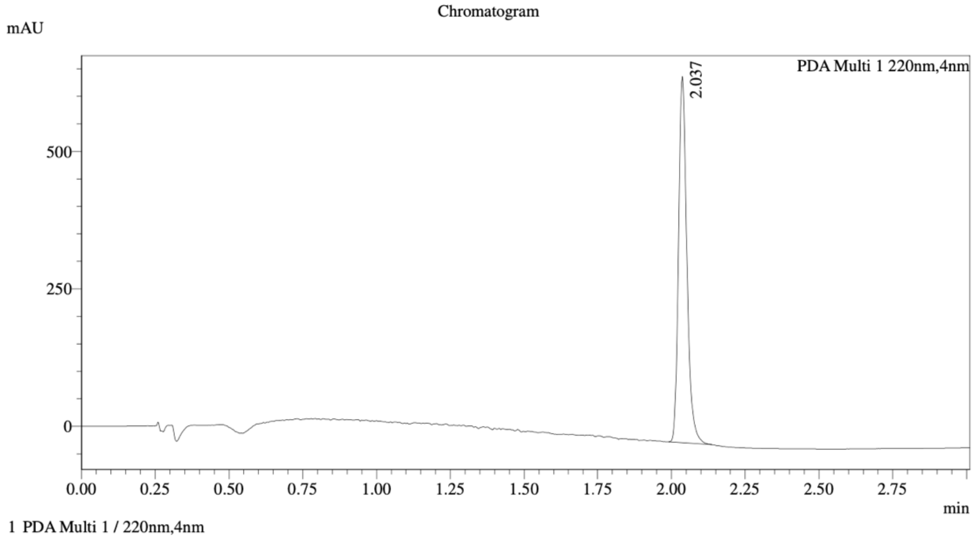


Integration Result

Peak Table

PDA Ch1 220nm

| Peak# | Ret. Time | Height | Height% | Resolution(USP) | Area | Area% |
|---|---|---|---|---|---|---|
| 1 | 2.037 | 655214 | 100.000 | -- | 1307209 | 100.000 |

Mixture of WX-02-922 and WX-02-921

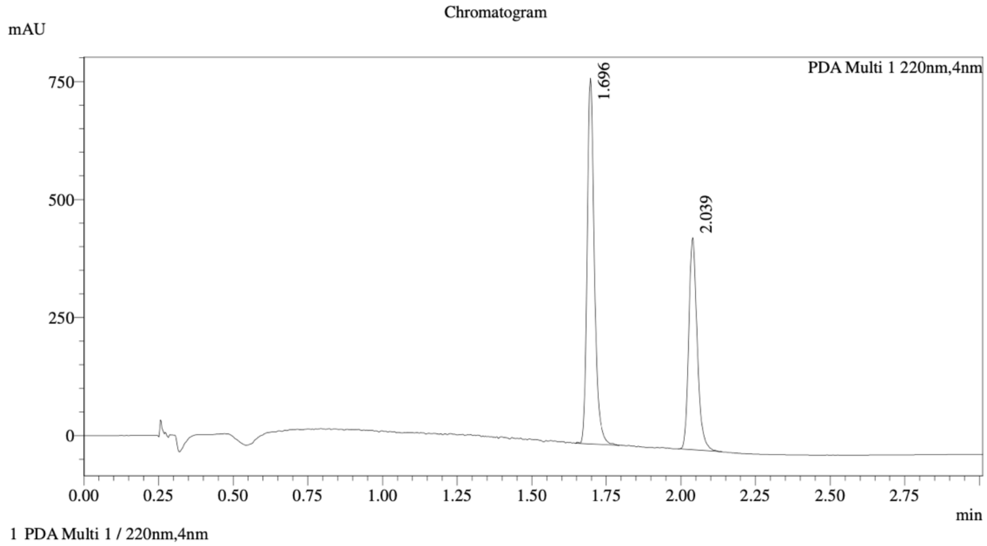


Integration Result

Peak Table

PDA Ch1 220nm

| Peak# | Ret. Time | Height | Height% | Resolution(USP) | Area | Area% |
|---|---|---|---|---|---|---|
| 1 | 1.696 | 727836 | 62.513 | -- | 1303449 | 59.620 |
| 2 | 2.039 | 436451 | 37.487 | 6.714 | 882829 | 40.380 |

Method details

Column: (S,S)Whelk-O1 50×4.6mm I.D., 3.5 µm
Mobile phase: A: $CO_2$, B: 2:1 *i*-PrOH/acetonitrile (0.05% diethylamine)
Elution: 20%-60% B (gradient)

WX-02-924

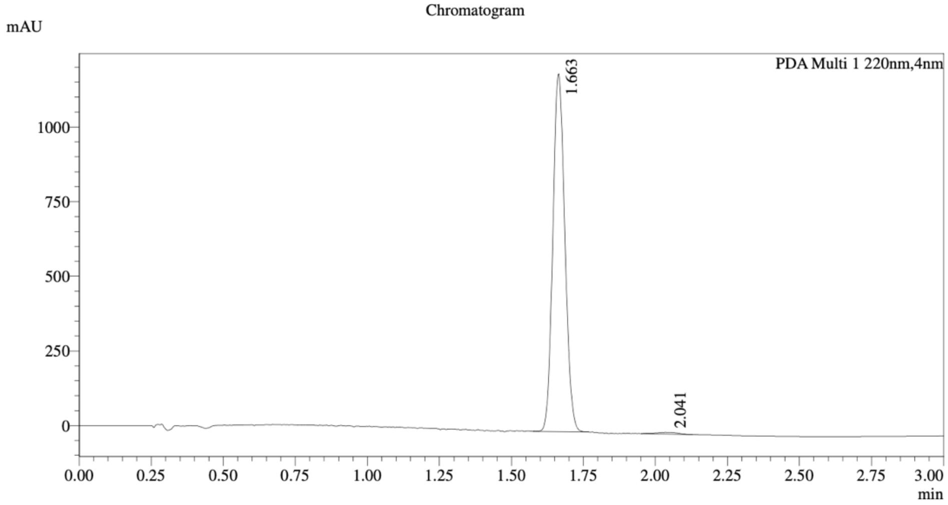


1 PDA Multi 1 / 220nm,4nm

Integration Result

Peak Table

PDA Ch1 220nm

| Peak# | Ret. Time | Height | Height% | Resolution(USP) | Area | Area% |
|---|---|---|---|---|---|---|
| 1 | 1.663 | 1186915 | 99.499 | -- | 3473877 | 99.075 |
| 2 | 2.041 | 5979 | 0.501 | 3.362 | 32445 | 0.925 |

WX-02-923

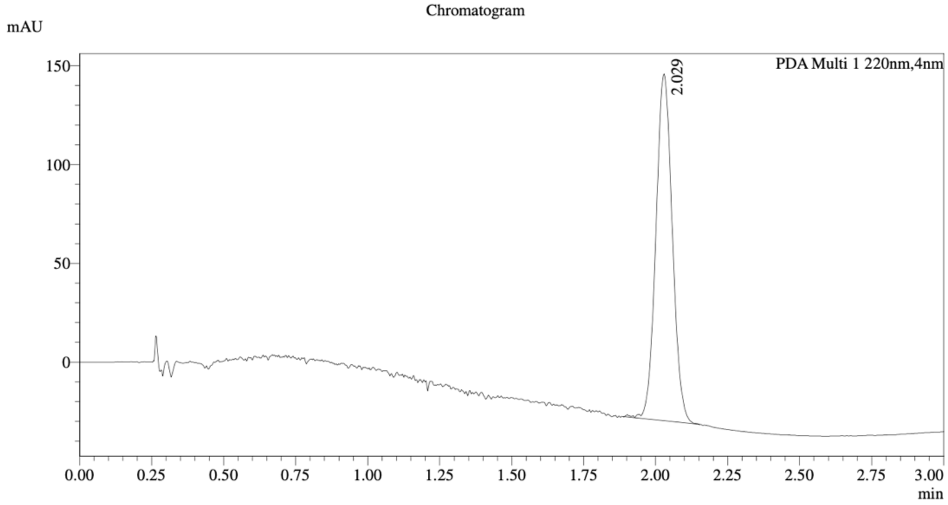


1 PDA Multi 1 / 220nm,4nm

Integration Result

Peak Table

PDA Ch1 220nm

| Peak# | Ret. Time | Height | Height% | Resolution(USP) | Area | Area% |
|---|---|---|---|---|---|---|
| 1 | 2.029 | 174758 | 100.000 | -- | 694690 | 100.000 |

Mixture of WX-02-924 and WX-02-923

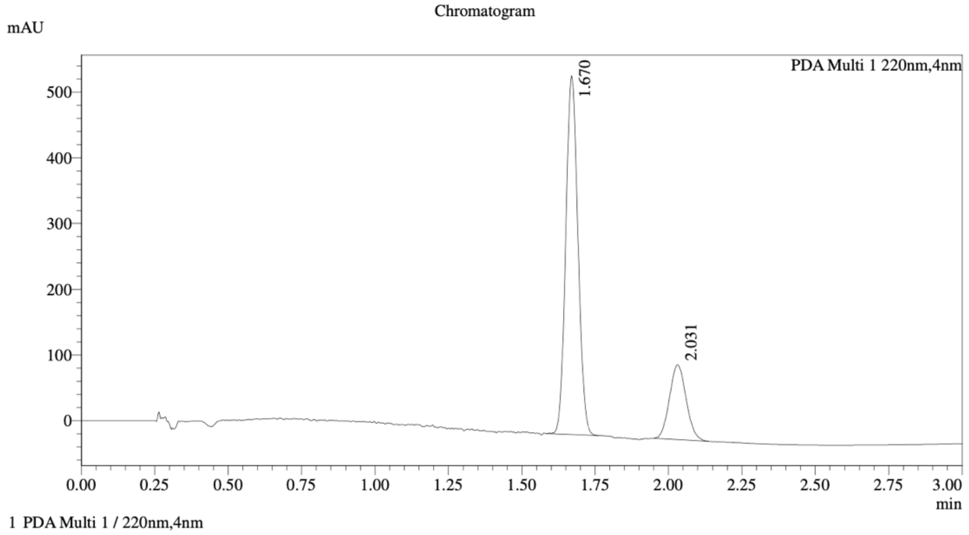


Integration Result

Peak Table

PDA Ch1 220nm

| Peak# | Ret. Time | Height | Height% | Resolution(USP) | Area | Area% |
|---|---|---|---|---|---|---|
| 1 | 1.670 | 542372 | 82.731 | -- | 1576119 | 77.849 |
| 2 | 2.031 | 113213 | 17.269 | 3.887 | 448472 | 22.151 |

Method details

Column: Chiralpak IK-3 50×4.6mm I.D., 3 µm
Mobile phase: A: $CO_2$, B: EtOH (0.05% diethylamine)
Elution: 5%-25% B (gradient)

### References


1. Fulmer, G.R., Miller, A.J.M., Sherden, N.H., Gottlieb, H.E., Nudelman, A., Stoltz, B.M., Bercaw, J.E., and Goldberg, K.I. (2010). NMR Chemical Shifts of Trace Impurities: Common Laboratory Solvents, Organics, and Gases in Deuterated Solvents Relevant to the Organometallic Chemist. Organometallics *29*, 2176-2179. 10.1021/om100106e.
2. Feldman, H.C., Merlini, E., Guijas, C., DeMeester, K.E., Njomen, E., Kozina, E.M., Yokoyama, M., Vinogradova, E., Reardon, H.T., Melillo, B., et al. (2022). Selective inhibitors of SARM1 targeting an allosteric cysteine in the autoregulatory ARM domain. Proc Natl Acad Sci U S A *119*, e2208457119. 10.1073/pnas.2208457119.
3. Njomen, E., Hayward, R.E., DeMeester, K.E., Ogasawara, D., Dix, M.M., Nguyen, T., Ashby, P., Simon, G.M., Schreiber, S.L., Melillo, B., and Cravatt, B.F. (2024). Multi-tiered chemical proteomic maps of tryptoline acrylamide-protein interactions in cancer cells. Nat Chem *16*, 1592-1604. 10.1038/s41557-024-01601-1.
